# Exploring the Evolution of the Cariogenic Oral Pathobiont *Streptococcus mutans* Using Ancient DNA

**DOI:** 10.64898/2025.12.02.691581

**Authors:** Megan Michel, Aida Andrades Valtueña, Murat Akar, Carmen Alonso-Fernández, Micaela Alvarez Calmet, Maria Andreadaki-Vlazaki, Gonzalo Aranda Jiménez, Eszter Bánffy, Rodrigo Barquera, Maria Inés Barreto Romero, Paula Becerra Fuello, Stuart Bedford, Andrey B. Belinskiy, Natalia Berezina, Kamen Boyadzhiev, Yavor Boyadzhiev, Alexandra Buzhilova, Luca Cappuccini, José Enrique Ccencho Huamaní, Eva Celdrán Beltrán, Geoffrey Clark, Mantas Daubaras, Patxuka de Miguel Ibáñez, Marta Díaz-Zorita Bonilla, Miroslav Dobeš, Veit Dresely, Yılmaz Selim Erdal, Michal Ernée, Tiago Ferraz, Levy Figuti, Joan Francès Farré, James Flexner, Susanne Friederich, Dylan Gaffney, Gabriel García Atiénzar, Rafael Garrido Pena, Jorge A. Gómez-Valdés, Maria Giuseppina Gradoli, Marc Guàrdia i Llorens, Evelyn K. Guevara, Svend Hansen, Diana Iraiz Hernandez-Zaragoza, Alina N. Hiss, Tara Ingman, Rimantas Jankauskas, Javier Jiménez Echevarría, Anatoliy R. Kantorovich, Kitti Köhler, Takaronga Kuautonga, Carles Lalueza-Fox, Bastien Llamas, Isabel López-Cortés, Vicente Lull, Rabadan G. Magomedov, Kerttu Majander, Lourdes Marquez, Andrés Martínez Rodríguez, Vladimir E. Maslov, Ken Massy, Harald Meller, Rafael Micó Pérez, Lara Milesi García, Alissa Mittnik, Alan G. Morris, Kathrin Nägele, Elizabeth A. Nelson, Gunnar U. Neumann, Camila Oliart Caravatti, Päivi Onkamo, Anett Osztás, Luka Papac, Cláudia R. Plens, Elisa Pompianu, Juana Ponce García, Cosimo Posth, Eftychia Protopapadaki, Sabine Reinhold, Cristina Rihuete Herrada, Roberto Risch, Manuel A. Rojo Guerra, Alejandro Romero, Antti Sajantila, Domingo C. Salazar-García, Tasneem Salie, Margarita Sánchez Romero, Duncan Sayer, Stephan Schiffels, Torsten Schunke, Arda Sevkar, Rula Shafiq, Richard Shing, Robin Skeates, Eirini Skourtanioti, Alessandra Sperduti, Maria A. Spyrou, Philipp W. Stockhammer, André Strauss, Glenn Summerhayes, Anna Szécsényi-Nagy, Cristina Tejedor Rodriguez, Frédérique Valentin, Guido Valverde, Andrejs Vasks, Carlos Velasco Felipe, Miriam Vílchez Suárez, Vanessa Villalba-Mouco, Edson Willie, Kutlu Aslihan Yener, Gunita Zariņa, Wanda Zinger, Christina Warinner, Wolfgang Haak, Alexander Herbig, Johannes Krause

## Abstract

The oral pathobiont *Streptococcus mutans* can contribute to dental caries development through metabolism of dietary carbohydrates. Adoption of carbohydrate-rich agricultural diets is associated with increased prevalence of dental caries in archaeological populations; however, the evolutionary impact of changing subsistence strategies on cariogenic microbes like *S. mutans* remains to be explored. Here, we use a novel hybridization capture reagent to generate genome-wide ancient DNA data from a global set of 75 *S. mutans* strains spanning the last 8,000 years. Most virulence-associated genes predate the origins of agriculture; however, we highlight loci regulating genetic competence, bacteriocin production, and biofilm formation which are absent in 5 strains from pre-agricultural ancient hunter-gatherers, suggesting that their acquisition may have been associated with adaptation to carbohydrate-rich agricultural diets. Together, our study highlights ancient DNA as a promising tool for exploring the dynamic interplay between subsistence strategy, microbes, and dental pathology in human populations through time.

## INTRODUCTION

Dental caries, also known as cavities, are a nearly ubiquitous affliction of modern industrialized populations, with over 2.3 billion people suffering from caries in permanent teeth each year^1^. Caries can also cause significant morbidity and mortality in populations lacking access to antibiotics; for example, the weekly London Bills of Mortality published throughout the 17th and 18th centuries consistently list “teeth”, likely referring to dental infections beginning as caries, as a leading cause of death^2^. Caries result from oral microbial metabolism of fermentable carbohydrates, producing organic acids that demineralize tooth enamel and/or dentine^3^. Historically, caries have been primarily attributed to infection with particular microbial species, including the gram-positive oral microbe *Streptococcus mutans*^3^. However, caries are increasingly recognized as a multifactorial disease^4,5^; the ecological plaque hypothesis posits that dental caries result when diverse biological, environmental, behavioral, and cultural factors, including diet, the oral microbiota, dental hygiene practices, and systemic health shift the oral plaque microenvironment towards a state of dysbiosis^5,6^.

Beyond their modern impact, caries posed a significant health burden for past populations. Dental caries have affected modern humans since the Pleistocene^7^ and have also been identified in ancient hominins and wild primates, suggesting a long history of dental disease in the hominid lineage (e.g.^8–13)^. However, the burden of caries has increased significantly in recent evolutionary time. Development of new subsistence strategies, including plant-based agriculture, animal husbandry, and dairying have radically altered diets over the past millennia, exerting diverse selection pressures on both human populations and our resident microbes^14,15^.

Multiple studies report higher caries rates in agricultural populations compared to hunter-gatherer groups, which is generally attributed to an increased reliance on staple crops rich in fermentable carbohydrates^14^. Similarly, the rise of industrialization led to adoption of modern diets rich in highly processed, sugar-rich foods. The enormous burden of dental caries today has been described as an example of an evolutionary mismatch, in which pathology arises from consumption of diets radically different from those with which our bodies evolved^16^.

While changing subsistence strategies likely impacted caries prevalence over the past 10,000 years, the role of the oral microbiota in the shifting prevalence of dental pathologies remains unclear. Given that the oral cavity functions as a gateway for nutrient intake and supports the second-most diverse microbial community in the human body, one might expect that shifting subsistence strategies would drive concomitant adaptation in the oral microbiota^17^. In fact, analyses of oral communities in modern populations find a limited impact of diet on plaque microbial community composition^18^, perhaps because oral microbes metabolize nutrients in saliva and gingival crevicular fluid in addition to ingested foodstuffs^19^. Similarly, profiles of ancient oral microbiota preserved in archaeological dental calculus reveal limited correlation between subsistence strategy and community composition. Instead, many microbes are core members of the oral microbiota, meaning that they are shared by modern humans, including agriculturalists and hunter-gatherers, Neanderthals, African great apes, and even more distantly-related primate taxa^20,21^.

Despite stability in oral microbial community composition through time, the role of particular cariogenic taxa in the past prevalence of dental cavities remains to be explored. Once regarded as the major etiological agent of dental caries, the gram-positive oral pathobiont *S. mutans* exhibits multiple virulence phenotypes that contribute to its high cariogenic potential, particularly in the presence of fermentable carbohydrates^3^. For example, glucosyltransferases produced and secreted by *S. mutans* convert sucrose into extracellular polymeric substances (EPS), major structural components of oral biofilms^22^. *S. mutans* also expresses numerous cell surface proteins that facilitate adhesion to extracellular polysaccharides and the acquired enamel pellicle, facilitating the formation of cariogenic plaque^23^. Furthermore, *S. mutans* strains metabolize a diverse array of both simple sugars and complex polysaccharides such as starch, producing lactic acid that contributes to enamel demineralization^3,24^. In addition to its acidogenicity, *S. mutans* is acid-tolerant (aciduric), growing and outcompeting other oral taxa in low pH microenvironments^24^. Quorum sensing pathways enable *S. mutans* to respond to local environmental conditions, while peptide antibiotics known as bacteriocins facilitate interspecific competition in dynamic biofilm communities^24,25^. Finally, *S. mutans* exhibits competence, the capacity to take up and incorporate extracellular DNA into its own genome; consequently, strains display significant intraspecific genomic diversity, which may facilitate survival and adaptation to a range of environmental conditions^24,26,27^.

Despite widespread clinical interest in *S. mutans*, the bacterium’s role in Neolithic-associated changes in dental pathology remains unclear. Demographic models based on modern genomes infer exponential growth of *S. mutans* populations c. 10,000-5,000 years ago, linking the species’ proliferation with the adoption of agriculture in Eurasia and the Americas^26,28^. The assembly of the first ancient *S. mutans* genome from a Bronze Age individual from Killuragh, County Limerick, Ireland (KGH2-B) demonstrates the presence of *S. mutans* in European populations as early as c. 4,000 years ago^29^. In contrast, PCR-based analyses of ancient DNA preserved in dental calculus suggest an absence of *S. mutans* in European Mesolithic and Neolithic contexts, instead linking the bacterium’s spread with the recent adoption of modern, industrialized diets^30^. Phylogenetic analyses incorporating the ancient KGH2-B *S. mutans* genome support the hypothesis of recent *S. mutans* population expansion, possibly linked to industrial-era dietary shifts^29^.

Beyond the bacterium’s antiquity in human populations, questions persist regarding when and how *S. mutans* acquired specific virulence phenotypes and whether these may have contributed to past changes in dental pathology. Each *S. mutans* genome contains roughly 2,000 genes, which can be subdivided into core and accessory genome components. The core genome includes the common set of roughly 1,000 to 1,500 orthologous genes shared by nearly all members of the species. The remainder of each strain’s genetic repertoire belongs to the accessory genome, genes present in some but not all *S. mutans* strains^26,27^. Together, the core and accessory gene content observed across strains comprises the *S. mutans* pan-genome. While the core genome supports fundamental metabolic processes, strains harbor enormous variation in accessory gene content. In *S. mutans*, the accessory genome includes a significant proportion of virulence-associated loci, making a pan-genomic approach critical for exploring the evolution of cariogenicity^26,27^. However, perhaps due to the dynamic nature of the genome, with frequent recombination and intra-and interspecific exchange of genetic material, modern *S. mutans* populations exhibit limited phylogeographic signal^26^; uncertainty regarding patterns of strain relatedness complicates attempts to infer patterns of gene acquisition and loss using modern genomes alone.

Recent advances in the field of ancient DNA have made it possible to generate genome-wide data from a variety of pathogenic bacteria, viruses, and protozoa preserved in skeletal material^31,32^. Several studies have succeeded in detecting *S. mutans* using metagenomic techniques^30,33^, and the recent publication of an ancient *S. mutans* genome assembly represents a critical step towards understanding the history of this cariogenic taxon^29^. Nevertheless, previous work has been hampered by small sample sizes, particularly from early Holocene contexts, leading to uncertainty regarding this species’ antiquity in human populations. Furthermore, the availability of only a single ancient *S. mutans* genome-wide dataset has precluded analyses of variation in virulence-associated gene content through time. To address these gaps, we used a custom hybridization capture array to generate ancient *S. mutans* genome-wide data from a global set of 75 strains spanning the Mesolithic to the Modern Era (c. 8000 to 150 years before present). To our knowledge, this dataset provides the first evidence for the presence of *S. mutans* in hunter-gatherer-fisher populations from Eurasia, Africa, and the Americas. While we observe limited phylogeographic signal in our ancient dataset, we document the presence of *S. mutans* across c. 8,000 years of bacterial evolution. Finally, using an ancient pan-genomic approach, we explore spatial and temporal variation in virulence gene content, shedding light on the evolutionary history of this cariogenic oral pathobiont in ancient human populations.

### Ancient *S. mutans* Data Generation

To identify ancient individuals exhibiting evidence of *S. mutans* DNA preservation, we screened previously-produced shotgun DNA sequencing datasets obtained from teeth and bones using the metagenomic analysis pipeline HOPS (Heuristic Operations for Pathogen Screening) (**Methods**)^34^. Prior studies have shown that because oral microbes contribute to postmortem tissue decomposition, they can be recovered from approximately 15% of skeletal remains, thus preserving information about ancient salivary and other oral bacteria^35^. Libraries from 75 candidate *S. mutans*-positive individuals were enriched using a custom in-solution hybridization capture reagent targeting the full genomic diversity present in 191 modern *S. mutans* assemblies (**Methods; Figure 1; Supplementary Table 1**). Assessment of characteristic patterns of ancient DNA damage supports the authenticity of our ancient genomes; as observed for other pathogenic microbes^36^, we document correlation between 5’ C to T substitution rates on the first base pair of sequencing reads aligning to human and *S. mutans* genomes, respectively (**Figure 2**). After mapping to the *S. mutans* UA159 reference genome, 63 out of 75 libraries yielded a median genomic coverage greater than or equal to 1.0x, with median coverage depths ranging from 1.0 - 83x (**Supplementary Table 2)**. Together, this dataset demonstrates the presence of *S. mutans* in ancient individuals spanning the Mesolithic to the early Modern period and deriving from the territory of 22 modern countries on six continents and the Pacific Islands. Notably, 35 of these genomes attained a median coverage of the UA159 reference greater than or equal to 10x, facilitating analyses requiring higher genomic coverage. Based on the frequency of multiallelic SNPs in these higher coverage datasets, we infer that 19 of 35 (54%) *S. mutans* genomes likely derive from single-strain infections, while sixteen genomes (46%) appear to represent mixtures of multiple *S. mutans* strains and/or closely related species, either in symmetric or asymmetric proportions (**Supplementary Table 2; Supplementary Figure 1**).

**Figure 1.**
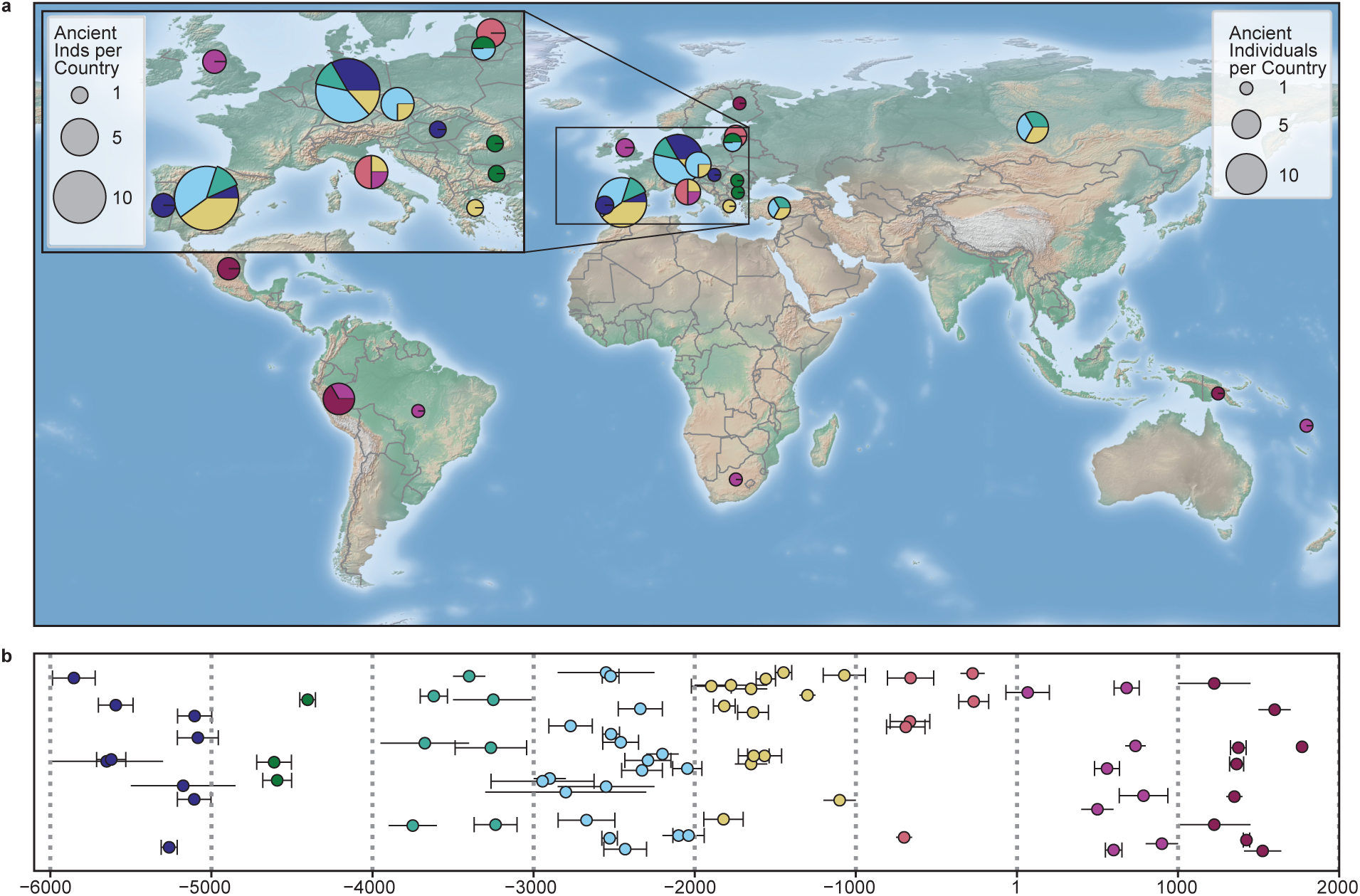
Spatial and temporal distribution of 75 ancient individuals yielding *S. mutans* genome-wide data. **a.** *S. mutans*-positive ancient individuals included in this study summarized by modern-day country. Pie-point size is proportional to the number of ancient *S. mutans* genomes per country, while color reflects the millennial age estimation for each ancient individual (bottom panel). Map inset shows ancient *S. mutans* strains derived from Europe. Pie points were plotted using previously-published centroids of modern countries (https://github.com/gavinr/world-countries-centroids/releases). **b.** Timeline depicting the temporal span of ancient *S. mutans*-positive individuals. Circles mark the midpoint of sample age estimates. Capped error bars indicate 95% confidence intervals for radiocarbon-dated samples, while uncapped error bars depict date ranges estimated based on archaeological context.

**Figure 2.**
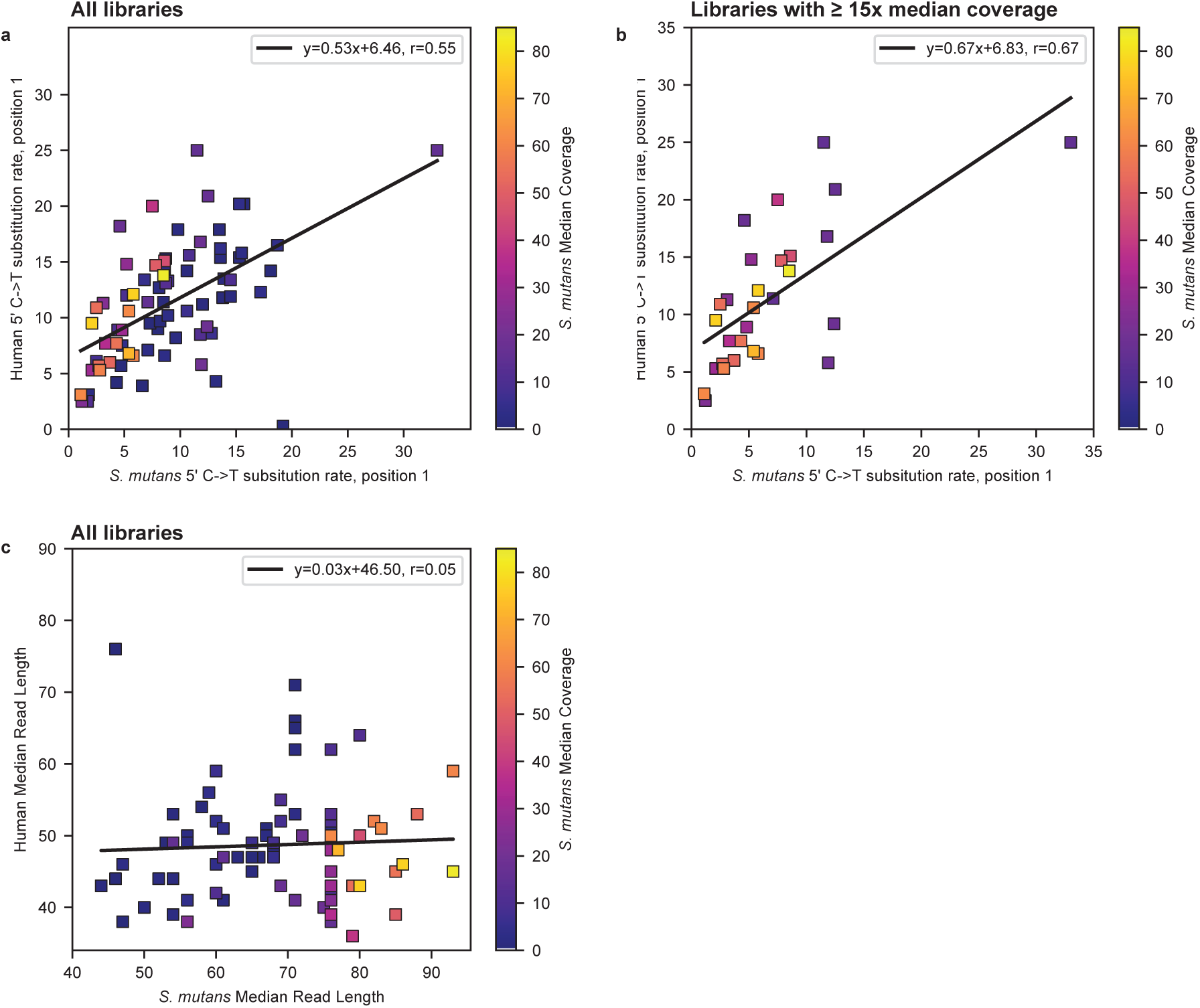
Ancient DNA damage and fragmentation **a.** Following hybridization capture, libraries exhibit a positive correlation between 5’ C to T substitution rates at the first position of reads mapping to the *S. mutans* UA159 and human (HG19) reference genomes. All 75 ancient libraries yielding *S. mutans* genome-wide data are shown. **b.** Correlation between 5’ C to T substitution rates on the first base pair of reads mapping to the *S. mutans* UA159 and human (HG19) reference genomes. Only libraries yielding a median coverage of the UA159 reference greater than or equal to 15x are shown. **c.** Libraries exhibit limited correlation between median lengths of reads mapping to the *S. mutans* UA159 and human (HG19) references, respectively. In panels a-c, points are colored according to the median coverage of the UA159 reference genome. Linear regressions were computed using the SciPy (v. 1.9.0) module stats^98^.

### *S. mutans* Spread Predates Transition to Agriculture

As evidenced by both ancient metagenomic analyses^33,37,38^ and genome assembly^29^, *S. mutans* affected European populations by the Bronze Age; however, to our knowledge, the impact of this cariogenic microbe on ancient pre-agricultural populations remains to be substantiated using paleogenomic data. Similarly, although PCR-based studies point to the presence of *S. mutans* in the Americas before and after European contact^39^, genome-wide datasets from outside of Europe are currently lacking. Here, we report the first ancient *S. mutans* genome-wide data from six pre-agricultural hunter-gatherers and/or hunter-gatherer-fishers from Eurasia, Africa, and the Americas. Deriving from three continents and spanning thousands of years of human history, these individuals provide robust evidence for *S. mutans* infection in pre-agricultural populations reliant on heterogeneous dietary sources.

Of the newly-reported *S. mutans* genome-wide datasets, the oldest derive from a pair of Mesolithic hunter-gatherer individuals from the Muge region of Portugal. CMS002 (c. 6000-5300 BCE) from Concheiro da Moita do Sebastao yielded low-coverage *S. mutans* genome-wide data, while CCA002 (5986-5721 cal BCE^40^) from Concheiro do Cabeço da Arruda produced a median coverage of the *S. mutans* UA159 reference genome of 2.0x (**Supplementary Table 2**). Both sites form part of a larger, regional series of late Mesolithic shell middens, and dietary isotope analysis suggests individuals from Concheiro da Moita do Sebastao and Concheiro do Cabeço da Arruda likely consumed a mixed marine/terrestrial diet^40^. Interestingly, paleopathological assessment of c. 100 burials from the Mesolithic site of Concheiro da Moita do Sebastao noted a high proportion of dental caries, possibly linked to consumption of cariogenic fruits by the local population^40–42^.

Two additional pre-Neolithic European *S. mutans* genomes were recovered from sites in modern-day Germany and Lithuania. A genetically female Mesolithic individual (BOT005) from the site of Bottendorf in modern-day Sachsen-Anhalt yielded an *S. mutans* genome with a median coverage of 6.0x. Radiocarbon-dated to 5636-5572 cal BCE, BOT005 harbors exclusively Western hunter-gatherer (WHG)-related ancestry. Evidence from dietary isotopes suggests that the inhabitants of Bottendorf practiced an opportunistic subsistence strategy, likely consuming both terrestrial game and limited quantities of freshwater resources^43,44^. Finally, a 35-40-year-old genetically female individual (DON004) from the central north-western Lithuanian site of Donkalnis yielded a high-coverage *S. mutans* genome (77x median coverage)^45^. While use of the settlement and associated cemetery spanned the Mesolithic to the Middle Bronze Age, DON004 is radiocarbon dated to 4720-4530 calBCE and associated with the pottery producing Mesolithic Narva culture^45^. Isotope data suggests that individuals from Donkalnis and other inland Subneolithic sites relied on consumption of freshwater fish and forest game with the likely addition of wild plant material^46^.

Finally, in addition to these European Mesolithic and Subneolithic cases, we recovered *S. mutans* genomes from one hunter-gatherer-fisher from Brazil’s Atlantic Coast and one Later Stone Age South African hunter-gatherer. Excavated from the site of Pavão 16 in Sambaqui on the Atlantic Coast of modern-day Brazil, individual PVA001 yielded an *S. mutans* genome with 3.0x median coverage. Radiocarbon dated to 482-637 cal CE, PVA001 was buried in a small, rounded shell mound containing the remains of two other individuals^47^. Lastly, located in the Western Cape Province of South Africa, the Faraoskop rock shelter contained the remains of 12 individuals and associated Later Stone Age artifacts^48–50^. Dated to 67 cal BCE-202 cal CE, the 40-50-year-old male individual FAR004 yielded an *S. mutans* genome with a median coverage of 23x after hybridization capture^48,50^. Interestingly, FAR004 suffered from carious lesions, as did 3 of the other 4 Faraoskop individuals for whom dentition was recovered. Dietary isotopic evidence suggests a largely terrestrial diet, and researchers posit that the high prevalence of caries in the Faraoskop individuals may be due to consumption of starch rich tubers^49^.

### Limited Phylogeographic Signal in Ancient Datasets

For many ancient bacteria and viruses, phylogenetic and population genetic analyses provide a unique glimpse into patterns of historical population structure, shedding light on the timing and routes of global dissemination for pathogenic taxa^32^. To explore the genetic affinities of our ancient *S. mutans* strains, we analyzed the newly-reported genomes alongside shotgun sequencing data from one previously-published ancient *S. mutans*-positive individual (KGH2-B) and 456 publicly available, modern *S. mutans* assemblies downloaded from the NCBI Refseq database (**Methods; Supplementary Table 3**). We computed a maximum-likelihood phylogeny including the 35 newly-published ancient *S. mutans* genomes with a median genomic coverage of at least 10x, using an alignment of 40,679 biallelic segregating core-genome SNPs present in all strains (n = 1000 bootstrap replicates; **Figure 3**). Consistent with previous studies, we find that both modern and ancient *S. mutans* strains exhibit limited phylogeographic structure, even at a continental scale. Furthermore, as indicated by the prevalence of low bootstrap support values in our maximum-likelihood tree, many clades within the *S. mutans* phylogeny are poorly resolved (**Figure 3)**. This may be due to the high prevalence of recombination throughout the *S. mutans* genome, as horizontal transfer of genetic material violates the assumptions of standard phylogenetic approaches modeling clonal evolution^26^.

**Figure 3.**
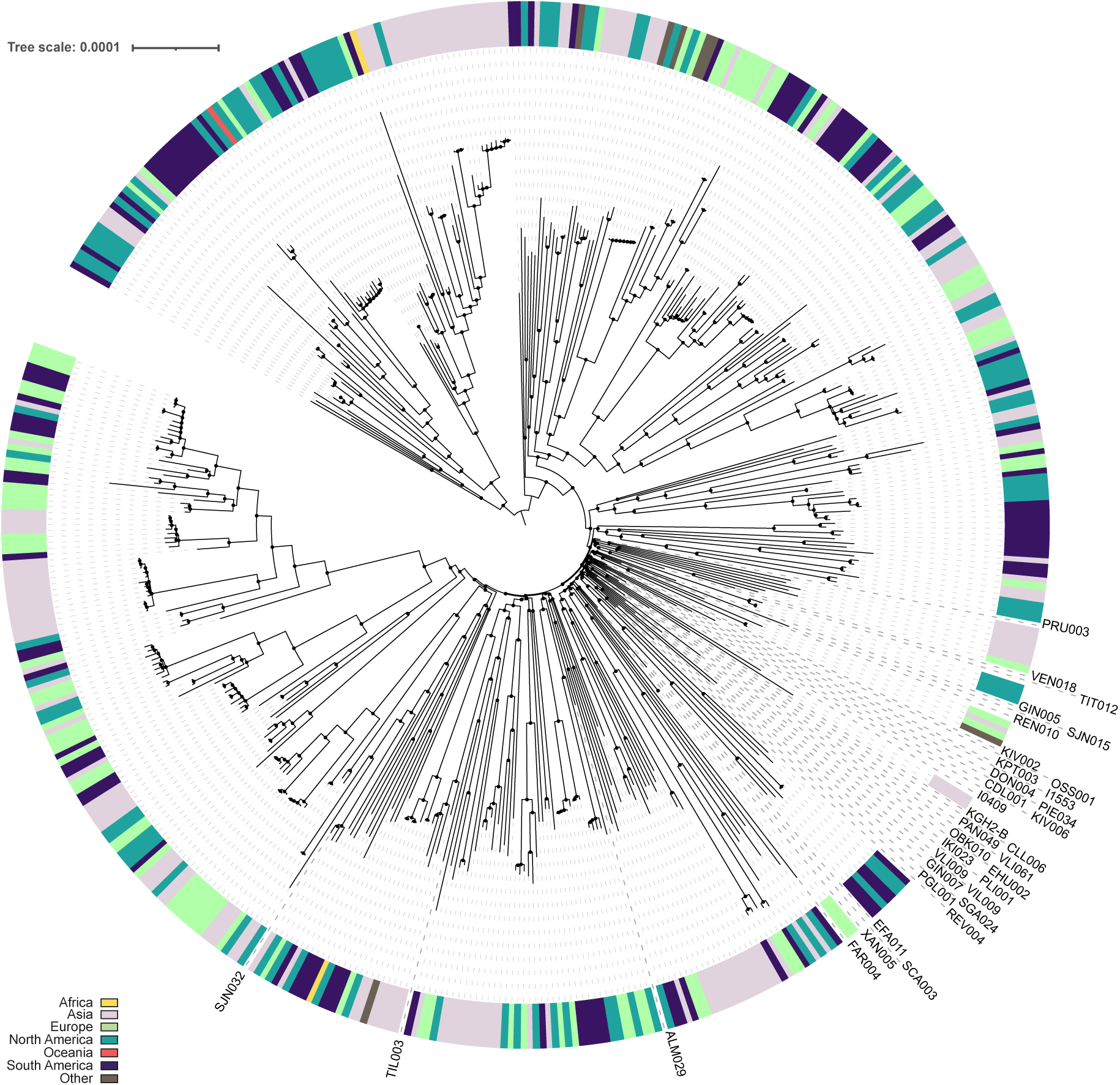
Phylogeny of modern and ancient *S. mutans* strains Midpoint-rooted maximum-likelihood phylogeny (RAxML-NG, v. 1.1) including 456 modern *S. mutans* assemblies, the previously-published ancient KGH2-B dataset, and 35 newly-reported ancient *S. mutans* genomes exceeding 10x median coverage. Modern strains are colored according to continent/region of origin, and ancient genomes are labelled with individual names. Black points mark nodes receiving at least 90% bootstrap support (1,000 bootstrap replicates).

To account for the potential impact of recombination on population structure, we constructed a principal components analysis using 118,675 high-quality, biallelic core SNPs genotyped in greater than or equal to 95% of the 456 modern *S. mutans* strains. Again, we observe limited phylogeographic substructure; the modern *S. mutans* assemblies form three clines, each of which is composed of mixed strains deriving from multiple continents (**Methods; Supplementary Figure 2**). When projected onto these axes of variation, both the previously-published KGH2-B genome and newly-reported ancient *S. mutans* genomes with greater than or equal to 3x median genomic coverage fall close to the intersection of the three clines, partially overlapping a gap in the modern genomic diversity. Within the ancient cluster, we observe that strains from different temporal and geographical contexts cluster together, even after exclusion of individuals with mixed-strain *S. mutans* infections (not shown). While this finding is consistent with the results of other studies, the causes underlying the apparent lack of phylogeographic signal in both ancient and modern *S. mutans* strains remain to be elucidated (**Discussion)**^26^.

### *S. mutans* Pan Genome

The majority of paleogenomic studies to date rely on mapping to a modern reference genome to reconstruct genetic variation among ancient strains. However, as *S. mutans* exhibits striking variation in accessory gene content, a strategy reliant solely on reference-based mapping may fail to recover variation in evolutionarily relevant and/or virulence associated accessory gene clusters. Moreover, use of a modern reference precludes the possibility of recovering gene clusters that are unique to ancient strains. Therefore, taking advantage of the high coverage after capture, we *de novo* assembled all ancient datasets with a minimum coverage of 10x (35/75) together with the previously reported shotgun *S. mutans* genome KGH2-B (ref. ^29^, Methods). We obtained 36 *S. mutans* Metagenomic Assembled Genomes (MAGs) with a mean completeness of 94.15%±7.24 (range 63.55-100%) and mean contamination of 0.52%±0.54 (range 0-2.74%) as reported by CheckM^51^ (**Supplementary Table 4**). Next, we constructed a pan-genome incorporating the diversity present in both the ancient MAGs and 456 publicly available modern *S. mutans* assemblies (**Methods**). Analysis of this hybrid ancient/modern *S. mutans* genome provides an unprecedented opportunity to explore variation in *S. mutans* core and accessory gene content as well as pan-genome size over a period of roughly eight millenia.

Consistent with the results of previous studies, we find that ancient and modern *S. mutans* strains harbor a stable core-genome and a large pan-genome with significant variation in accessory genome content. The total pan-genome size estimated for *S. mutans* contained 10,651 genes, of which 1,076 gene clusters are shared by more than 95% of strains (core genes, **Figure 4a-b**); this finding is broadly consistent with the results of previous studies, which inferred core-genome sizes of 1,083 and 1,490 genes based on analyses of 183 and 56 assemblies, respectively^26,27^. The accessory genome, defined here as the set of orthologous gene clusters present in less than 95% of *S. mutans* lineages, accounts for the majority of gene clusters reconstructed (9,575/10,651, 89.9%). Most accessory gene clusters are rare and/or unique (**Figure 4c**), meaning that sampling of additional *S. mutans* strains is likely to yield additional genomic diversity, which is typical of species with an open pan-genome.

**Figure 4.**
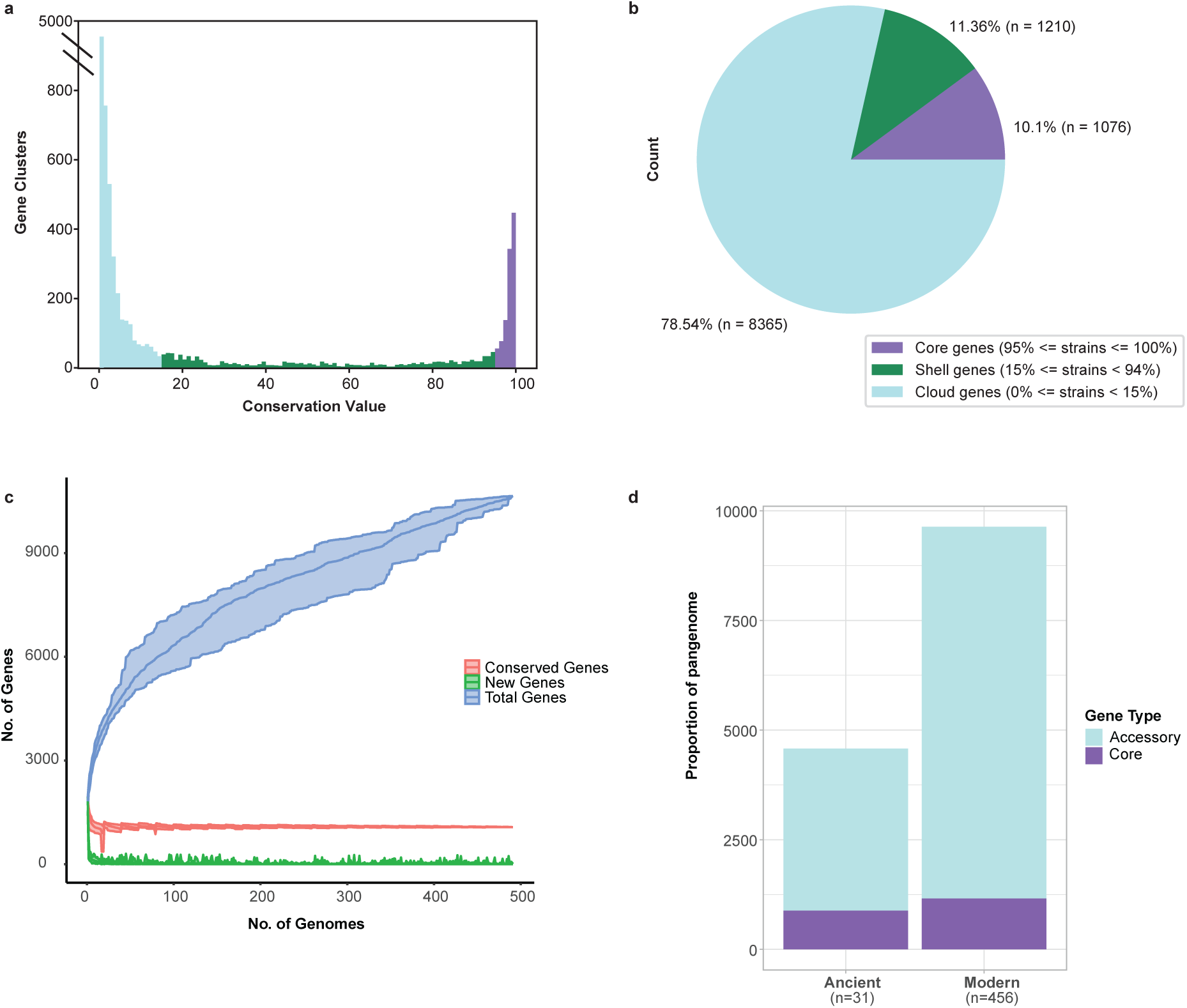
*S. mutans* pan-genome reconstruction. **a.** Histogram depicting the percentage of modern *S. mutans* assemblies in which gene clusters from our reconstructed pan-genome are present. Bin coloration reflects genes categories, following the legend in panel b. **b.** Pie chart showing the number and percentage of reconstructed gene clusters defined as core, soft core, shell and cloud. **c.** Rarefaction curves depicting how counts of conserved genes, new genes, and total genes change with sampling of additional modern *S. mutans* genomes. **d.** Gene counts of core genes and accessory genes in the modern and ancient assemblies.

To further explore the gene content of the ancient assemblies and compare it to the modern genomes, we subdivided the hybrid ancient/modern pan-genome into ancient vs modern pan-genome by redefining the core and accessory on the ancient MAGs with a minimum completeness of 90% subset versus the modern assemblies subset. A gene will be considered core if present in a minimum of 95% of the subset and accessory if it does not fulfill this requirement. The pan-genome size for the modern assemblies is bigger than in the ancient MAGs (**Figure 4d**), while the proportion of core genes appears to be higher in ancient assemblies when compared to the modern ones. These observations are likely due to the difference in sample size for these two groups (31 ancient vs 456 modern strains) and *S. mutans* having an open pan-genome (see above). The probes used to generate the data for the reconstructed MAGs, with exception of KGH2-B, were based on only a subset of the modern *S. mutans* genomes (191/457, **Methods**). To account for the potential loss of diversity in our MAGs due to biases introduced during the capture enrichment experiment, we mapped the probes to the reconstructed hybrid ancient/modern pan-genome. We found that the capture probes spanned at least 90% of the sequence length for 8,382 of 10,651 (78.7%) gene clusters. Of the genes with less than 90% of the sequence length covered by the probes, we observe the presence of 739(32.57%) of these genes in our ancient MAGs. The remaining genes not covered by the probes are part of the accessory genome component of *S. mutans* and present on average in 6 genomes (sd=25.91).

Overall this findings suggests a robust capacity to evaluate the presence of common loci in our ancient capture datasets. We then compared the core genes identified in the hybrid pan-genome, to those identified as core in the ancient subset. Of the 1,062 core genes in the hybrid pan-genome, 807 genes are also categorized as ancient core (75.28%). The frequency of the remaining 265 core gene clusters in the ancient *S. mutans* MAGs ranges from 45-90% (mean= 78.99, sd=11.11). We also identified 74 genes that are core in the ancient (**Supplementary Table 5**). Those genes are part of the accessory pan-genome in the modern *S. mutans* strains (mean=331.11, sd=112.84).

To explore whether the difference between ancient and modern pan-genome sizes is consistent with a difference in genome size, we compared the number of genes in the ancient MAGs and modern assemblies. We observe a smaller mean genome size in the assembled ancient genomes when compared to the modern genomes (**Supplementary Figure 3**). To account for potential gene loss during *de novo* assembly, MALT filtered reads for the ancient genomes were mapped to the pan genome reference, and we calculated the genome size based on the genes identified by mapping (Ancient Mapping, see **Methods**) and with the union between the genes recovered from both mapping and assembly (Ancient Merged, see **Methods**). In both cases, we observe a much larger genome size than that of the assembled ancient genomes as well as the modern assemblies. This can be explained either by genes present in regions of the genome that could not be assembled, or by missmapping of other closely related organisms.

When we compare the difference in the genes identified only by mapping (n=2,831), we observe that with the exception of 6 core genes, the rest are accessory genes. The vast majority of the accessory genes are considered cloud accessory genes (n=2,672, genes present in less than 15% of the strains) and only a minority are shell accessory genes (n=153, present in 15-95% of strains).

We also checked whether the number of genes present in the assemblies could be explained by their mean coverage of the UA159 reference, the dating of the individual, or the MAG completeness. We see only a correlation with the latter (**Supplementary Figure 4**). It is also noteworthy that the KGH-2B genome, the only shotgun genome included in our assemblies, displays the highest number of genes of all assembled MAGs. Given the potential missmapping observed in the dataset, this could have an effect on the quality of the assembly, where regions of the genome prone to missmapping would be more difficult to assemble and result in chimeric sequences that would be discarded during the contig binning step. This is an avenue for future research. Nevertheless, our results highlight the importance of performing *de novo* assembly regardless of the mean coverage.

### Subsistence Strategy and *S. mutans* Virulence Capacity

Finally, in order to explore the acquisition of virulence-associated phenotypes in ancient *S. mutans* strains, we focused on a subset of 93 previously-identified virulence-associated loci linked to phenotypes such as acidogenicity and/or acidurity, biofilm formation, competence development, intrabiofilm competition, nutrient acquisition and reservation, and cell signaling (**Supplementary Table 6**)^27^. To investigate the antiquity of these loci, we tested the proportion of each gene cluster covered in a set of 50 ancient *S. mutans* strains with a median depth of coverage of the UA159 reference genome greater than or equal to 3x (**Figure 5**). Notably, many of the virulence-associated loci are present in strains up to c. 8,000 years old, suggesting that they are evolutionarily ancient. Loci linked to cariogenicity are also present in ancient human populations practicing diverse subsistence strategies, both before and after the introduction of agriculture; however, we note that three genes, *comC*, *comD*, and *comE*, are missing in 3 of 4 high-coverage hunter-gatherers in our dataset (**Figure 5**). The remaining one high-coverage hunter-gatherer, the 2000-year-old ancient South African individual FAR004, harbors a mixed strain infection; based on the relative coverage of surrounding genomic regions, we hypothesize that *comCDE* are likely absent in one of the two strains present (**Figure 5**).

**Figure 5.**
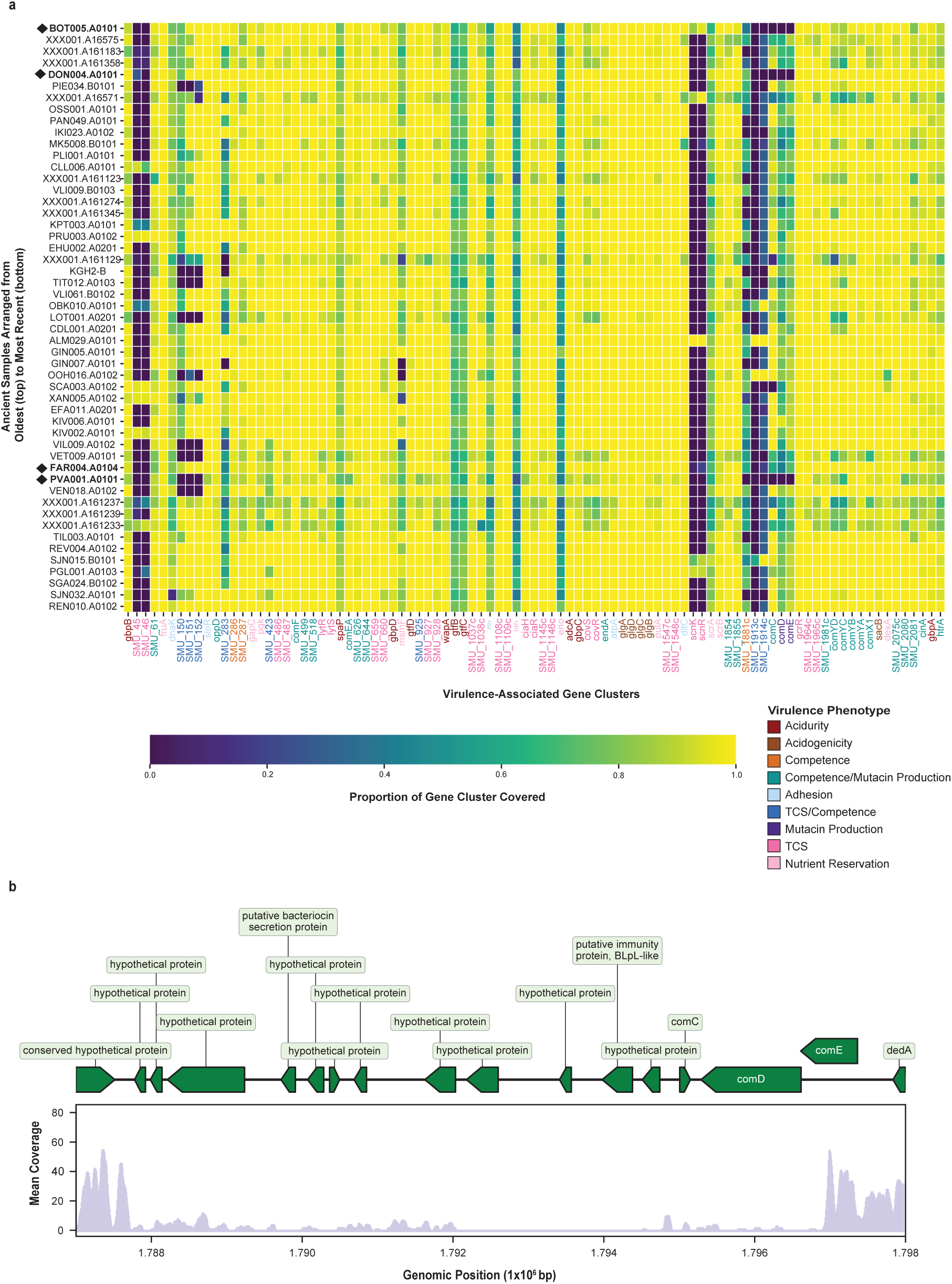
Ancient *S. mutans* virulence gene content (next page) **a.** Virulence-associated gene content in ancient *S. mutans* strains with greater than or equal to 3x median coverage of the UA159 reference genome. Coloration reflects the proportion of gene cluster length covered to a depth of at least 1x. Ancient strains are ordered from oldest (top) to most recent (bottom), and n = 4 strains isolated from hunter-gatherers and/or hunter-gatherer-fishers are marked with a diamond symbol. Gene names are colored according to the virulence phenotype(s) with which they are associated, following ref. ^27^. **b.** Genomic coverage of the comCDE region in the *S. mutans* strain(s) isolated from the South African hunter-gatherer FAR004. Coding sequences are shown in green, and the bottom panel depicts mean genomic coverage computed in adjacent 10 bp windows.

ComC, ComD, and ComE are key upstream regulators of genetic competence, the physiological state in which *S. mutans* can take up and incorporate DNA from the extracellular environment into its own genome. In *S. mutans*, competence is regulated via a complex signaling cascade triggered by environmental signals; in addition to regulating the bacterium’s capacity for genetic transformation, the competence cascade also impacts biofilm formation and bacteriocin production^52,53^. The absence of *comCDE* in strains from several hunter-gatherer individuals suggests a potentially physiologically relevant difference between *S. mutans* strains present in populations practicing different subsistence strategies. While this result remains to be substantiated with larger datasets from diverse ancient populations, we underscore the potential utility of exploring ancient *S. mutans* accessory gene content to gain an understanding of virulence acquisition in this cariogenic taxon.

## Discussion and Conclusions

As observed in the archaeological record, the higher rates of dental caries in Neolithic populations are often linked to the adoption of carbohydrate-rich agricultural diets. However, dental caries is a multifactorial disease, and both diet and the oral microbiota play key roles in caries pathogenesis. While studies of both ancient and modern oral microbiota suggest a negligible effect of subsistence strategy on microbial community composition^18,20,21^, the potential selective impact of diet on individual cariogenic taxa remains underexplored. Here, we address this gap through presentation of genome-wide *S. mutans* data from 75 ancient individuals, including several hunter-gatherers. The largest collection of ancient *S. mutans* genomes presented to date, this dataset provides an unprecedented opportunity to explore patterns of relatedness in modern and ancient *S. mutans* strains.

Consistent with the findings of previous studies, we observe that *S. mutans* lineages do not cluster by country or continent of origin, suggesting a lack of phylogeographic substructure in ancient and modern populations^26^. This observation is particularly surprising given current models of *S. mutans* transmission, which suggest that children acquire strains primarily from parents and/or primary caregivers^54–56^. While the causes for this apparent contradiction remain to be elucidated, we hypothesize that the pervasiveness of recombination throughout the *S. mutans* core and accessory genomes may play a role; in modern populations with a recent history of interconnectivity and globalization, recombination from genetically divergent lineages may contribute to an apparent lack of phylogeographic substructure. For ancient metagenomic datasets, mismapping of short reads from closely-related oral taxa may introduce false-positive SNP calls and mask signals of clonal descent, particularly in regions with a recent history of horizontal gene transfer. Finally, analyses of heterozygosity indicate that many of the ancient individuals analyzed here harbor mixtures of multiple *S. mutans* strains. Challenges associated with separating mixed strains are exacerbated by the low coverage and highly fragmented nature of ancient DNA data; therefore, we have treated these chimeric genomes as single entities in population genetic analyses. Although necessarily coincident in time and space, we cannot exclude the possibility that mixed strains derive from divergent source populations, leading to uncertainty in PCA and phylogenetic placement.

Beyond the impacts of recombination, mismapping, and strain mixtures, biases associated with sample collection may contribute to the apparent lack of phylogeographic structure in modern *S. mutans* datasets. Currently available *S. mutans* assemblies derive primarily from clinical contexts in multiethnic urban centers and are often accompanied by incomplete metadata. Strain isolation location may not accurately reflect host genome geography, while admixture linked to a recent history of globalization may further obscure patterns of clonal descent. Furthermore, modern genomes from human populations practicing traditional lifeways and diverse subsistence strategies are currently completely lacking. While our ancient data suggests possible differences in competence-associated gene content in hunter-gatherer populations compared to agriculturalists, further sampling of both modern and ancient groups practicing diverse subsistence strategies is needed to substantiate these findings.

Alongside the need to generate additional datasets, we highlight the importance of performing *de novo* assembly instead of relying on mapping approaches alone to infer the gene content of past strains. The inflated genome size observed from the mapping approach versus that of *de novo* assembly could be explained by mis-mapping of other oral taxa. This could have led to wrong assumptions that ancient *S. mutans* displayed a larger genome than its modern counterparts, while *de novo* assembly points towards a similar genome size between ancient and modern strains. The analyses presented here show that inferring gene presence/absence by using the proportion of the gene covered and mean coverage normalisation are oversimplistic for species such as *S. mutans*, which form part of the oral microbiome, where closer relatives are also found in abundance, and that are prone to DNA uptake and recombination. Instead, future studies may consider performing *de novo* assemblies to explore the pan-genome variation of the species. A systematic comparison between the genes detected from *de novo* assemblies and mapping could help develop statistical models for inference of presence/absence of genes that take into account the significant variability in genome-wide coverage of datasets generated via hybridization capture. Finally, the apparent difference of the core genes in ancient strains and modern ones warrants further investigation, potentially highlighting genes important for modern *S. mutans* strains. As a starting point, future systematic comparison of datasets generated via hybridization capture and shotgun sequencing will be needed to exclude potential biases in core genome recovery, as well as inference in the genome size.

Methodological challenges associated with population genetic analysis and pan-genome reconstruction notwithstanding, our 75 newly-reported ancient *S. mutans* datasets provide unprecedented insights regarding both the antiquity and virulence of this cariogenic taxon.

Contrary to previous studies^30^, our recovery of *S. mutans* DNA from six geographically and temporally diverse hunter-gatherers and hunter-gatherer-fishers provides robust evidence for the widespread prevalence of *S. mutans* prior to the development and adoption of agriculture.

Similarly, modern *S. mutans* virulence-associated loci appear in some of our earliest reconstructed genomes, suggesting that the cariogenic potential of this microbe predated the widespread adoption of Neolithic diets. These observations are consistent with evidence for dietary heterogeneity and significant consumption of carbohydrate-rich plant foods in some hunter-gatherer populations (e.g. ^40,49^). We cannot exclude the possibility that *S. mutans* increased in prevalence following major dietary transitions, as suggested by some population genetic studies of modern and ancient *S. mutans* strains^26,29^. However, our dataset suggests that broad shifts in diet rather than changes in microbial virulence likely played a primary role in Neolithic-associated increases in dental caries rates. Together, the results presented here set the stage for further work exploring the acquisition of cariogenicity and pan-genomic evolution of *S. mutans* and other oral microbes using ancient DNA.

## METHODS

### Metagenomic Analysis

To identify ancient individuals exhibiting evidence of *S. mutans* preservation, we performed a metagenomic analysis of libraries previously produced by colleagues at the Max Planck Institute for Evolutionary Anthropology in Leipzig, Germany, the Max Planck Institute of Geoanthropology (formerly MPI for the Science of Human History, MPI-SHH) in Jena, Germany, the Australian Centre for Ancient DNA (ACAD) in Adelaide, Australia, and/or the Institute for Archaeological Sciences in Tübingen, Germany. While the majority of screened datasets were shotgun sequenced, we also screened off-target reads from libraries subjected to hybridization capture with a bait set targeting approximately 1.2 million polymorphic positions in the human genome^57–59^. Briefly, screening datasets were subjected to adapter clipping and read merging using either ClipAndMerge (v. 1.7.4) or AdapterRemoval (v. 2.3.2)^60,61^. Libraries processed using ClipAndMerge were quality trimmed with a threshold of 20 (-qt-q 20), and reads shorter than 30 bp were discarded^61^. For libraries processed with AdapterRemoval, an adapter overlap of at least 1 bp was required for read trimming (--minadapteroverlap 1), ambiguous bases and bases with a quality below 30 were trimmed on 3’ read ends only (--trimns--trimqualities --minquality 30 --preserve5p), and reads below 30 bp long were discarded (--minlength 30)^60^. For a subset of libraries produced using 5 or 7 bp internal barcodes, adapter clipping and demultiplexing were performed using one of two approaches. Some libraries had the P5 barcode clipped during demultiplexing, after which we ran AdapterRemoval with the reverse complement of the P7 barcode appended to the 5’ end of the adapter1 sequence^60^. For libraries with intact P5 and P7 barcodes, simultaneous demultiplexing and adapter clipping was performed using AdapterRemoval. Barcode sequences were supplied in a text file (--barcode-list), and we retained only reads with fewer than two mismatches in the mate 1 and mate 2 read barcodes (--barcode-mm 1)^60^. For all barcoded libraries, trimmed reads containing more than 3 ambiguous bases and/or shorter than 30 bp were discarded (--maxns 3 --minlength 30)^60^.

Following preprocessing, read mapping and metagenomic analysis was performed using the nf-core/eager pipeline (v. 2.4.6)^62^. After merging reads across sequencing lanes, libraries were mapped to the human genome hs37d5 using BWA aln (v0.7.17-r1188) with the following parameters:-n 0.01-l 1024-k 2-o 2 (ref. ^63^). Samtools (v. 1.12) was used to output unmapped reads in fastq format^64^. Metagenomic analysis was performed using the MEGAN alignment tool (MALT, v. 0.6.1) against a custom database containing a selection of bacterial, viral, and eukaryotic genomes from the RefSeq Genome database, including 6 *S. mutans* and 304 genomes of other Streptococci (**Supplementary Table 7**)^65^. MALT performed semi-global alignment (-at SemiGlobal) using BlastN mode (-m BlastN) with a minimum percent identity of 85 (-id 85) and a maximum alignments per query of 100 (-mq 100). Taxonomic binning was performed using the lowest common ancestor (LCA) algorithm with a top percent value of 1 (-top 1) and a minimum support value of 1 (-sup 1)^65^. Finally, we executed MaltExtract (v. 1.7) as implemented in the HOPS pipeline (Heuristic Operations for Pathogen Screening) to tabulate reads assigned to a user-defined list of taxa of interest, including *S. mutans*^34^. Libraries were evaluated based on the number of reads assigned to the *S. mutans* node, evenness of coverage across the reference, edit distance relative to the reference, and the presence of characteristic patterns of ancient DNA damage^34^. We selected 75 individuals for further analysis based both on these metrics and maximizing temporal and geographic diversity.

### Hybridization Capture Design and Evaluation

To enrich ancient libraries for *S. mutans* DNA, we designed a custom in-solution hybridization capture assay utilizing 191 modern *S. mutans* genomes available on the NCBI Genome database as of 2018-11-02 (**Supplementary Table 8**). Briefly, the combined sequences were used to generate two sets of 52 bp probes with an 8 bp 3’ linker sequence, as previously described^57^. Each probe set was produced using 6 bp tiling; by incorporating a 3’ offset between the two arrays, we attained an effective tiling density of 3 bp. Next, we used dustmasker (v. 2.2.32) to mask low-complexity regions in the probe sequences^66^. Redundant baits and those with more than 20% of their sequence masked were filtered, yielding 926,935 and 926,903 unique probes for the two sets, respectively. We ordered two 1 million Agilent SureSelect DNA Capture Arrays, randomly replicating probes to reach full capacity (968,000). Finally, baits were cleaved from the array surfaces to generate an in-solution DNA capture library following previously published methods^57^.

To assess the probe design’s taxonomic specificity, we used MALT (v. 0.4.0) to query the baits against the custom database described above (**Metagenomic Analysis**)^65^. Parameters followed those used in the metagenomic screening, except that the minimum percent identity was set to 90.0 (-id 90). In total, 1,243,261 of 1,451,926 assigned queries (85.6 %) were assigned to either the *S. mutans* species node or daughter strain nodes. 70,803 probes (4.9 %) were assigned at the *Streptococcus* genus level, while 118,409 of the aligned baits (8.2 %) matched other species within the genus *Streptococcus*. Assignment of the majority of probes to *S. mutans* and related taxa suggests that our capture reagent shows sufficient taxonomic specificity for application to ancient libraries.

### Sample Processing for Hybridization Capture

Following metagenomic screening for *S. mutans* DNA preservation, we selected 75 individuals from 61 archaeological sites for further analysis. All sample preparation steps had been performed in dedicated clean room facilities at the Australian Centre for Ancient DNA (ACAD) in Adelaide, Australia, the Institute for Archaeological Sciences in Tübingen, Germany, the Max Planck Institute of Geoanthropology (formerly MPI for the Science of Human History, MPI-SHH) in Jena, Germany, and/or the Ancient DNA Core Unit of the Max Planck Institute for Evolutionary Anthropology in Leipzig, Germany. One tooth from each individual was processed to obtain between 21.6 and c. 200 mg of bone powder following previously established methods (protocol: dx.doi.org/10.17504/protocols.io.bqebmtan). While some teeth exhibited carious lesions, in other cases no pathology was noted at the time of sampling.

For 74 individuals, DNA extraction was performed following either a modified version of a published silica column-based method (n=48; protocol: dx.doi.org/10.17504/protocols.io.baksicwe)^67^ or an in-solution silica-based approach (n=26)^59,68^ (**Supplementary Table 9**). In both protocols, bone powder was incubated overnight in an extraction buffer (0.45 M EDTA, 0.25 mg/mL proteinase K, pH 8) at 37°C with rotation^59,67,68^. For the column based method, c. 30-50 mg of bone powder was treated with 1 mL extraction buffer^67^, while the in-solution approach utilized c. 200 mg bone powder in 4 mL extraction buffer^59,68^. The in-solution method also included a second 55°C 1 hour incubation with proteinase K^59,68^. For the column-based extraction, the lysate was mixed with 10 mL binding buffer (5 M GuHCl, 40% Isoporpanol) and c. 400 µL 3M sodium acetate buffer (pH 5.2); following passage through a High Pure Extender Assembly (High Pure Viral Nucleic Acid Large Volume Kit, Roche), the DNA was treated with two ethanol washes and eluted in 100 µL TET (0.05% Tween-20 v/v)^67^. For the in-solution extraction, the lysate was combined with a medium-sized silica suspension in 16 mL binding buffer (90% Qiagen QG buffer with c. 5 M guanidinium thiocyanate, 220 mM sodium acetate, 25 mM sodium chloride, 1% triton X100).

Following a 1 hour binding step and two ethanol washes, DNA was eluted in 200 µL TET buffer (0.05% Tween-20 v/v)^59,68^. Finally, for individual PAN049, DNA was extracted from 60.8 mg of bone powder following an automated version of the silica-based method described above ^67,69^.

### 150 µL of lysate was processed using an automated liquid handling system (Bravo NGS Workstation B, Agilent Technologies) with silica-coated magnetic bead and binding buffer D in a final elution volume of 30 µL^69^

Next, for 74 individuals, extracted DNA was prepared into double-stranded, double-indexed libraries using either (1.) a UDG-half library preparation approach which repairs internal deaminated cytosines while preserving signals of damage at the termini of DNA fragments (n=70), or (2.) a full-UDG treatment utilizing USER enzyme for the removal of all deaminated cytosines in ancient molecules (n=4) (**Supplementary Table 9**). For the UDG-half treated dataset, a subset of 23 libraries were prepared in dedicated clean-room facilities in Adelaide following a previously published protocol^70^, while the remaining 47 UDG-half treated libraries were produced in Jena or Tübingen following a modified approach (protocol: dx.doi.org/10.17504/protocols.io.bmh6k39e^70,71^). For libraries prepared in Adelaide, library preparation began with a damage repair step utilizing between 15 and 30 µL of extract and USER enzyme mix (Buffer Tango: 1 x, dNTPs: 100 µM each, ATP: 1 mM, USER Enzyme: 0.06 U/µL) in a final volume of 52.2 µL^70^. For libraries prepared in Jena/Tübingen, 25 µL of extract was treated with 25 µL USER enzyme mix (Buffer Tango: 1.2 x, ATP: 1.2 mM, BSA: 0.2 mg/mL, dNTPs: 0.1 mM each, USER Enzyme: 0.072 U)^70,71^. In both protocols, libraries were incubated for 30 min. at 30°C, after which the reaction was inhibited by adding 3.6 µL UGI (2 U stock concentration), followed by another 30 min. incubation at 37°C^70,71^. Next, fragments were blunt-end repaired by adding T4 PNK and T4 Polymerase; libraries prepared in Adelaide were treated with 4.2 µL enzyme mix (T4 PNK: 0.5 U, T4 Polymerase: 0.1 U), while those produced in Jena/Tübingen required 4.65 µL enzyme mix (T4 PNK: 0.515 U, T4 Polymerase 0.085 U)^70,71^. In both protocols, libraries were then incubated at 25°C for 20 min. and 12°C for 10 min., followed by a cleanup using a MinElute PCR purification Kit (Qiagen). Adapter ligation was performed in a total volume of 40 µL (T4 DNA ligase buffer: 1 x, P5/P7 adapters: 0.25 µM each, T4 DNA ligase: 0.125 U)^70,71^; for libraries prepared in Adelaide, the reaction also included 5% PEG-4000, and P5 and P7 adapters included a unique library-specific combination of 5 or 7 bp barcode sequences to aid in detection of cross-contamination^70^. Libraries prepared in Adelaide were incubated for 30 min. at room temperature, while those produced in Jena/Tübingen were incubated for 20 min. at 22°C. In both protocols, the adapter ligation was followed with a second MinElute cleanup (Qiagen)^70,71^. For libraries prepared in Adelaide, adapter fill-in was performed in a total volume of 40 uL (ThermoPol buffer: 1 x, dNTPs: 250 µM each, Bst Polymerase: 0.4 U), and libraries were incubated for 20 min. at 37°C followed by 20 min. at 80°C^70^. Libraries prepared in Jena and Tübingen utilized a mastermix containing 1 x Isothermal Buffer, dNTPs at a concentration of 0.125 each, and 0.4 U Bst Polymerase, and the fill-in reaction included a 30 min. incubation at 37°C followed by 10 min. at 80°C^70,71^.

For the full-UDG treated dataset, 3 libraries were prepared in dedicated clean-room facilities in Adelaide following a previously published protocol^72^, while the one library was produced in Tübingen following a modified approach (protocol: dx.doi.org/10.17504/protocols.io.bqbpmsmn^71,73^). In contrast to the UDG-half protocols, for full-UDG libraries excision of deaminated cytosines and blunt-end repair were performed simultaneously. Between 50 and 150 µL of mastermix was added, depending on the amount of input extract (NEB Buffer 2: 1x, T4: 0.4 U, USER: 1 U), after which the reactions were incubated at 37°C for 3 hours^71–73^. Next, libraries produced in Adelaide were treated with T4 polymerase (0.059 U) followed by a 30 min. incubation at 25°C^72^, while the protocol used in Jena/Tübingen utilized 0.115 U T4 polymerase and an incubation of 20 min. at 25 °C followed by 10 min. at 12°C^71,73^. Both protocols included a MinElute purification step, after which adapters were ligated using a mastermix containing 1x Quick ligase buffer and 0.125 U Quick Ligase. For libraries produced in Adelaide, the ligation mastermix included P5 and P7 adapters with a unique library-specific combination of 5 or 7 bp barcode sequences (0.0625 µM), after which the reaction was incubated for 5 min. at 25°C^72^. Libraries produced in Tübingen utilized short non-barcoded adapters at a final concentration of 0.25 µM, and reactions were incubated for 20 min. at 22°C^71,73^. Following a second MinElute purification step, both protocols included an adapter fill-in reaction. For libraries produced in Adelaide, the 50 µL reaction was incubated at 37°C for 30 min., followed by 20 min. at 80°C (Thermopol buffer: 1x, dNTPs:.25 µM each, *Bst* DNA polymerase:.32 U)^72^; the Tübingen protocol, on the other hand, included an incubation of 37°C for 30 min. followed by 10 min. at 80°C (40 µL reaction, Isothermal buffer: 1x, dNTPs: 0.125 mM each, *Bst* DNA polymerase: 0.4 U)^71,73^.

Finally, 30 µL of DNA extracted from individual PAN049 was prepared into a single-stranded, partially UDG treated library following an automated version (protocol: dx.doi.org/10.17504/protocols.io.kqdg32bdpv25/v1) of previously published protocols^74,75^.

Briefly, uracils in the interior of DNA strands were excised via addition of *E. coli* Uracil-DNA-glycosylase and *E. coli* endonuclease VIII to the dephosphorylation master mix. Two quantitative PCR assays were performed to estimate DNA yields and quantify the efficiency of library preparation^75^.

Following qPCR quantification, all libraries were amplified and tagged with sample-specific barcoded Illumina adapters. For libraries prepared in Adelaide, a volume of 13.4 µL was used for isothermal amplification with the TwistAmp Basic kit (TwistDx Ltd). The reaction proceeded for 44 min. at 37°C, after which libraries were purified using gel electrophoresis and quantified via Nanodrop^59^. Samples processed in Jena and/or Tübingen were indexed and amplified using a Pfu Turbo Cx Polymerase as described elsewhere (protocol: https://www.protocols.io/view/illumina-double-stranded-dna-dual-indexing-for-anc-4r3l287x3l1y/v2). Single-stranded libraries produced in Leipzig were indexed and amplified using AccuPrime *Pfx* DNA polymerase as described elsewhere^75^. Finally, indexed, amplified libraries were subjected to hybridization capture using the custom in-solution bait set described above and following previously described methods^57^. Captured libraries were sequenced to a depth between 3,000,000 and 41,000,000 reads on a HiSeq 4000, a NextSeq 500, or a NovaSeq X Plus. All sequencing was performed at either the Max Planck Institute of Geoanthropology (formerly MPI for the Science of Human History, MPI-SHH) or the Max Planck Institute for Evolutionary Anthropology using paired end chemistry (2 x 76 + 8 + 8 or 2 x 101 + 8 + 8 cycles).

### Modern Comparative Dataset

To obtain a modern comparative dataset for population genetic analysis, we downloaded 459 modern, annotated *S. mutans* genomes available from the NCBI Refseq database as of 2023-10-18 (**Supplementary Table 3**). Two genomes marked as “atypical” were excluded from downstream analysis. Next, we processed the modern genomes using a custom script (https://github.com/alexherbig/Genome2Reads) to generate 100 bp reads with 1 bp tiling. We mapped the resulting artificial reads to the *S. mutans* UA159 reference genome (GCA_000007465.2) using the nf-core/eager implementation of BWA aln with strict parameters (-n 0.1-l 32)^24,62,63^. Following deduplication with Picard MarkDuplicates (v. 2.26, http://broadinstitute.github.io/picard/), we genotyped the modern strains using the GATK UnifiedGenotyper (v. 3.5) with default parameters and the output mode ‘EMIT_ALL_SITES’^76^.

Next, we identified the set of core regions in the UA159 reference genome shared across modern *S. mutans* isolates. Briefly, we filtered the modern genotype calls using MultiVCFAnalyzer (v. 0.87); we required a minimum coverage of 5 and a minimum genotype quality score of 30 for a base call, and heterozygous SNPs were excluded by setting the minimal allele frequency for both a homozygous and a heterozygous call to 0.9^77^. After excluding the reference from the resulting full genome alignment, we used SeqKit (v. 2.4.0) to identify and exclude one duplicate sequence (ASM346685v1, GCF_003466855.1)^78^. Using a custom python script (multifasta_core.py), we parsed the full genome alignment to produce a bed file containing all positions in the UA159 reference genotyped in at least 95% of modern *S. mutans* strains. Adjacent windows were merged with BEDtools merge (v. 2.25.0), and we defined the UA159 core genome as the set of regions at least 500 bp in length present in at least 95% of the 456 modern comparative genomes^79^. Finally, we used BEDtools complement (v. 2.25.0) to identify accessory regions of the UA159 reference falling outside this core genome set^79^. We merged these non-core windows with positions falling into highly conserved genomic regions susceptible to mismapping from environmental taxa, including loci encoding transfer RNAs, ribosomal RNAs, and one transfer RNA pseudogene (GCA_000007465.2_ASM746v2_regions2exclude_rRNA_tRNA.gff). Together, the accessory and conserved genomic regions comprise 515,186 bp (25.3 %) of the 2,032,925 bp UA159 reference genome; unless stated otherwise, these sequences were excluded from subsequent population genetic analyses (GCA_000007465.2.noncore.regions_exclude.bed).

### Ancient Data Preprocessing and Authentication

Preprocessing of shotgun sequencing and capture datasets was performed separately for libraries with and without internal barcodes. For samples with internal barcodes, poly-g tails of at least 10 bp were trimmed using fastp (v. 0.20.1,-g --poly_g_min_len 10)^80^. Next, AdapterRemoval (v. 2.3.2) was used for simultaneous adapter trimming, demultiplexing, and merging of reads. A separate text file containing library-specific internal barcode sequences was supplied using the flag --barcode-list, and only reads with the expected barcode combinations were retained^60^. Ambiguous and low-quality bases were trimmed while preserving 5’ read ends (--trimns --trimquailities --minquality 20 --preserve5p), and reads shorter than 30 bp were removed (--minlength 30). We required a minimum adapter overlap of 1 bp (--minadapteroverlap 1) for trimming^60^. Following adapter clipping, merged and unmerged reads with correct barcode combinations were concatenated, read prefixes were updated for downstream compatibility using AdapterRemovalFixPrefix (v. 0.0.5; https://github.com/apeltzer/AdapterRemovalFixPrefix), and libraries were merged across lanes.

For libraries without internal barcodes, including published datasets from individual KGH-2B^29^, poly-g trimming, adapter clipping, and read merging was performed within the nf-core/eager pipeline (v. 2.4.6)^62^. Briefly, FastQC v. 0.11.9 was executed both before and after adapter trimming to assess sequence quality^81^. For samples sequenced on either an Illumina NextSeq 500 or a NovaSeq X Plus, poly-g trimming was performed using fastp (v. 0.20.1) with the same parameters as the barcoded libraries, except that length filtering of reads below 15 bp was disabled using the flag-L^80^. AdapterRemoval (v. 2.3.2) was executed as for the barcoded libraries, except that the demultiplexing step was omitted^60^. Barcoded and non-barcoded libraries were processed identically using the nf-core/eager pipeline for all downstream analysis steps.

Next, we mapped all newly-generated libraries as well as previously published non-UDG-treated libraries from KGH-2B (ERR11660452, ERR11660453, ERR11660454, ERR11660455) to the *S. mutans* UA159 genome assembly to check for characteristic patterns of ancient DNA damage and fragmentation^24^. Mapping was performed using the nf-core/eager implementation of BWA aln (v. 0.7.17-r1188)^63^ with the following parameters:-n 0.01,-l 16.

We filtered alignments using Samtools (v. 1.12) with a quality threshold (-q) of 37, and deduplication was performed using MarkDuplicates (Picard) with default parameters (http://broadinstitute.github.io/picard/)^64^. Cytosine deamination rates and fragment length distributions were evaluated using DamageProfiler (v. 0.4.9), and the results of nf-core/eager data processing steps were summarized using MultiQC (v. 1.13)^82,83^. Finally, as previous studies have documented correlation between damage rates on reads mapping to endogenous ancient microbial and human genomes^36,84^, we repeated the above damage assessment steps after mapping to the human genome (HG19). All other parameters were kept constant, and DamageProfiler (v. 0.4.9) was used to evaluate deamination patterns and read lengths of human mapping reads from the same libraries (**Figure 2)**^82^.

Following this initial quality control assessment, reads were trimmed to remove positions affected by cytosine deamination prior to genotyping. Briefly, adapter-trimmed and merged UDG-half libraries were processed using the nf-core/eager implementation of fastP (v. 0.20.1) to remove two base pairs from 5’ and 3’ read ends (--trim_front1 2 --trim_tail1 2)^62,80^. Adapter clipping, polyG trimming, quality filtering, and length filtering were disabled using the following flags:-A-G-Q-L. Trimmed UDG-half and untrimmed full-UDG libraries were mapped to the *S. mutans* UA159 reference genome using BWA aln (v. 0.7.17-r1188) with strict parameters:-n 0.1-l 32^24,63^. Alignments were filtered using Samtools (v. 1.12) with a mapping quality threshold (-q) of 37^64^. The endogenous DNA content of each library was computed using EndorS.py (v 0.4; https://github.com/aidaanva/endorS.py), after which duplicate removal was performed using the nf-core/eager implementation of Picard MarkDuplicates (v. 2.26.0) with default parameters (http://broadinstitute.github.io/picard/)^62^. Damage profiles following clipping were generated using DamageProfiler (v. 0.4.9) as described above, and coverage statistics were computed using Qualimap (v2.2.2-dev)^82,85^. Finally, genotyping was performed using the GATK UnifiedGenotyper (v. 3.5) with the output mode set to ‘EMIT_ALL_SITES’, and results were summarized using MultiQC (v. 1.13)^76,83^.

To identify samples exhibiting evidence of multiple *S. mutans* strains and/or contamination from closely related taxa, we used MultiVCFAnalyzer (v. 0.87) to explore the distribution of multiallelic positions in our dataset^77^. We filtered genotype calls with a GATK genotyping quality score below 30 and/or a base coverage less than 5. Finally, we excluded positions falling into highly conserved genomic regions susceptible to mismapping from environmental taxa (GCA_000007465.2_ASM746v2_regions2exclude_rRNA_tRNA.gff).

Genotypes were called using a minimal allele frequency for a homozygous call of 0.9, a minimal allele frequency for a heterozygous call of 0.1, and outputting allele frequencies for each variant position^77^. Finally, in order to identify cases of strain mixture and/or background contamination, we visualized the proportion of reads supporting the derived allele across SNP positions (**Supplementary Figure 1**)^36^.

To reduce the impact of read mismapping and/or background contamination on subsequent analyses, we performed an additional filtering step using MALT to extract reads taxonomically assigned at or below the *S. mutans* species node^65^. Briefly, preprocessed reads were mapped to the human genome (hs37d5) using BWA aln (v. 0.7.17-r1188) with the following parameters:-n 0.01,-l 1024,-k 2,-o 2. Next, we used the nf-core/eager (v. 2.4.6) implementation of MALT (v. 0.6.1) to align unmapped reads against a custom database including all sequences available in the NCBI nucleotide database (ftp://ftp-trace.ncbi.nih.gov/blast/db/FASTA/) as of Oct. 26, 2017^62,65^. Parameters followed those outlined previously (**Metagenomic Analysis**), except that we set the minimum percent identity to 90 (-id 90). Next, we visualized MALT’s.rma6 output using the MEtaGenome ANalyzer (MEGAN6) to identify reads taxonomically assigned at or below the *S. mutans* species node^86^. Seqtk subseq (v. 1.2-r94) was used to extract these reads from the preprocessed fastq files (https://github.com/lh3/seqtk). To evaluate filtering efficacy, we compared the distribution of multiallelic positions in each sample before and after MALT filtering. Briefly, damage-trimmed, filtered fastq files were mapped to the UA159 reference genome, genotyped, and processed using MultiVCFAnalyzer (v. 0.87) as described above^77^. Following visualization of derived allele frequencies for multiallelic positions, we observed that filtering with MALT did not cause appreciable loss of SNP positions and reduced the prevalence of heterozygous SNP calls in some samples (**Supplementary Figure 1**); thus, unless otherwise stated, we utilized the MALT-filtered dataset for downstream analysis.

### Population Genetic Analysis

Prior to merging our ancient and modern datasets for phylogeographic analysis, we filtered genotype data from 456 modern genomes (**Modern Comparative Dataset**) to identify a set of high-quality, biallelic segregating SNPs falling within core regions of the UA159 reference genome. Briefly, we filtered variants using MultiVCFAnalyzer (v. 0.87) as described above, requiring a minimum base coverage of 5 and a minimum genotype quality score of 30, setting the minimum allele frequency for both homozygous and heterozygous base calls to 0.9, and excluding variants falling in non-core and conserved regions of the UA159 reference genome^77^. Using a custom python script (multivcf_to_eigenstrat.py), we further filtered the SNP alignment output by MultiVCFAnalyzer to exclude multiallelic and invariant positions, as well as sites genotyped in less than 95% of our modern strains. The resulting alignment consisted of 118,675 SNP positions genotyped in 456 modern isolates and was output in multifasta and EIGENSTRAT formats.

Next, we performed principal components analysis to explore population structure in modern and ancient *S. mutans* strains (**Ancient Data Preprocessing and Authentication**). As described above, we used MultiVCFAnalyzer (v. 0.87) to filter variants in 456 modern genomes, one ancient published strain, and 75 newly-reported ancient *S. mutans* datasets, lowering the minimum coverage for a base call to 3 to facilitate analysis of low-coverage samples ^77^. Using a custom script (multivcf_to_eigenstrat.py), we filtered the resulting SNP alignment to exclude multiallelic sites and output the data in eigenstrat format. Using the trident software (v. 1.1.6.0), we subset the eigenstrat data to include only the 118,675 high-quality, biallelic core SNP positions ascertained in our modern dataset (trident forge --selectSnps)^87^. Finally, we produced a principal components analysis (PCA) of modern and ancient *S. mutans* genetic variation using smartPCA with default parameters (v. 16000, EIGENSOFT v7.2.1)^88,89^. Modern strains were subdivided into populations based on the country of isolation. smartPCA computed principal components based on 443 modern samples (excluding outliers), and ancient individuals with a median coverage of the UA159 reference genome greater than or equal to 3x were projected onto these axes of variation (lsqproject: YES).

Finally, we performed phylogenetic analysis to further explore patterns of relatedness between ancient and modern *S. mutans* strains. To mitigate biases introduced via missing data, we generated a complete deletion alignment using modern assemblies, published ancient datasets, and newly-reported ancient *S. mutans* strains with a median coverage of the UA159 reference genome greater than 10x. Modern and ancient genotypes were filtered using MultiVCFAnalyzer (v. 0.87) as described above with a minimum read depth of 5 for a base call^77^. The output of MultiVCFAnalyzer was further filtered to exclude multiallelic SNPs and sites with missing data, yielding a 40,679 bp alignment which was output in multifasta format. Finally, we computed a maximum likelihood phylogenetic tree using RAxML-NG (v. 1.1) with a general time reversible (GTR) substitution model and gamma rate variation with four site categories (G4)^90^. We used the Stamatakis ascertainment bias correction (ASC_STAM), counting invariant sites in the complete fasta alignment using the command snp-sites-C (v. 2.5.1)^91^. Support values were estimated using 1,000 bootstrap replicates.

### Pan-Genome Reconstruction

In order to incorporate the ancient *S. mutans* genomes into the pan-genome reconstruction, we *de novo* assembled ancient genomes with a minimum mean coverage of 10x after capture, as well as the previously published ancient shotgun genome KGH2-B^29^, with nf-core/mag^92^ with the following command: nextflow run nf-core/mag-profile eva,archgen,local_paths-r 3.4.0 --input input.csv --outdir results --clip_tool adapterremoval --host_fasta hs37d5.fa --ancient_dna --skip_spades --binqc_tool checkm --refine_bins_dastool --postbinning_input both --skip_prokka-c mag.config. The config file can be found here: https://github.com/meganemichel/mutans_project_scripts. The *S. mutans* MAG for each sample were identified based on the results from GTDB-Tk (v.2.4.0)^93^ and the best quality MAG for each sample was selected to maximise completeness, minimise contamination and maximise the number of genes found in the assembly as reported by CheckM (v.1.2.3)^51^ and metaQUAST (v.5.0.2)^94^.

Next, we used Roary (v.3.13.0)^95^ to construct a pan-genome encompassing the variation present in 456 modern *S. mutans* assemblies downloaded from the NCBI Genome database (**Modern Comparative Dataset**), as well as the newly assembled 35 ancient *S. mutans* MAGs. Assembly level of modern strains ranged from contigs to complete genomes. For consistency, all genomes were re-annotated using Prokka (v. 1.14.6)^96^ with default parameters and specifying the genus and species as follows: --genus Streptococcus --species mutans. The output was written in Genbank/ENA/DDJB format using the flag --compliant^96^. Roary (v. 3.13.0) was used for pan-genome construction, specifying that a gene cluster must be present in 95% of the strains to be considered core (-cd 95)^95^. Following conventions used by Roary, we define genes present in more than 95% of *S. mutans* strains as core, while genes present in less than 95% of strains are termed accessory. The matrix produced by Roary (gene_presence_absence.Rtab) was used as input for the Rnotebook to calculate the number of genes per assembly. We plotted the number of genes in correlation to MAG completeness and mean coverage, and coloured the plots based on contamination (as estimated by CheckM), dating of the individual, or whether a strain mixture was detected. Only a correlation with the completeness was observed, and none of the colouring criteria showed a trend. The gene_presence_absence.Rtab matrix was also used to re-compute the pan-genome for only the ancient assemblies and another one for only the modern assemblies, classifying genes as described above: if present in 95% or more of the assemblies the gene is considered core, and otherwise an accessory gene.

To test whether the *S. mutans* hybridization capture baits span a majority of our reconstructed pan-genome, we mapped the probe sequences to the multifasta reference generated by Roary. Briefly, baits were aligned using the nf-core/eager (v. 2.4.6) implementation of BWA aln (v. 0.7.17) with strict mapping parameters (-n 0.1-l 32)^62,63^. Alignments were filtered using Samtools (v. 1.12) with a mapping quality (-q) of 0, after which deduplication was performed with Picard MarkDuplicates (v. 2.26.0; http://broadinstitute.github.io/picard/)^64^. Finally, using BEDtools (v. 2.25.0) we computed the breadth and the depth of coverage for each pan-genome cluster^79^. The bed files were used as input for the Rnotebook and we considered a gene cluster as covered if it had a minimum of 90% of its length covered. After verifying that the probes span a majority of pan-genome gene clusters (**Pan Genome Reconstruction**), we repeated the mapping, filtering, deduplication, and coverage computation to assess pan-genomic coverage using as input the MALT-filtered ancient hybridization capture datasets (**Ancient Data Preprocessing and Authentication**). All steps were performed as described for the hybridization capture probes. A gene was considered covered if it had an equal or higher breadth of coverage compared to the breadth of coverage when mapped to the reference genome (**Ancient Data Preprocessing and Authentication**), and a depth between half and two times the depth of the dataset mapped to the reference genome. We compared the number of reads detected in the *de novo* assembly, with the ones detected during mapping, and the union of both (referred as Ancient Merged) with the function ggbetweenstats of the ggstatsplots (v.0.13.0) R package^97^.

All the code for the analysis described in this section can be found here, including that for generating the figures: https://github.com/meganemichel/mutans_project_scripts.

### Assessment of Virulence-Associated Gene Content

Finally, to explore the evolution of virulence using our ancient datasets, we identified a subset of previously-reported virulence-associated loci in the *S. mutans* UA159 reference genome^27^. We mapped pre-processed, MALT-filtered data to the UA159 reference as described above (**Ancient Data Preprocessing and Authentication**). Supplying virulence-associated loci in bed format, we used a custom script to compute the proportion of each virulence-associated gene covered to a depth of at least 1x.

## ACKNOWLEDGEMENTS

1. M. M., G.U.N., E.S., P.W.S., V.V.-M, and C.W. received support from the Max Planck-Harvard Research Center for the Archaeoscience of the Ancient Mediterranean. We thank the Ministerio de Ciencia, Innovación y Universidades for their support of R.M.P. (grant PID2023-146504NB-I00, Programa ICREA Acadèmia), C.O.C. (grant PID2023-146504NB-I00), and C.R.H. (grant PID2023-146504NB-I00, Generalitat de Catalunya-DGR). M.D. and M.E. were supported by the Praemium Academiae Award of the Czech Academy of Sciences. We thank the Kone foundation, the Otto Malm foundation, and the Foundation for Research on Viral Diseases for supporting E.K.G. G.R.S. and D.G. received support through the University of Otago Research Grant. G.C. received support from the Australian Research Council Grant DP200102872. J.F.F. received support from the Generalitat de Catalunya. A.Strauss was supported by funds from the FAPESP (17/16451-2). C.W. received support from the Werner Siemens Foundation and the DFG Excellence strategy (EXC 2051 #390713860). Thank you to Luca Bondioli, Michael Kunst, Rainer Linke, Peter van Dommelen, and Noreen Tuross for facilitating access to skeletal materials for archaeogenetic study. Some of the data presented here was generated by the Ancient DNA Core Unit of the Max Planck Institute for Evolutionary Anthropology, which is funded by the Max Planck Society. We thank the laboratory staff in the Department of Archaeogenetics at the former Max Planck Institute for the Science of Human History, as well as the Core Unit of the Max Planck for Evolutionary Anthropology for supporting generation of ancient DNA datasets, and Kay Prüfer for handling the raw sequencing data.

## AUTHOR CONTRIBUTIONS

A.A.V. conceived of and initiated the study under the supervision of A.H. and J.K.; C.L.-F., B.L., K.N., P.O., C.P., A.Sajantila, S.S., M.A.S., P.W.S., E.K.G. and W.H. organized and assembled datasets for ancient genomic analysis. M.A., C.A.-F., M.A.C., M.A.-V., G.A.J., E.B., M.I.B.R., P.B.F., S.B., A.B.B., N.B., K.B., Y.B., A.B., L.C., J.E.C.H., E.C.B., G.C., M.Daubaras, P.d.M.I., M.D.-Z.B., M.Dobeš, V.D., Y.S.E., M.E., L.F., J.F.F., J.L.F., S.F., D.G., G.G.A., R.G.P., J.A.G.-V., M.G.G., M.G.L., E.K.G., S.H., D.I.H.-Z., T.I., R.J., J.J.E., A.R.K., K.K., T.K., I.L.-C., V.L., R.G.M., L.M., A.M.R., V.E.M., K.Massy, H.M., R.M.P., L.M.G., A.G.M., C.O.C., A.O., C.R.P., E.Pompianu, J.P.G., E.Protopapadaki, S.R., C.R.H., R.R., M.A.R.G., A.R., D.C.S.-G, T.Salie, M.S.R., D.S., T.Schunke, R.Shafiq, R.Shing, R.Skeates, A.Sperduti, A.Strauss, G.R.S., A.S.-N., C.T.R., F.V., A.V., C.V.F., M.V.S., E.W., K.A.Y., G.Z., and W.Z. performed archaeological excavation, anthropological assessment, and/or curation of analyzed skeletal materials and facilitated the compilation of archaeological contextual information. R.B., T.F., A.N.H., K.Majander, A.M., E.A.N., G.U.N., L.P., C.P., E.S., A.Sevkar, M.A.S., G.V., and V.V.-M. curated datasets and performed computational and laboratory work. A.A.V. and A.H. designed the *S. mutans* hybridization capture reagent. M.M. conducted metagenomic screening of ancient DNA datasets. M.M. performed bioinformatic processing and computational and population genomic analysis of *S. mutans* hybridization capture datasets. A.A.V. produced and analyzed ancient *S. mutans de novo* assemblies. A.A.V and M.M analyzed the *S. mutans* pan-genome. M.M., A.A.V., A.H., and J.K. wrote the manuscript with input from all coauthors.

## DATA AND CODE AVAILABILITY

Data and code needed to reproduce these analysis are available on github: https://github.com/meganemichel/mutans_project_scripts

Raw sequencing data associated with this project has been deposited in the European Nucleotide Archive, accession number PRJEB104722.

## SUPPLEMENTARY FIGURES

**Supplementary Figure 1.**
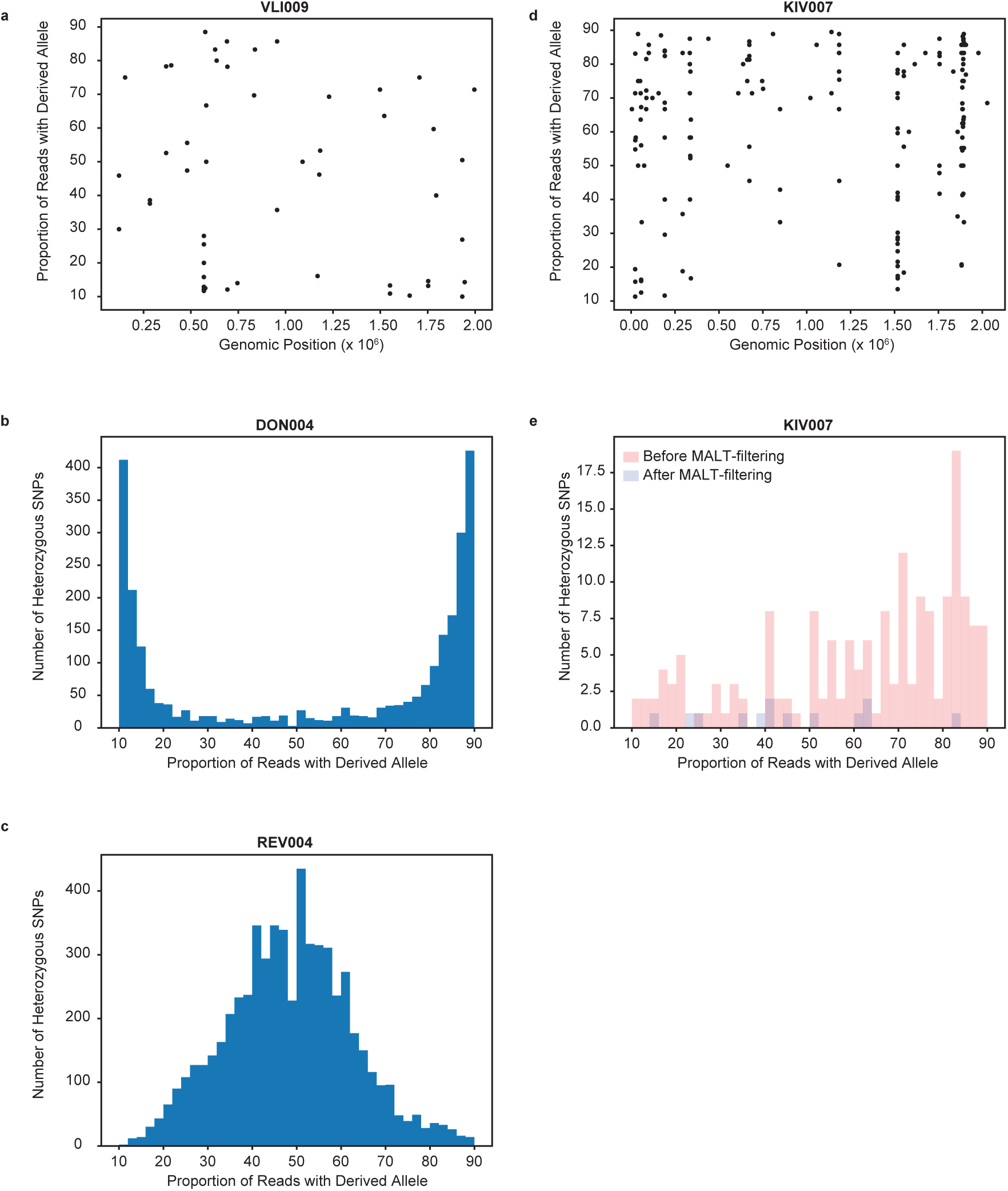
Strain heterozygosity in ancient *S. mutans* genomes (previous page) Analysis of strain heterozygosity illustrating **a.** monomorphic *S. mutans* strains, **b.-c.** mixed-strain infections, and **d.-e.** background contamination in ancient *S. mutans* genomes. For each genome, SNP calling was performed using MultiVCFAnalyzer with a minimum coverage threshold of 5. Heterozygous SNPs were defined as those having a frequency of the derived allele between 0.1 and 0.9. Unless otherwise stated, plots depict SNPs called after MALT-filtering. **a.** Scatterplot depicting the genomic position and percentage of reads supporting the derived allele for heterozygous SNP calls in the VLI009 *S. mutans* genome. **b.-c.** Histograms binning heterozygous SNPs by the percentage of reads supporting the derived allele in the VLI009 and DON004 *S. mutans* genomes, respectively. **d.** Scatterplot depicting the genomic position and percentage of reads supporting the derived allele for heterozygous SNP calls in the KIV007 *S. mutans* genome. **e.** Histogram showing the percentage of reads supporting the derived allele for heterozygous SNP calls in the KIV007 genome both before (red) and after (blue) MALT filtering.

**Supplementary Figure 2.**
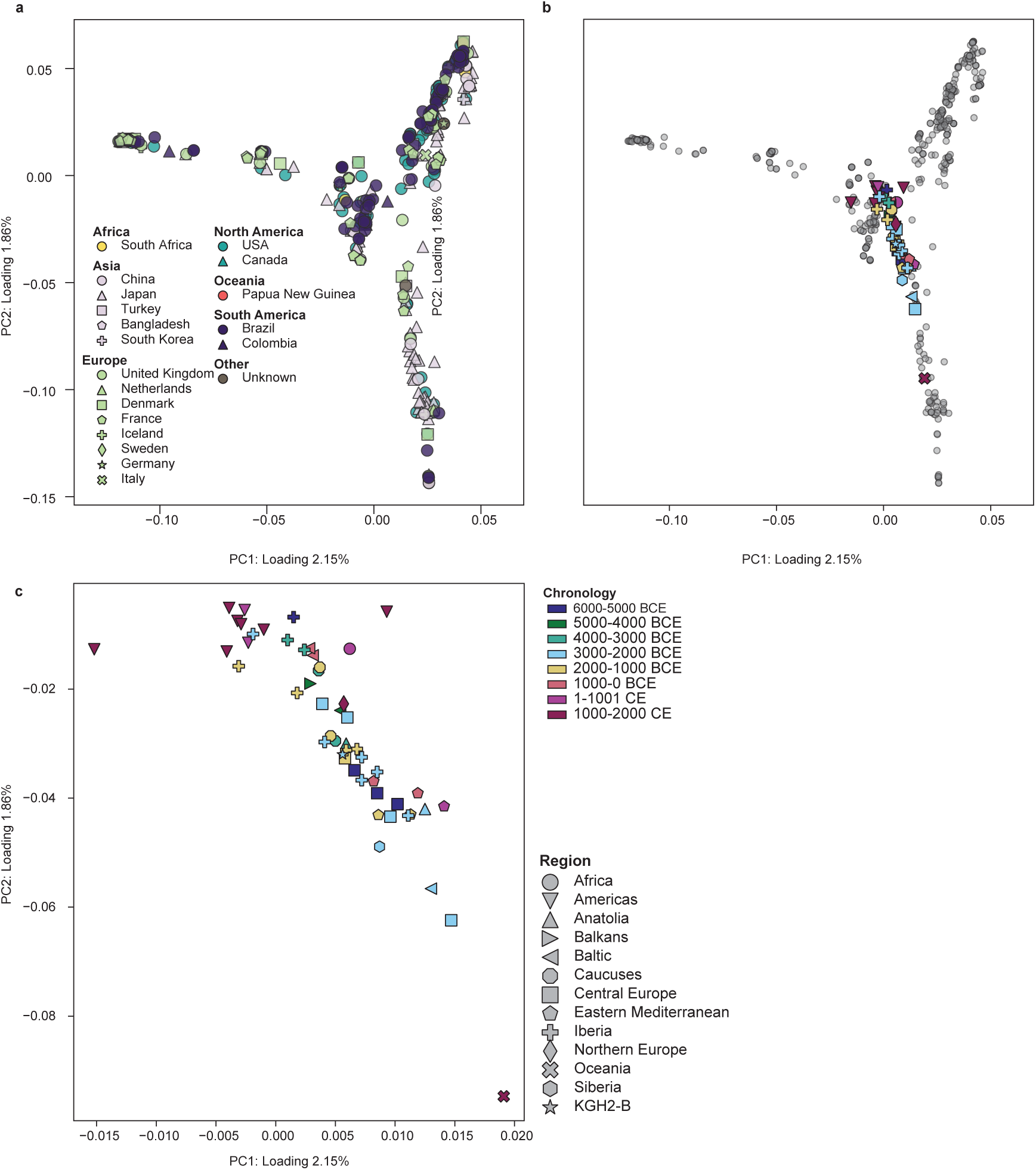
Principal components analysis of modern and ancient *S. mutans* strains **a.** Principal components analysis including 456 modern, publicly available *S. mutans* assemblies. Strain coloration reflects regional groupings, while shape indicates the country of origin. **b.** Ancient strains (≥ 3x median coverage) projected onto the diversity of modern *S. mutans* clones (small gray circles). Coloration reflects millennial age estimations of ancient genomes, while marker type indicates geographic region of origin. **c.** Zoomed view of panel b showing relative positioning of ancient *S. mutans* strains in PCA space. Modern *S. mutans* assemblies have been omitted for clarity.

**Supplementary Figure 3.**
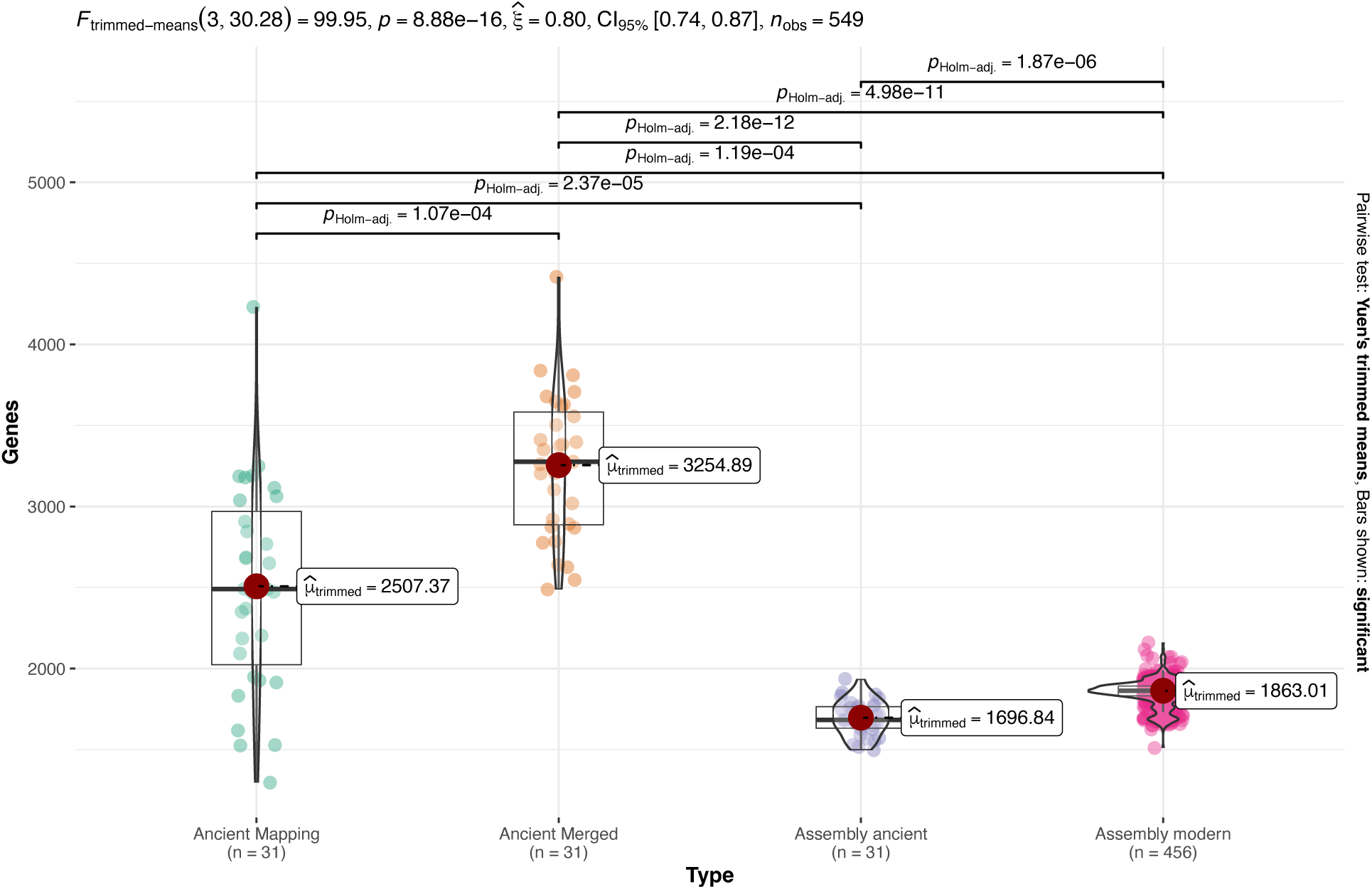
Genome size estimated for modern genomes and ancient genomes. For ancient genomes the estimated genome size was done based on the assemblies (Assembly ancient), the mapping of the reads to the pan genome reference (Ancient Mapping) and the union of both (Ancient Merged).

**Supplementary Figure 4.**
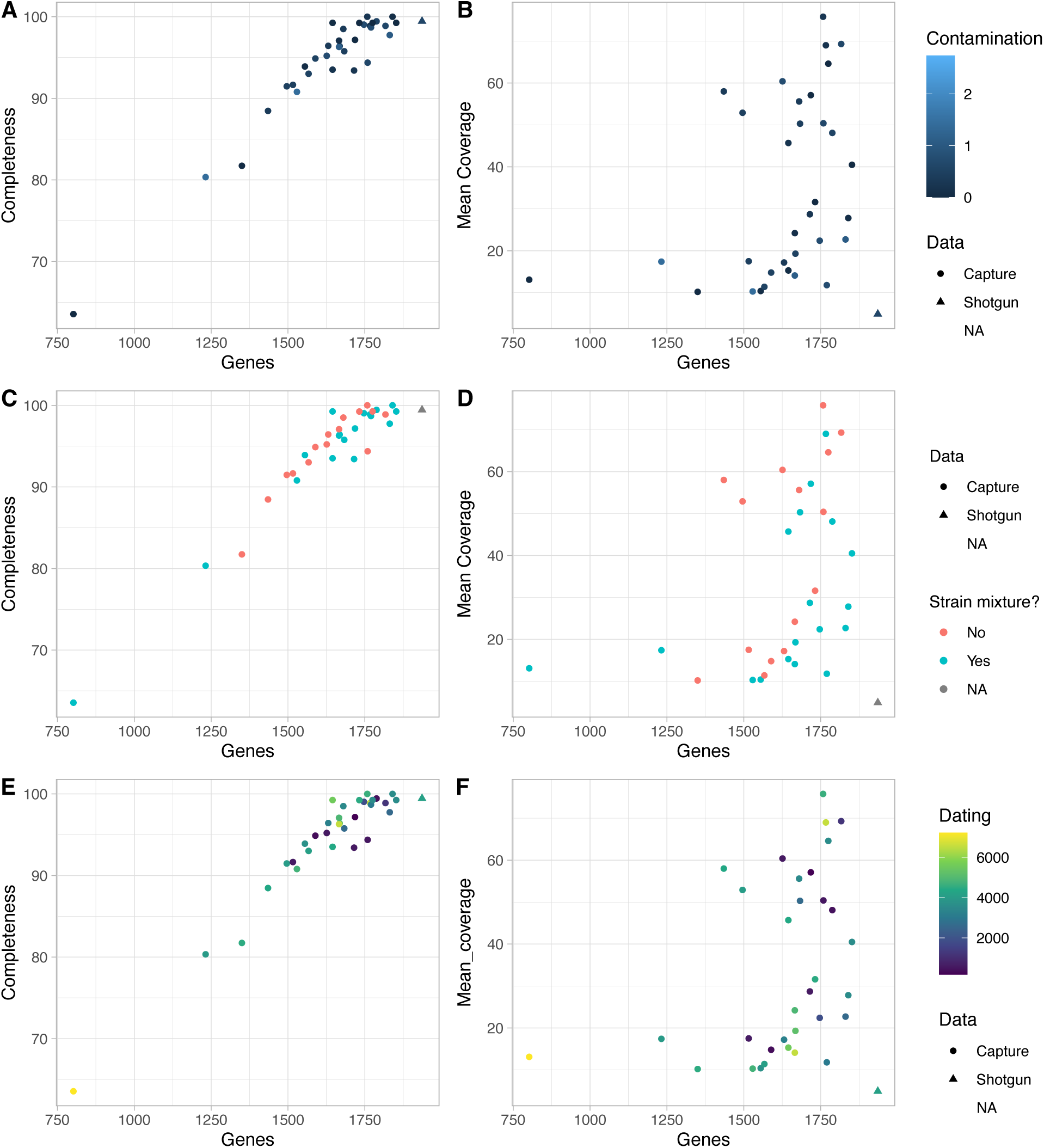
Correlations between gene counts and completeness or mean coverage. Gene content compared to completeness (A,C, E), mean coverage (B,D,F), coloured either by the contamination of the genomes (A,B), whether a strain mixture or contamination has been detected (C,D) or their mean Before Present dating (E,F). The triangle represents shotgun genomes while the dots represent capture datasets.

