## Supplementary Notes 1 and 2 for "Exploring the Evolution of the Cariogenic Oral Pathobiont *Streptococcus mutans* Using Ancient DNA"

### SUPPLEMENTARY MATERIAL

#### S1. Archaeological and Anthropological Context for Analyzed Samples

Unless otherwise stated, radiocarbon dates were calibrated using OxCal v4.4 with the IntCal20 calibration curve.

#### I. AFRICA

##### 1.1 Faraoskop

*Country:* South Africa
*Region:* Western Cape
*Coordinates:* -32.1253°, 18.6144°
*Sample Date:* 350 BCE- 50 CE
*Radiocarbon Date (FAR004, bone, Pta-5283):* 2000 ± 50 BP, 68 calBCE - 202 calCE (2-sigma, SHCal13)[1]
*Excavation Details:* Excavations by A. Manhire and colleagues in 1987-1988[1]. The human remains from Faraoskop are stored in the Department of Human Biology at the University of Cape Town in South Africa[2].
*Human Genetic Analysis:* Skoglund *et al.* 2017 (ref. [2])

*Summary:*

Faraoskop consists of a rock shelter situated on an elevated ridge (300 m) approximately 30 km inland from Elands Bay in the Western Cape Province of South Africa[1,3]. The site preserves a rich archaeological assemblage of Later Stone Age artifacts, including stone tools, shell artifacts, ostrich eggshell beads, leather, and twine[2]. A total of 12 individuals were recovered from Faraoskop; 7 were retrieved by a local landowner prior to excavation, while the remaining 5 were unearthed during controlled excavations by A. Manhire and colleagues in 1987-1988[1]. Overlapping radiocarbon dates from 6 individuals suggest that the burials may have been deposited in a single interment event c. 2300 to 1900 years BP[1].

Notably, 4 out of 5 individuals with recovered dentition display carious lesions, a comparatively high rate relative to contemporaneous coastal hunter-gatherer groups. Isotopic analysis suggests that the individuals from Faraoskop relied largely on terrestrial food resources despite their proximity to the coast. This terrestrial diet, which may have included starch-rich tubers, could account for the high prevalence of caries in this hunter-gather group[3].

- **FAR004 (UCT-386, I9133)** is a 40–50-year-old genetically male individual[1,2]. Two out of 30 teeth from this individual display carious lesions, including the tooth analyzed in this study[3,4]. Interestingly, a previous study including individual UCT-386 failed to identify high levels of *S. mutans* DNA in dental calculus, suggesting that the pulp chamber and/or dentine may provide a better source for identification and genomic reconstruction of opportunistic oral pathogens[5].

#### II. AMERICAS

Central America

##### 2.1 Hospital Real de San José de los Naturales

*Country:* Mexico
*Region:* Mexico City
*Coordinates:* 19.4313°, -99.1416°
*Site Occupation:* 1500-1700 CE
*Radiocarbon Date (SJN015, tooth, MAMS-35847):* 554 ± 17 BP, 1326-1423 calCE (2-sigma)
*Excavation Details:* The installation of a new subway line in downtown Mexico City prompted excavation of the Hospital Real de San José de los Naturales between 1988 and 1994. Excavations were led by Salvador Pulido Méndez and María de Jesús Sánchez Vázquez[6].
*Human Genetic Analysis:* Three individuals from the Hospital Real de San José de los Naturales were analyzed in Barquera *et al.* 2020 (ref. [6]), although SJN015 and SJN032 were not included in that study.

*Summary:*

Situated in downtown Mexico City, the Hospital Real de San José de los Naturales was founded between 1529 and 1531 to serve the indigenous inhabitants of the Viceroyalty of New Spain[6], in particular sufferers of the multiple epidemics of smallpox and other infectious diseases that decimated local Indigenous populations in the post contact period. The majority of the hospital's patients were likely Nahuatl and Otomi speakers from the central region of Mexico; however, genetic, isotopic, and osteological analyses identified three individuals as first generation African individuals transported to Central America through the trans-Atlantic slave trade[6]. Excavations uncovered multiple individuals in 13 of 16 excavational units, suggesting the presence of one or more elevated mortality events. Archaeogenetic analyses have succeeded in reconstructing genomes of hepatitis B virus (HBV) and *Treponema pallidum* sub. *pertenue*, the causative agent of yaws, pointing to a high incidence of infectious disease in the individuals interred at the Hospital Real de San José de los Naturales[6]. Two individuals from the hospital are included in the present study:

- **SJN015 (TU258/San José de los Naturales SLU 11B 1992 4)** is a skeleton from a young adult male (30-35 years old), with a gracile complexion. Based on identified enamel hypoplasia that would have formed between 3-5 years of age, it is hypothesised that the individual suffered from vitamin or mineral deficiency or a gastrointestinal condition during childhood.

  The individual has lesions highly compatible with trepanomatosis. All the long bones present periostitis and bone striations along the diaphysis, suggestive of severe inflammation. The fibulas and left tibia (right tibia was missing) are the most affected bones with enlargement in all the bones. The left ulna and radius present mixed lesions including remodeling and destruction of bone, as well as bone enlargement. Periostitis is also present in the tubular bones of the hands.

In the cranium, we observe a small focus of destruction and remodelling of the bone in the frontal bone, which could be indicative of the initiation of *caries sicca*, a specific lesion of infections by treponema. The frontal bone also presents a small bulge. The occipital bone is flattened, giving the cranium a slight brachiocephalic shape. The individual also presents caries in the majority of its teeth, as well as calculus, and the antemortem loss of the first right molar accompanied by alveolar resorption.

- **SJN032 (Caja Calotas SN)**: no more information is available from this individual.

#### 2.2 San Gregorio Atlapulco

*Country:* Mexico
*Region:* Xochimilco, Mexico City
*Coordinates:* 19.2511°, -99.0556°
*Site Occupation:* Colonial (1521-1810 CE)
*Sample Date:* 1410-1641 CE, based on radiocarbon dating of multiple individuals from the site
*Radiocarbon Date:* -
*Excavation Details:* The archaeological site of El Japón, known since the 1960s, was excavated by Raúl Ávila López in 1995 as part of a salvage excavation, due to the construction of the Distrito de Riego San Gregorio in 1993 without the authorization of the INAH. In 1990, a prospection of the surface of the site was conducted in the ejido (communal farmland) of San Gregorio. During the excavation, bones and ceramic materials were rescued. As part of this process, an initial osteological analysis was conducted in the skeletonised individuals, including biological profiling (age and sex) and assessment of pathologies.

*Summary:*

Located in the modern neighborhood of San Gregorio Atlapulco, Xochimilco, Mexico City, El Japón was a site of seasonal occupation in the late postclassic period, with permanent settlements established around the time of European contact[7–10]. The inhabitants of El Japón practiced intensive agriculture using a system of chinampas, raised beds constructed in shallow lakes[9,10]. Excavations in 1993 unearthed the remains of ten habitation platforms as well as 400 primary burials and two ossuaries, which have been dated to the pericontact period (c. 1521 CE)[9,11]. In addition to showing skeletal markers of occupational stress, osteological analysis identified pathological lesions linked to malnutrition and/or infection in the population of El Japón. The site's age-at-death profile is also skewed towards juveniles and young adults, suggesting a population with a high growth rate, high infant mortality, and low life expectancy[9,12,13].

- **SGA024 (San Gregorio U69 93-94 Ent. 275)**: a young male individual (24-28 years old). The skeleton presents cribra orbitalia, porotic hyperostosis, and enamel hypoplasia (formed around 2-3 years of age), pointing towards malnutrition, vitamin deficiencies, or gastrointestinal issues in childhood. Notably, the individual presents trauma likely due to a fall, seen in the coxofemoral articulation that severely affected the head of the femur, leading to the destruction and proliferation of the acetabular tissue. The right lower extremity was also affected, seen by lesions in the diaphysis of the fibula. In general, the individual shows loss of bone density, possibly linked to the reduced mobility caused by the injury. They also present bone reabsorption between the right parietal and occipital bones.

#### South America

##### 2.3 Huaca Pucllana

*Country:* Peru
*Region:* Lima
*Coordinates:* -12.1150°, -77.0335°
*Site Occupation:* 500-1450 CE (ref. [14])
*Sample Date(s):*

XXX001.A161241: 500-700 CE
XXX001.A161237: 800-1000 CE
XXX001.A161233: 1000-1450 CE
XXX001.A161239: 1000-1450 CE

*Radiocarbon Date (XXX001.A161241, OxA-31120):* 1493 ± 30 BP, 549-652 calCE (2-sigma, SHCal13)[14]
*Excavation Details:*
The excavations where the contexts related to this research were recovered were led by Isabel Flores Espinoza and held between 1995 and 2012. Huaca Pucllana was a monumental ceremonial center of the Lima Culture (100-650 CE) and was used as a cemetery during the Wari (550-1100 CE) and Ychsma (1100-1470 CE) periods. Around 150 tombs were excavated from this site.
*Human Genetic Analysis:* Mitochondrial data published in Valverde *et al.* 2016 (ref. [14]). Nuclear data from XXX001.A161241 (I0975), XXX001.A161233 (I0966), and XXX001.A161239 (I0972) published in Nakatsuka *et al.* 2020 (ref. [15]).

*Summary:*

Huaca Pucllana is a Central Andean site located in the Miraflores district of Lima, Peru. The site's history spans almost a millennium, with periods of prehispanic occupation corresponding to the Lima culture (450-650 CE), the Wari (850-1000 CE), and the Ychsma culture (1000-1470 CE)[14]. Huaca Pucllana functioned in the Lima period as a ceremonial center, with monumental architecture for feasting and worship. Mitochondrial analyses of individuals from Huaca Pucllana find no evidence for population turnover associated with the Wari expansion and subsequent transition to the Ychsma culture, suggesting that cultural shifts may have been driven by social and political processes rather than genetic turnover[14]. Similarly, a regional time transect of Central Andean populations documented broad patterns of genetic continuity after ~2000 BP[15].

In chronological order, the following individuals from Huaca Pucllana were included in this analysis:

- **XXX001.A161241 (LP54.12,10817A, I0975)** is an adult individual identified as female based on genetic analyses. Contextually associated with the Early Intermediate Period Lima culture, radiocarbon analysis dates the individual to c. 549-652 calCE.
- **XXX001.A161237 (LP54.7,10773A, I0970)** is an adult of undetermined sex associated with the Middle Horizon Wari culture, which dates from c. 800-1000 CE.
- **XXX001.A161233 (LP54.3, 10725A, I0966)** is an adult genetically male individual associated with the Late Intermediate Period Ychsma culture (c. 900-1470 CE).
- **XXX001.A161239 (LP54.9,10800A, I0972)** is a young adult genetically male individual with an estimated age of c. 17-20 years old. XXX001.A161239 is associated with the Late Intermediate Period Ychsma culture and dated to 900-1470 CE based on archaeological context.

#### 2.4 Pavão 16

*Country:* Brazil
*Region:* Sambaqui, Atlantic Coast
*Coordinates:* -24.6493°, -48.7830°
*Site Occupation:* Riverine shellmound occupation (1700–1200 cal yr BP)[16]
*Radiocarbon Date (PVA001, tooth, MAMS-39012):* 1552 ± 22 BP, 482-637 calCE (2-sigma, SHCal20)[17]
*Excavation Details:* Excavated under the direction of Prof. Paulo A.D. Deblasis - MAEUSP, September 2002, within the context of the research project “Investigações Arqueológicas e Geofísicas dos sambaquis fluviais do vale do Ribeira de Iguape, Estado de São Paulo,” coordinated by Prof. Levy Figuti (MAEUSP) between 2000-2003.
*Human Genetic Analysis:* Ferraz *et al.* 2023 (ref. [17])

*Summary:*

Pavão 16 is located within the Atlantic rainforest in Itaoca county in the Ribeira de Iguape Valley in southeastern São Paulo State, Brazil. It is situated in an urban area at 150 masl. The site contains partially razed riverine *sambaqui* (shell mount) measuring c. 40 cm in depth[17]. Found in clusters throughout the Ribeira de Iguape Valley, these small, rounded mounds are generally less than 2 m high with an average area of c. 1000 m$^2$ (ref. [16]). Riverine *sambaquis* are composed mostly of landsnail shells along with bones from terrestrial fauna, lithic and bone artifacts, and human burials. They are thought to have functioned in funerary rites and associated festivities[16]. *Sambaqui* construction in the Ribeira de Iguape Valley spanned 10,000 - 1,000 yBP and may be linked to the construction of similar coastal shellmounds along 2,000 km of the Brazilian seashore[16]. The *sambaqui* at Pavão contained the remains of three individuals, including the one analyzed here[17].

- **PVA001 (Pavão XVI)** is an individual of undetermined sex dated via radiocarbon analysis to c. 482-637 calCE (2-sigma).

##### 2.5 Pongumal (Pongumál)

*Country:* Peru
*Region:* Conila district, Luya province, Amazonas department
*Coordinates:* -6.1167°, -78°
*Site Occupation:* 1000-1475 CE
*Radiocarbon Date (PGL001, tooth, MAMS-35095):* 545 ± 19 BP, 1405-1445 calCE (2-sigma, SHCal20)
*Excavation Details:* Archaeological material from Pongumal was recovered in 1984 by anthropologists Henry and Paul Reichlen during a survey of various sites along the upper Utcubamba river in Amazonas[18]. The remains discussed here were found among a commingled assemblage of bones representing a minimum number of three individuals[19].

*Summary:*

Culturally affiliated with the Chachapoya, the site is located on the Ayshpachaka ravine, on the left bank of the Luya River. It comprises a series of funerary mausoleums or *chullpas*, situated near the present-day village of Conila. At the time the Reichlens explored the site, remnants of paint were still visible in the façade of the structures[18].

- **PGL001 (CHA052)** is an adult individual with osteologically undetermined sex and genetic sex indicating a female. It is associated with the Late Intermediate Period Chachapoya culture.

##### 2.6 Revash

*Country:* Peru
*Region:* Santo Tomás district, Luya province, Amazonas department
*Coordinates:* -6.3666°, -78.0012°
*Site Occupation:* 1000-1475 CE
*Radiocarbon Date (REV004, tooth, MAMS-35098):* 627 ± 16 BP, 1320-1407 calCE (2-sigma, SHCal20)
*Excavation Details:* Revash was also surveyed by anthropologists Henry and Paul Reichlen in 1948. While exploring nearby cliffs, they discovered a burial cave (Revash 2) containing mummy bundles whose wrappings had already deteriorated leaving only skeletal remains. Part of this osteological material could be safely recovered and transported by the researchers[18].

*Summary:*

Also associated with the Chachapoya culture, Revash consists of funerary and residential sectors, as well as a burial cave originally containing mummy bundles. Located at approximately 1800 masl on the slopes and summit of Cerro Carbón, the site features quadrangular chullpas built into semi-artificial rock shelters on sheer cliffs above the left bank of the Santo Tomás valley. These structures were constructed using small stones and a mortar of clay and straw, then painted with red and beige pigments. Floors and door frames were made of wooden logs tied with ropes[20]. Although the chullpas had been looted prior to the Reichlens' survey, the burial cave appeared to be intact at that time[18].

- **REV004 (CHA066**) is an adult male, confirmed by both osteological and genetic analysis. This individual, associated with the Late Intermediate Period Chachapoya culture, was recovered from the Revash 2 burial cave alongside remains of other individuals.

#### III. EURASIA

##### Anatolia

##### 3.1 Alalakh (Tell Atchana)

*Country:* Türkiye
*Region:* Hatay Province
*Coordinates:* 36.2378°, 36.3847°
*Site Occupation:* 2200-700 BCE
*Radiocarbon Date (ALA002, petrous bone, MAMS-33676):* 3158 ± 22 BP, 1499-1398 calBCE (2-sigma)[21]
*Excavation Details:* Excavations in 1937-1939 and 1946-1949 by Sir Leonard Woolley under the auspices of the Trustees of the British Museum, from 2003-2019 by Kutlu Aslıhan Yener, and from 2020-present by Murat Akar under the auspices of the Turkish Ministry of Culture and Tourism[21].
*Human Genetic Analysis:* Skourtanioti *et al.* 2020 (ref. [21])

*Summary:*

Alalakh (Tell Atchana) is a riverside settlement located in the Amuq Valley in the modern state of Hatay, Türkiye. The site's occupation spans the terminal Early Bronze Age/earliest Middle Bronze Age (c. 2200-2000 BCE) through the Late Bronze Age (c. 1300 BCE), followed by a limited reoccupation during the Iron Age[21]. During the Bronze Age, a succession of political entities exerted influence over Alalakh, including the kingdoms of Yamhad and Mittani, as well as the Hittites[21]. A detailed description of the site's shifting chronology and material culture, as well as human population genetic analysis, can be found in Skourtanioti *et al*. 2020.

Three hundred and forty-five burials have been recovered from Alalakh during both the original excavations by Sir Leonard Woolley (1937-1939 and 1946-1949) and ongoing excavations begun in 2003 under the direction of K. Asıhan Yener, currently directed by Murat Akar; however, skeletal remains from Woolley's project were not preserved. Burials were located both in an extramural cemetery and in various locations within the city walls, including abandoned buildings, courtyards, and an intramural cemetery area. Diverse grave types are present at the site, including several constructed tombs, and the burials differ greatly in the number and type of grave goods present[21].

- **ALA002 (45.71, Locus 03-3017, Pail 246, Skeleton S04-8)** is a young adult, genetically male individual with an estimated age between 19 and 21 years old based on skeletal evidence[21]. This individual was recovered from the Plastered Tomb located in the extramural cemetery in Area 3. Paleopathological analyses identified both *cribra orbitalia* and porotic hyperostosis, and both humeri exhibit a non-metric trait known as Septal Aperture. The individual was buried with six bronze pins, a bone needle, gold appliques and a gold ring, and a necklace and beads of gold, carnelian, and white vitreous material. Radiocarbon dating places the individual between 1499 and 1398 calBCE.

##### 3.2 İkiztepe

Country: Türkiye
*Region:* Bafra, Samsun Province
*Coordinates:* 41.6137°, 35.8711°
*Site Occupation:* 4000-3000 BCE
*Sample Date:* 3500-3000 BCE
*Excavation Details:* Excavation from 1974 to 1981 under the directions of Uluğ Bahadır Alkım. Subsequent excavation began in 1981 led by Önder Bilgi of İstanbul University[22]. Human remains from the site are housed at the Hacettepe University Skeletal Biology Laboratory (Husbio-L)[21].
*Human Genetic Analysis:* Individuals from İkiztepe analyzed in Skourtanioti *et al.* 2020 (ref. [21]), although the individual analyzed here (IKI023) was not included in that study.

*Summary:*

Situated on the Bafra plain in the contemporary Samsum province of Türkiye, İkiztepe comprises four mounds whose occupation spanned the early Chalcolithic through the Hittite period[21,23]. Since 1974, excavations at İkiztepe have proceeded under the direction of Uluğ Bahadır Alkım and later Önder Bilgi[23]. A total of 700 Late Chalcolithic pit graves containing ~766 individuals have been unearthed from an extramural cemetery on Mound I[22]. The majority of individuals from İkiztepe were buried in simple pit burials in supine position, and grave goods included metal objects of an arsenic-copper alloy, gold, and silver[21,23].

Despite its present-day location approximately 7 km from the Black Sea coast and 1.5 km from the Kızılırmak River, researchers infer that İkiztepe was likely sited along both the riverbank and coastline during its principal period of occupation[23]. Nevertheless, analyses of sulfur and nitrogen isotopes support limited reliance on seafood and freshwater resources; instead, carbon isotopic analyses of Chalcolithic and Bronze Age inhabitants support a diet based principally on terrestrial $C_3$ food sources[23]. Similarly, archaeobotanical analyses have identified grains, including emmer wheat, einkorn, and barley, pulses such as bitter vetch, grass pea, and pea, and wild fruits and nuts. Zooarchaeological evidence supports exploitation of both wild and domesticated fauna, with strontium isotope data possibly supporting some level of mobile pastoralism/seasonal transhumance[22].

- **IKI023 (SK 624)** is an adult male with an estimated age between 33-44 years old. Osteological analyses revealed healed fractures of both the left clavicle and right ulna (“parry fracture”), as well as cranial injuries, possibly including both blunt and sharp force trauma. Additional pathologies include nonspecific lower limb infection and a mild osteoma of the right frontal bone[22]. Further details are summarized in Irvine *et al*. 2020 (ref. [23]).

##### 3.3 Titriş Höyük

*Country:* Türkiye
*Region:* Şanlıurfa Province
*Coordinates:* 37.4762°, 38.6761°
*Site Occupation:* 2900-2100 BCE
*Sample Date:* 2300-2100 BCE
*Excavation Details:* Excavations from 1991-1999 led by Guillermo Algaze (University of California San Diego) and Timothy Matney (University of Akron)[21,22].
*Human Genetic Analysis:* Data from Titriş Höyük (excluding individual TIT012 analyzed here) was published in Skourtanioti *et al.* 2020 (ref. [21]).

*Summary:*

Located in the Karababa basin in present-day southeastern Türkiye, Titriş Höyük is a large 43-hectare settlement situated along the banks of the Tavuk Çay, a tributary of the Euphrates River. The site consists of a central acropolis, a surrounding Lower Town, and a dispersed Outer Town with a relatively short occupation period spanning the Early to Late EBA[21,22]. During its peak in the Mid-Late EBA, Titriş Höyük was likely one of the largest settlements in the region. Situated along both riverine and overland transit routes, the cosmopolitan center may have functioned as a regional capital, and archaeological evidence supports the integration of Titriş Höyük in long-distance trade networks connecting various regions of Anatolia, the Aegean, and Mesopotamia[22]. The site experienced a shift beginning in the late EBA, with abandonment of the Outer Town and construction of a fortification wall. This transition also marked a shift in burial practices; Middle EBA inhumations were concentrated in an extramural cemetery situated approximately 400 m from the settlement, giving way to intramural burials associated with households in the Late EBA. Burials consisted of small pits, stone cysts, and pithoi, with multiple burials of articulated and/or disarticulated individuals in both Mid and Late EBA contexts[21].

Archaeobotanical and isotopic studies infer a diet heavily reliant on wheat and barley, with cultivation of other domesticated plants such as legumes and grapes[22].The inhabitants of Titriş Höyük also practiced animal husbandry mainly focused on sheep, goat, and cattle. Dietary isotope studies infer a reliance on a terrestrial $C_3$-based mixed diet, while low $\delta^{15}N$ values reflect a lower protein consumption compared to other sites. There is some evidence for centralized coordination in the production/distribution of food resources, possibly consistent with the relatively limited diversity in plant and animal species consumed at the site[22].

- **TIT012 ('96 TH 63202/1)** is an adult male individual recovered from a cyst grave and contextually dated to the late Early Bronze Age period.

<u>Balkans</u>

#### 3.4 Pietrele

*Country:* Romania
*Region:* Măgura Gorgana, Giurgiu County
*Coordinates:* 44.0681°, 26.1562°
*Site Occupation:* c. 4700-4200 BCE (copper age settlement mound)
*Sample Date:* c. 4700-4200 BCE, Chalcolithic period
*Radiocarbon Date (PIE034, bone, MAMS-47819):* 5741 ± 22 BP, 4681-4502 calBCE (2-sigma)[24]
*Excavation Details:* First excavated in the 1940s by Dumitru Berciu. Excavations since 2002 under the auspices of the German Archaeological Institute in Berlin, the Institute for Archaeology 'Vasile Pârvan' of the Romanian Academy of Sciences in Bucharest, and the Institute for Geography of Goethe University in Frankfurt[24].
*Human Genetic Analysis:* Penske *et al.* 2023 (ref. [24])

*Summary:*

Situated approximately 7 km north of the Danube River on the site of a former paleo-lake, the site of Pietrele consists of a Copper Age tell (c. 4700-4200 BCE) built on the ruins of a former Neolithic settlement[24]. Sitting up to 11 m above the surrounding landscape, the settlement mound rose gradually as destroyed/abandoned structures were repeatedly filled and used as a base for later structures. From c. 4450 BCE, construction expanded to encompass the flat area around the settlement mount. During its peak the site was a center of long-distance trade with evidence of economic specialization in craft production[24]. Pietrele caught fire and was subsequently abandoned in c. 4250 BCE, mirroring the contemporaneous demise of other settlements along the Lower Danube[24].

- **PIE034 (P08 B 108)** is a genetically female individual of c. 15-16 years old exhibiting evidence of blunt force trauma to the cranium. The individual was recovered from the settlement mount in an area of fill outside of the household structures. Radiocarbon analysis of this individual gives a later date than surrounding houses, consistent with the hypothesis that the material may have been transported from the surrounding area to fill old structures in preparation for new construction[24].

#### 3.5 Yunatsite

*Country:* Bulgaria
*Region:* Pazardzhik
*Coordinates:* 42.23°, 24.26°

*Site Occupation:* 5000-2200 BCE
*Sample Date:* Final phase of the Late Chalcolithic in Thrace (Karanovo VI culture, 3rd phase)
*Radiocarbon Date (I0785, MAMS-28135):* 5578±23 BP, 4451-4354 calBCE (2-sigma)
*Excavation Details:* Excavations in 1939 and since 1976 till present, the latter under the auspices of the National Archaeological Institute with Museum, Sofia. Excavation of Early Bronze Age layers between 1978-1990; excavations of Chalcolithic contexts between 1991-1997 by Bulgarian-Russian team, between 2002-2011 by Bulgarian-Greek team and since 2012 by Bulgarian team.
*Human Genetic Analysis:* Mathieson *et al.* 2018 (ref. [25])

*Summary:*

Located on a river terrace in the Thracian Plain, Yunatsite is a tell site measuring c. 12 m high and 100-110 m in diameter. The tell is in fact the fortified part of a larger settlement, occupied in the Chalcolithic and Early Bronze Age. The mound's use period spans the Chalcolithic (5th millennium BCE), Early Bronze Age (3rd millennium BCE), Iron Age (1st millennium BCE), Roman period and, finally, Medieval period, when the site was used as a cemetery. The remains analyzed here derive from the last Chalcolithic settlement (building level), which is separated from the EBA phase by loose soils suggesting a hiatus of about 1000 years. Individual I0785 was recovered from a destroyed building in the latest Chalcolithic level (LC I). It is associated with the end of the Chalcolithic in Thrace, but shows mixed features of the Karanovo VI culture to the east and the Krivodol culture to the west.

- **XXX001.A161155 (I0785, Yunatsite99b)** is an elderly genetically female individual of c. 70 years old.

#### Baltic

##### 3.6 Donkalnis

*Country:* Lithuania
*Region:* Lake Biržulis, Telšiai District
*Coordinates:* 55.8078°, 22.4222°
*Site Occupation:* 6000-2500 BCE
*Sample Date:* 5500-2900 BCE
*Radiocarbon Date (DON004, fibula, Poz-61574):* 5770 ± 40 BP, 4718-4503 calBCE (2-sigma)[26]
*Excavation Details:* Excavated in 1981-1983 by Adomas Butrimas following reports of human remains unearthed during gravel quarrying activities at the site[27].
*Human Genetic Analysis:* Mittnik *et al.* 2018 (ref. [27])

*Summary:*

Situated atop a small knoll on the shores of Lake Biržulis in central north-western Lithuania, the site of Donkalnis consists of a settlement and associated cemetery located on a small former island with a use period spanning the Mesolithic through the Middle Neolithic.

Excavations in 1981-1983 recovered both 7 intact inhumation burials and 6 additional fragmented graves likely disrupted during quarrying activities[27]. The inhumation graves were accompanied by animal teeth pendants evidently used as clothing decorations, pottery fragments and flint artifacts. Dates based on radiocarbon analysis and archeological context indicate that the burials derive from both the Mesolithic and the Middle Neolithic periods[27].

The individual analyzed in this study, DON004, dates from the Subneolithic period and is associated with the Narva culture[27]. The first strong evidence for plant and animal domestication in the Eastern Baltic appears in the subsequent Neolithic period around 3200/2700 calBCE, in association with the Corded Ware and Globular Amphora cultural horizons[28]. Stable isotope analysis suggests that Mesolithic and Subneolithic individuals from Donkalnis and other inland Baltic sites relied on both freshwater fish and forest game, with a likely addition of wild plant material; this contrasts sharply with the diet of subsequent Neolithic groups, which relied heavily on terrestrial domesticates as a protein source[26].

- **DON004 (Donkalnis6)** is an approximately 35-40-year-old genetically female individual associated with the Mesolithic Narva culture and radiocarbon dated to 4718-4503 calBCE[27].

##### 3.7 Kivutkalns

*Country:* Latvia
*Region:* Vidzeme
*Coordinates:* 56.8521°, 24.2720°
*Site Occupation:* 1st millennium BCE[29]
*Radiocarbon Date (KIV002, tooth, Hela-3743):* 2511 ± 30 BP, 787-541 calBCE (2-sigma)[27]
*Radiocarbon Date (KIV006, tooth, Hela-3740):* 2573 ± 30 BP, 809-570 calBCE (2-sigma)[27]
*Radiocarbon Date (KIV007 (1), mandible, Hela-3741):* 2497 ± 30 BP, 778-516 calBCE (2-sigma)[27]
*Radiocarbon Date (KIV007 (2), maxilla, Hela-3742):* 2556 ± 30 BP, 804-551 calBCE (2-sigma)[27]
*Sample Date(s):* 1300-500 BCE (KIV002, KIV006, and KIV007)
*Excavation Details:* Excavations in 1966 and 1967 led by Jānis Graudonis. The site was completely excavated prior to the construction of the Riga Hydroelectric Plant[27].
*Human Genetic Analysis:* Mittnik *et al.* 2018 (ref. [27])

*Summary:*

Situated on the island of Dole in the lower River Daugava, the hillfort and cemetery site of Kivutkalns constitutes the largest known LBA bronze-working center in Latvia[27]. In an arrangement atypical for the region, the Kivutkalns hillfort was constructed on top of the cemetery; radiocarbon analysis dates the hillfort to the 1st millennium BCE, while the cemetery's use period seems to have overlapped that of the hillfort. Jānis Graudonis's excavations of the site in 1966-1967 revealed 247 inhumation burials and 21 cremations with good overall preservation[27].

- **KIV002 (Kivutkalns207)** is an approximately 40-50-year-old genetically female individual from the Late Baltic Bronze Age buried inside an oak log. Grave goods include a bone needle, and white sand was placed beneath the burial.
- **KIV006 (Kivutkalns42)** is an approximately 35-42-year-old genetically female individual associated with the Late Baltic Bronze Age Cist Grave Culture. This individual was interred without grave goods, possibly in a linden coffin.
- **KIV007 (Kivutkalns209)** is an approximately 25-33-year-old genetically male individual associated with the Late Baltic Bronze Age Cist Grave Culture. The individual was buried inside an oak log, with stones at the head and white sand below the burial.

##### 3.8 Plinkaigalis

*Country:* Lithuania
*Region:* Kaunas County
*Coordinates:* 55.4102°, 23.6468°
*Site Occupation:* The majority of burials date from the 3rd to the 8th century CE, while two graves (including Plinkaigalis242) derive from the Late Neolithic (c. 2900-2300 BCE)
*Sample Date:* 2900-2300 BCE
*Radiocarbon Date (PLI001, OxA-5936):* 4280 ± 75 BP, 3264-2625 calBCE (2-sigma)[27]
*Excavation Details:* The site was discovered during gravel digging operations in 1975 and was excavated between 1977-1984 by Vytautas Kazakevičius.
*Human Genetic Analysis:* Mittnik *et al.* 2018 (ref. [27])

*Summary:*

Situated on the river Šušvė near the modern village of Plinkaigalis in central Lithuania, the burial site contains a total of 373 excavated graves, including 364 inhumations and 9 cremation burials. Despite being located only 400 m southeast of an Iron Age settlement and hillfort, 371 of the 373 burials at Plinkaigalis date from the 3rd to the 8th century CE[30]. Two graves from the northern area of the site date to the Late Neolithic period. The inhumation grave was accompanied by 3 flint knives.

- **PLI001 (Plinkaigalis242)** is an adult female individual likely over 40 years of age. PLI001 has been dated to the Neolithic period and is associated with the Corded Ware/Battle Axe archaeological culture.

#### Caucasus

##### 3.9 Ginchi

*Country:* Russian Federation
*Region:* Dagestan

*Coordinates:* 42.4290°, 46.6404°
*Site Occupation:* 2400-1550 BCE
*Radiocarbon Date (GIN005, tooth, MAMS-42349):* 3351 ± 20 BP, 1733-1542 calBCE (2-sigma)[31]
*Radiocarbon Date (GIN007, tooth, MAMS-42364):* 3345 ± 20 BP, 1729-1538 calBCE (2-sigma)[31]
*Excavation Details:* Excavated in 1956-1960 by the Excavation Mountain Expedition of the Dagestan branch AN SSSR under the leadership of M. Gadzhiev[31].
*Human Genetic Analysis:* Ghalichi *et al.* 2024 (ref. [31])

*Summary:*

The Middle Bronze Age cemetery of Ginchi is situated on the right bank of Gichinoor creek near the village of Tidib in the mountainous region of Dagestan. The cemetery was partially eroded prior to excavation in 1956-1960. Work led by M. Gadzhiev unearthed the remains of 15 crypts, two child burials, and 15 burials in ceramic vessels from an area of c. 350 $m^2$. Each grave contained the remains of numerous individuals along with various grave goods[31].

- **GIN005 (11556)** is a genetically female individual recovered from a multiple burial in crypt 3, which contained the remains of 16 individuals interred in four layers separated by soil.
- **GIN007 (11558)** is a genetically male individual also interred in crypt 3.

##### 3.10 Marinskaya 5

*Country:* Russian Federation
*Region:* Stavropolskiy kray
*Coordinates:* 43.9054°, 43.5219°
*Site Occupation:* The site was in use with interruptions from the early 4th millennium BCE to the 1st millennium CE.
*Radiocarbon Date (MK5008, tooth,* MAMS-29810*):* 4544 ± 25 BP, 3369-3103 calBCE (2-sigma)[32]
*Excavation Details:* Excavations in 2009 under the leadership of Lomonosov Moscow State University, the Institute of Archaeology RAS, and the local heritage organization 'Nasledie'[32].
*Human Genetic Analysis:* Wang *et al.* 2019 (ref. [32])

*Summary:*

Located north of the Caucasus mountains in the herb-grass Steppe ecozone, the site of Maryinskaya 5 consists of a single multi-phase burial kurgan on a terrace overlooking the river Kura[32]. Encircled by a 1.5 m deep ditch, the massive oval-shaped mound measures c. 34-40 m across and 4.3 m high. In 2009, excavations led by Lomonosov Moscow State University, the Institute of Archaeology RAS, and the local heritage organization 'Nasledie' uncovered four mound-shell construction phases composed of earth and stone[32]. A total of 34 burials and a ritual complex dating back to the Bronze and Iron Ages were discovered in the site. The earliest group of burials was made during the Early Bronze Age and belonged to the Maikop culture[33].

Three mound-shells dated to the Early Bronze Age and contained two Early and three Late Maykop-associated graves. Following a hiatus of c. 600 years, a fourth shell mound with additional burials was added in the Middle Bronze Age[32]. The site's use period continued through the Iron Age Sarmatian period, with the latest burials dating to the 3rd to the 1st century CE[32]. Paleoproteomic analysis of dental calculus from Early/Late Maykop individuals from Maryinskaya 5 identified sheep milk proteins, consistent with the pastoral lifestyle and reliance on *Ovis* dairying observed in other Maykop individuals from the North Caucasus[34].

- **MK5008 (BZNK-066/2, 066/3)** is a juvenile genetically male individual recovered from grave 16 in the center of kurgan 1. Associated with the Late Maykop cultural horizon, radiocarbon dating places the individual in the Early Bronze Age (3369- 3103 calBCE).

#### 3.11 Nogir 3

*Country:* Russian Federation
*Region:* North Ossetia- Alania
*Coordinates:* 43.0844°, 44.6327°
*Site Occupation:* 3800-3600 BCE
*Sample Date:* Early Maykop
*Radiocarbon Date (OSS001, tooth, MAMS-29813):* 4857 ± 23 BP, 3704-3535 calBCE (2-sigma)[32]
*Excavation Details:* Rescue excavations conducted in 2016.
*Human Genetic Analysis:* Wang *et al.* 2019 (ref. [32])

*Summary:*
Nogir is situated in the Caucasus forest steppe ecozone in the modern town of Vladikavkaz. After incurring damage during road construction, mount 3 was unearthed during 2016 rescue excavations[32].

- **OSS001 (kurgan 3, grave 4, ind. 1)** is a genetically female individual buried in the Maykop cultural tradition. Radiocarbon analysis dates this individual to 3704-3535 calBCE.

#### 3.12 Velikent

*Country:* Russian Federation
*Region:* Dagestan
*Coordinates:* 42.1798°, 48.0661°
*Site Occupation:* The Velikent archaeological complex dates from the first half of the 3rd millennium BCE to the beginning of the 2nd millennium BCE[35]
*Sample Date:* 3000-2800 BCE
*Excavation Details:* Excavations by a Russian-American team in 1994 and 1997[32].

*Human Genetic Analysis:* Wang *et al.* 2019 (ref. [32])

*Summary:*

Situated on the coast of the Caspian Sea, the site of Velikent spans over 30 ha, including several settlements and burial mounds. The inhabitants of the site were agro-pastoralists associated with a northern variant of the Kura-Araxes culture. Excavations in 1994-1995 uncovered the partially destroyed remains of a collective catacomb grave, which included bronze artifacts and skeletal remains constituting a minimum number of seven individuals[32].

- **VEK006 (BZNK-486/6;8)** is a genetically female individual associated with the Kura-Araxes culture and contextually dated to c. 3000-2800 BCE.

#### Central Europe

##### 3.13 Alsónyék-Bátaszék, Mérnökségi telep

*Country:* Hungary
*Region:* South-east Transdanubia*,* Tolna County
*Coordinates:* 46.2086°, 18.7000°
*Site Occupation:* From Early to Late Neolithic, c. 5750 BCE to c. 4300 BCE
*Radiocarbon Date (XXX001.A16566, MAMS-11939, right ulna)*: 6695 ± 40 BP, 5712-5531 calBCE (2-sigma) (ref. [36], Tab. 1, p. 121)
*Excavation Details:* Excavated from 2006-2009 by A. Osztás and I. Zalai-Gaál[37]
*Human Genetic Analysis:* Haak *et al.* 2015 (ref. [38]), Mathieson *et al.* 2015 (ref. [39])

*Summary:*

The Early to Late Neolithic site of Alsónyék-Bátaszék is situated in south-east Transdanubia, on the border between the hilly landscape of the Szekszárd Hills and the former river beds of the Danube river, and also in the cultural contact zone between the Neolithic populations of the Northern Balkans and Central Europe. The site, which covers an area of approximately 80 hectares, was inhabited almost continuously from the Early Neolithic to the end of the Late Neolithic (Starčevo, Linearbandkeramik (LBK), Sopot and Lengyel periods), c. 5750 cal BCE to c. 4300 cal BCE. The occupation reached its greatest extent during the Lengyel period- thousands of features including c. 2300 burials and 122 surface-level, timber-framed houses were discovered. Mérnökségi telep subsite forms the southeastern part of this large site and it was investigated in 2008-2009. The excavation on this subsite is significant due to the discovery of a large Starčevo occupation with numerous settlement features and 25 burials, which constitutes the first manifestation of the Neolithic in Central Europe[37]. This early occupation began at 5775-5740 calBCE (68 % probability), lasted for 190-245 years (68 % probability), and probably ended in 5560–5525 calBCE (68% probability)[36].

- **XXX001.A16566 (BAM25a, I0174)** is a 20-30-year-old genetically male individual associated with the Early Neolithic Starčevo culture[38–40].

##### 3.14 Bottendorf

*Country:* Germany
*Region:* Roßleben, Kyffhäuserkreis, Sachsen-Anhalt
*Coordinates:* 51.3053°, 11.4036°
*Radiocarbon Date (BOT005, OxA-27248):* 6685 ± 38, 5701-5486 calBCE (2-sigma)[41]
*Excavation Details:* After its discovery during sand extraction, the site was excavated in 1939 by the local community in collaboration with experts.[41]
*Human Genetic Analysis:* Rivollat *et al.* 2020 (ref. [41])

*Summary:*

The site of Bottendorf is located c. 45 km southwest of Halle in the Kyffhäuserkreis district of Sachsen-Anhalt, Germany. The site contains two single and one double burial from the Mesolithic period situated approximately 15 meters apart, as well as an Early Bronze Age burial ground[41]. The individual analyzed here was recovered from burial II, which contained the remains of an adult female and a child of c. 5-7 years old. The burial was partially disturbed and no grave goods were recovered, although the skeletal remains were colored with red ochre[41]. Dietary isotope analysis of the individuals is consistent with an opportunistic subsistence strategy primarily reliant on terrestrial C3 resources[41,42].

- **BOT005 (HK39:147b BII/2)** is a genetically female individual dating to the Mesolithic period.

##### 3.15 Esperstedt

*Country:* Germany
*Region:* Saalekreis, Sachsen-Anhalt
*Coordinates:* 51.4235°, 11.6736°
*Site Occupation:* Esperstedt is situated in the district Saalekreis in the Mittelelbe-Saale region of Sachsen-Anhalt, Germany, c. 21 km west-southwest of Halle (Saale).
*Sample Date:* 2800-2200 BCE
*Radiocarbon Date (I1538, MAMS-34272):* 4008 ± 22 BP, 2574-2470 calBCE (2-sigma)
*Excavation Details:* Rescue excavations were initiated in 2005 as part of a large-scale project associated with construction of the A38 motorway.
*Human Genetic Analysis:* Mathieson *et al.* 2015 (ref. [39])

*Summary:*

Esperstedt consists of a residential and funerary site with a long use period, including phases associated with different Young Neolithic cultures, the Late Neolithic Corded Ware and

Bell Beaker cultures, and the Early and Late Bronze age cultures. Individuals recovered from the site of Esperstedt were associated with the Corded Ware culture[43] based on both burial orientation and ceramics[39].

- **XXX001.A161330 (ESP20, I1538)** is a genetically male individual recovered from archaeological feature 2200[39].The result of the anthropological analysis is Infans II (ref. [44], pg. 297).

##### 3.16 Eulau

*Country:* Germany
*Region:* Naumburg, Sachsen-Anhalt
*Coordinates:* 51.1787°, 11.8538°
*Site Occupation:* Occupational phases associated with both the Corded Ware and Únětice cultures
*Radiocarbon Date (XXX001.A161173, MAMS-22822):* 3650 ± 26 BP, 2135-1941 calBCE (2-sigma)
*Human Genetic Analysis:* Haak *et al.* 2015 (ref. [38]), Mathieson *et al.* 2015 (ref. [39])

*Summary:*
Located in Sachsen-Anhalt, Germany, Eulau sits on a loess-gravel promontory above the Salle river.

- **XXX001.A161173 (EUL41A, I0803)** is a genetically female individual associated with the Early Bronze Age Únětice culture. This individual was recovered from feature 882.

##### 3.17 Halberstadt-Sonntagsfeld

*Country:* Germany
*Region:* Harz, Sachsen-Anhalt
*Coordinates:* 51.8939°, 11.0523°
*Sample Date (HAL05a):* 5500-4850 BCE, based on archaeological context
*Sample Date (HAL14):* 5500-4850 BCE, based on archaeological context
*Radiocarbon Date (HAL05a, MAMS-21479):* 6136 ± 34 BP, 5211-4957 calBCE (2-sigma)[38]
*Radiocarbon Date (HAL14, MAMS-21480):* 6156 ± 35 BP, 5211-5002 calBCE (2-sigma)[38]
*Radiocarbon Date (HAL39b, KIA-40343):* 6144 ± 32 BP, 5210-4998 calBCE (2-sigma)[40,45]
*Radiocarbon Date (HAL36b, MAMS-21484):* 2889 ± 30 BP, 1202-940 calBCE (2-sigma)[38]
*Excavation Details:* The site of Halberstadt-Sonntagsfeld was excavated between 1999 and 2002 following its discovery during construction work. A total of 1324 archaeological features were identified in an area spanning 9947 $m^2$ (ref. [38]).
*Human Genetic Analysis:* Haak *et al.* 2015 (ref. [38]), Mathieson *et al.* 2015 (ref. [39]), Lipson *et al.* 2017 (ref. [40])

*Summary:*

Situated in the Mittelelbe-Saale region of Sachsen-Anhalt, Germany, Halberstadt-Sonntagsfeld consists of a residential and funerary site with a use period spanning the Neolithic through the Bronze Age. The site preserves a rich Linienbandkeramik (LBK) archaeological assemblage, including 42 graves and seven long houses, while additional finds are associated with the Middle Neolithic Bernburg culture and the Bronze age Únětice and Urnfield cultural complexes. Most LBK individuals, including I0046 and I0056 analyzed here, were interred in graves surrounding the LBK long houses. Two other graves, including individual I0099, were originally classified as LBK, while radiocarbon dating and genetic analysis suggest that they date from the Late Bronze Age period[38].

- **XXX001.A161358 (HAL05a, I0046)** is a genetically female Early Neolithic LBK individual interred in grave 2 (feature 241.1).
- **XXX001.A161364 (HAL14, I0056)** is a genetically male Early Neolithic LBK individual interred in grave 15 (feature 430).
- **XXX001.A161183 (HAL39b, I2037)** is a genetically male Early Neolithic LBK individual.
- **XXX001.A161384 (HAL36b, I2033/I0099)** is a genetically male individual recovered from grave 40 (feature 1114). Originally associated with the LBK culture based on the presence of characteristic ceramics above the grave, radiocarbon analysis places this individual in the Late Bronze Age period. Chronologically, this date associates the individual with the Mittelelbe-Saale Urnfield culture.

##### 3.18 Karsdorf

*Country:* Germany
*Region:* Burgenlandkreis, Sachsen-Anhalt
*Coordinates:* 51.2831°, 11.65°
*Sample Date (KAR11B):* 5500-4850 BCE, based on archaeological context
*Radiocarbon Date (KAR22A, MAMS-23344):* 3993 ± 21, 2572-2467 calBCE (2-sigma)
*Human Genetic Analysis:* Sample KAR22A published in Haak *et al.* 2015 (ref. [38]) and Mathieson *et al.* 2015 (ref. [39]).

*Summary:*

Situated in the Unstrut valley of Sachsen-Anhalt, Germany, the c. 50 acre site of Karsdorf consists of both a settlement and associated funerary features. The site's use period began with the earliest presence of the LBK culture in the region. More than 30 Neolithic houses have been excavated so far, with most graves consisting of pits situated in the center of the settlement area[38].

- **XXX001.A161110 (KAR22A, I0550)** is a genetically female individual recovered from feature 00191 at Karsdorf. Although originally attributed to the Baalberge culture based on archaeological context, genetic and radiocarbon analyses support the association of this individual with the Late Neolithic Corded Ware culture[38].

- **XXX001.A161166 (KAR11B, I0796)** is an individual of unidentified sex contextually associated with the Early Neolithic LBK culture. Previous work on this individual failed to yield sufficient data for human genetic analysis[38].

#### 3.19 Mikulovice

*Country:* Czech Republic
*Region:* Eastern Bohemia, Pardubice District
*Coordinates:* 49.9922°, 15.7767°
*Site Occupation:* 2200-1700 BCE (radiocarbon dating, archaeological context)
*Sample Date:* 1884-1749 BCE, Classical Únětice Culture (archaeological context and radiocarbon dating)[46]
*Radiocarbon Date (MIB002, rib, MAMS-30482):* 3494 ± 19 BP, 1884-1749 calBCE (2-sigma)[46,47]
*Excavation/Sample Provenance:* Excavated in 2009 by R. Sedláček.
*Human Genetic Analysis:* Papac *et al.* 2021 (ref. [46])

*Summary:*

The site of Mikulovice consists of a large, Early Bronze Age settlement with approximately 100 associated burials[46,47]. A complete analysis of the cemetery at Mikulovice has been published elsewhere[47].

- **MIB002 (Grave 55)** is c. 20-25-year-old genetically female individual interred in a right-sided crouched position and facing south. The burial includes a rich assemblage of grave goods, including an awl made of animal bone, two vessels including a classical únětice cup, a bronze pin, bronze earrings, and a necklace made of amber beads and bronze spirals[46].

#### 3.20 Königsbrunn - Obere Kreuzstraße

*Country:* Germany
*Region:* Augsburg, Bavaria
*Coordinates:* 48.2666°, 10.8786°
*Site Occupation:* 2150-1750 BCE
*Sample Date:* 2150-1950 BCE, Early Bronze Age (radiocarbon dating and archaeological context)
*Radiocarbon Date (OBK010, rib, MAMS-18901):* 3664 ± 24, 2136-1955 calBCE (2-sigma)[48,49]
*Excavation Details:* Excavations in 2007 associated with a construction project[49].
*Human Genetic Analysis:* Mittnik *et al.* 2019 (ref. [48])

*Summary:*

The Early Bronze Age site of Königsbrunn - Obere Kreuzstraße is situated in the Lech Valley in the southern German region of Bavaria. Excavations of the cemetery (c. 110 by 18 meters) yielded 50 inhumation burials in 48 grave pits[49]. Individuals were buried in a crouched body position and gender-specific orientation characteristic of the Bell Beaker/Early Bronze Age period. Grave goods recovered included copper, bone, and antler artifacts, bracelets and neck rings, bone/metal pins, and two daggers[49].

- **OBK010 (Grab 23, Bef. 80, OBKR_80)** is an approximately 30-40-year-old genetically male individual dating to the Early Bronze Age equipped with a copper pin and dagger.

#### 3.21 Oberottmarshausen

*Country:* Germany
*Region:* Augsburg, Bavaria
*Coordinates:* 48.2252°, 10.8498°
*Site Occupation:* 1700-1300 BCE
*Sample Date:* 1700-1500 BCE, Middle Bronze Age (radiocarbon dating and archaeological context)
*Radiocarbon Date (OOH016, femur, MAMS-21552):* 3297 ± 36 BP, 1672-1461 calBCE (2-sigma)[48,49]
*Excavation Details:* The site was almost completely excavated in 2004 in association with gravel mining operations[49].
*Human Genetic Analysis:* Mittnik *et al.* 2019 (ref. [48])

*Summary:*

Situated in the Lech Valley in Bavaria, the Middle Bronze Age cemetery of Oberottmarshausen contained 32 graves split into two burial groups[48]. A total of 35 individuals were interred in the cemetery in a stretched supine body position without gender-specific orientation. Both male and female burials lacked rich grave goods, although some bronze pins, pottery, and one dagger were recovered[48].

- **OOH016 (Bef. 141, OTTM_141_d)** is an approximately 30-50-year-old genetically female individual interred in a triple burial and equipped with two bronze pins and a ceramic vessel.

#### 3.22 Prague-Ruzyně

*Country:* Czech Republic
*Region:* Prague, Central Bohemia
*Coordinates:* 50.0908°, 14.3137°

*Sample Date:* 2550-2200 BCE, Chalcolithic/Eneolithic/Final Neolithic/Bell Beaker (archaeological context)
*Radiocarbon Date (PRU003, human bone, MAMS-30801):* 3957 ± 24 BP, 2570-2348 calBCE (2-sigma)[46]
*Excavation Details:* J. Vávra and P. Zelená led rescue excavations at the site in 2011-2012 in association with a construction project.
*Human Genetic Analysis:* Papac *et al.* 2021 (ref. [46])

*Summary:*

Situated in central Bohemia, the site of Prague-Ruzyně contained settlements dating to the Eneolithic, Bronze Age, and Early Iron Age periods, as well as inhumations associated with the Bell Beaker and La Tène cultures[46].

- **PRU003 (Grave 3)** is a c. 50-60-year-old genetically female individual associated with the Bell Beaker culture. Buried in a right-sided crouched position with the head facing south, the individual was accompanied by a bowl and two cups[46].

#### 3.23 Quedlinburg

*Country:* Germany
*Region:* Harz, Sachsen-Anhalt
*Coordinates:* 51.7868°, 11.1535°
*Sample Date (QLB2A):* 3950-3400 BCE, Middle Neolithic (archaeological context)
*Radiocarbon Date(QLB28b, MAMS-22820):* 3824 ± 25, 2433-2147 calBCE (2-sigma)
*Human Genetic Analysis:* Individual QLB28b published in Haak *et al.* 2015 (ref. [38]) and Mathieson *et al.* 2015 (ref. [39]).

*Summary:*

Situated in the northern Harz in present-day Germany, the site of Quedlinburg exhibits a long use period, including phases associated with the Baalberge culture, the Corded Ware and Bell Beaker cultures, and the Únětice culture[38].

- **XXX001.A161176 (QLB28b, I0806)** is a genetically male individual over c. 50 years of age. Buried in an extreme flexed position, the individual is one of a cluster of six attributed to the Bell Beaker culture based on archaeological context[38].
- **XXX001.A161116 (QLB2A, I0556)** is associated with the Middle Neolithic Baalberge culture, but this individual failed to yield sufficient data for human genetic analysis[38].

##### 3.24 Rothenschirmbach

*Country:* Germany
*Region:* Merseburg-Querfurt
*Coordinates:* 51.45°, 11.54°
*Sample Date:* Bell Beaker
*Radiocarbon Date (I0111, Er-8712):* 3881 ± 50 BP, 2474-2201 calBCE (2-sigma)[45]
*Human Genetic Analysis:* Haak *et al.* 2015 (ref. [38]), Mathieson *et al.* 2015 (ref. [39])

*Summary:*
Located on the loess plain of the 'Querfurter Platte', the site of Rothenschirmbach includes a Bell Beaker period cemetery with 14 burials[38]. The cemetery is noteworthy for the presence of structurally embellished graves, including menhirs, cists, and other architectural features, as well as for the oldest documented gold find in central Germany. Anthropological analysis suggests that children are overrepresented in the burial assemblage[38].

- **XXX001.A161203 (ROT4, I0111)** is a genetically female individual associated with the Bell Beaker culture. The burial included the remains of a wooden cist and an overlying 400 kg menhir stone (c. 1.5 x 0.55 m).

##### 3.25 Salzmünde

*Country:* Germany
*Region:* Saalekreis, Sachsen-Anhalt
*Coordinates:* 51.5254°, 11.8283°
*Sample Date:* 3400-3025 BCE, archaeological context
*Radiocarbon Date (I0554, Er-10329):* 4538 ± 47 BP, 3487-3043 calBCE (2-sigma)[45]
*Excavation Details:* Excavated between 2005-2008 by the State Office for Heritage management and Archaeology Saxony-Anhalt, Halle, Germany[45].
*Human Genetic Analysis:* Mitochondrial data in Brandt *et al.* 2013 (ref. [45])

*Summary:*
Located c. 7 km south of Halle/Saale in Saxony-Anhalt, Germany, the site of Salzmünde was inhabited as early as the Paleolithic period. Inhabitation intensified during the Neolithic; excavations of the site in 2005-2008 revealed a causewayed enclosure with two ditches associated with the eponymous Middle Neolithic Salzmünde culture. In addition to evidence of permanent habitation, the site contained burials of 141 individuals[39].

- **I0554 (SALZ88A):** No more information is available from this individual.

##### 3.26 Vliněves

*Country:* Czech Republic
*Region:* Mělník district, central Bohemia
*Coordinates:* 50.3729°, 14.4486°
*Site Occupation:* Large prehistoric polycultural site excavated in the large sand-pic (ca 80 ha). The most important is the Early Bronze Age phase of the occupation with hundreds of burials, many houses, hundreds of sunken features and thousands of other features. Important are also the inhumation cemeteries of Corded Ware Culture, Bell Beaker Culture, Iron Age, Migration Period, and Early Medieval Period and few pre-CW burials. Excavated cemeteries and burial finds were completely published elsewhere[50–53].
*Sample Date (VLI009):* 2900-2600 BCE, Chalcolithic/Eneolithic/Final Neolithic/Corded Ware (archaeological context)
*Sample Date (VLI061):* 2200-2000 BCE, Bronze Age (archaeological context)
*Radiocarbon Date (VLI009, tooth, MAMS-45797):* 4078 ± 25 BP, 2848-2495 calBCE (2-sigma)[46]
*Excavation Details:* Rescue excavation by P. Limburský, Ž. Brnić, V. Salač and I. Pleinerováin 1999-2008.
*Human Genetic Analysis:* Papac *et al.* 2021 (ref. [46])

*Summary:*

The large central Bohemian site of Vliněves experienced a long occupation spanning the Eneolithic through the Early Middle Ages. Thirty hectares (ha) of the 70 ha excavated area yielded settlement and funerary features, including a cemetery associated with the Corded Ware culture and containing 75 inhumation graves and 304 inhumation burials associated with the Únětice culture[46]. Other burials were associated with the Eneolithic Jordanów culture, the Late Michelsberg/Early Baalberge culture, the Globular Amphora culture, the Bell Beaker culture, and the Migration Period[46].

- **VLI009 (Grave 965)** is an adult female of approximately 40-60 years of age associated with the Eneolithic Corded Ware culture. The individual was interred in a left-sided crouched position with the head towards the southeast, and grave goods included a chipped industry blade and intrusive pebble artifacts[46].
- **VLI061 (Grave 517)** is an adult genetically female individual likely over the age of 40 years. Associated with the Bronze Age Proto-Únětice/Early Únětice culture, the individual was buried in a right-sided crouched burial with the head towards the southwest and accompanied by pottery vessels (incl. proto-únětice jug) and animal teeth[46]. EBA finds completely published in Limburský *et al.* 2018 (ref. [53]).

#### Central/Eastern Mediterranean

##### 3.27 Chania

*Country:* Greece
*Region:* Odos Palama Cemetery, Crete
*Coordinates:* 35.5162°, 24.0188°
*Sample Date:* 1350-1250 BCE[54]
*Excavation Details:* Excavations under the auspices of the 25$^{th}$ Ephorate of Prehistoric and Classical Antiquities (until 2014) and the Ephorate of Khania Antiquities (2015-present). Operations at Palama Street co-directed by Maria Andreadaki-Vlazaki and Elpida Hadijidaki. Anthropological assessment by Tina McGeorge[55].
*Human Genetic Analysis:* Individuals from Chania (excluding XAN005) are published in Skourtanioti *et al.* 2023 (ref. [55]).

*Summary:*

Situated on the northern shore of Crete, the city of Chania was founded in the 4th millennium BCE on Kastelli hill, which overlooks a natural harbor[55]. A Minoan Palace was later erected on the site; following its destruction in 1450 BCE, a new palace was constructed by peoples writing in Linear B (LM IIIA1/A2, c. 1370-1350 BCE)[55]. In the Final Palatial period (c. 14th-13th c. BCE) the site appears to have served as a Mycenaean fortified palatial complex. The city of Chania functioned as a multicultural commercial center, with links both to the Greek mainland and the broader Mediterranean sphere[55].

The individual analyzed here was interred in a large (c. 2 $km^2$) cemetery founded in the LMII-LMIII period. Situated roughly 800 m from the Bronze Age hilltop settlement of Chania, the Palama Street burials include 17 rock cut tombs in diverse styles (pit caves, chamber tombs, and a simple pit grave), all of which date to the LM IIIA2-LM IIIB1 period (c. 1350-1250 BCE)[54,55]. The tombs include 29 undisturbed inhumation burials with modest grave goods, including locally made pottery vessels and three silver signet rings of a non-local style[55].

- **XAN005 (Tomb 11)** is an approximately 30-40-year-old female individual based on anthropological assessment.

##### 3.28 Su Cannisoni

*Country:* Italy
*Region:* Seulo, Sardinia
*Coordinates:* 39.388°, 8.3793°
*Site Occupation:* 1600-1400 BCE, Middle Bronze Age (Nuragic I)[56]
*Sample Date:* 1600-1500 BCE, Middle Bronze Age (Nuragic I)[56]
*Radiocarbon Date (SCA003(115), MAMS-39530):* 3281 ± 28 BP, 1619-1498 calBCE (2-sigma)
*Excavation Details:* Excavations in 2009 led by Robin Skeates.

*Human Genetic Analysis:* Marcus *et al.* 2020 (ref. [57]), mitochondrial data in Olivieri *et al.* 2017 (ref. [58])

*Summary:*

Riparo sotto roccia Su Cannisoni 1 consists of a large rock-shelter located in the territory of Seulo in the South Sardinia province, Italy. Dated to the Middle Bronze Age, the site contains a secondary burial deposition covered by a cairn and situated below a natural spring[57].

- **SCA003 (115),** is the 2nd lower premolar of a genetically female individual, assignable to the Middle Bronze Age Nuragic I culture. It is genetically identical to the skull fragment of a genetically female individual from the same archaeological deposit: MA82/SC1004(117)/SCA002, radiocarbon dated to 1532-1429 calBC (OxA-22194)[57].

#### 3.29 Terme di Venosa

*Country:* Italy
*Region:* Potenza, Basilicata
*Coordinates:* 40.9607°, 15.8151°
*Site Occupation:* 300 BCE- c. 1000 CE[59]
*Sample Date:* 672-800 CE (based on radiocarbon analysis of archaeological layer, 2-sigma, IntCal13)[59]
*Excavation Details:* Excavated in the late 1980s.
*Human Genetic Analysis:* Posth *et al.* 2021 (ref. [59])

*Summary:*

Situated on the Appian Way, Venosa was a significant urban center occupied from the 3rd century BCE to around the 10th century CE[59]. The city experienced a turbulent history in the early Middle Ages, falling alternately under the influence of the Byzantine Empire and the Duchy/Principality of Benevento, a semi-autonomous polity with ties to the Longobard Kingdom[59]. Excavations within the city walls uncovered five adjacent multiple burials from this period (c. 8$^{th}$-10$^{th}$ century CE), each containing between 7-12 individuals. The remains lack skeletal indicators of trauma or chronic infection; instead, both demographic evidence and references to local disease outbreaks in historical sources support the hypothesis that the burials may derive from an epidemic mortality event[60].

- **VEN018 (fossa VI.C)** is a genetically female individual from one of the multiple burials at Venosa.

#### 3.30 Vetulonia (Necropolis of San Germano)

*Country:* Italy
*Region:* San Germano, Tuscany

*Coordinates:* 42.9252°, 10.9091°
*Site Occupation:* 801-200 BCE (radiocarbon analysis)
*Radiocarbon Date (VET009,* tooth*, MAMS-42833):* 2197 ± 20 BP, 361-175 calBCE (2-sigma)[59]
*Excavation Details:* Archaeological survey and short excavation campaigns in the 1960s led by Claudio Curri. Agostino Dani undertook site survey in 1967, and both Dani and the Soprintendenza per i Beni Archeologici della Toscana headed excavation campaigns in the 1970s. Ongoing work at the site is led by the S.A.G.A.S Department at the University of Florence and the Soprintendenza per i Beni Archeologici della Toscana.
*Human Genetic Analysis:* Posth *et al.* 2021 (Sample ID VET006.9, petrous and tooth from same individual)[59]

*Summary:*

Situated near the Etruscan city of Vetulonia, in the Sovata Valley in central Italy, the necropolis of San Germano has a period of use ranging from the 7th to the 3rd century BCE, with a brief pause in the 5th century BCE[59]. The necropolis was probably founded by noble Etruscan families, possibly from the nearby town of Vetulonia. The burials were accompanied by grave goods containing bronze, bucchero, black and red-figure Attic pottery[59]. After the site was abandoned in the 5th century BCE, some tumuli were reused from the 4th century BCE. The chronology is confirmed by the grave goods of the new burials, with kylikes, skyphoi and black glazed oinochoai[59].

- **VET009 (T8 n.666)** is a genetically female individual from the Etruscan period.

#### 3.31 Villamar

*Country:* Italy
*Region:* Marmilla, Sardinia
*Coordinates:* 39.6167°, 8.95°
*Site Occupation:* 350-180 BCE, Punic period (archaeological context and radiocarbon analysis of human remains)[61]
*Sample Date:* 350-200 BCE (archaeological context and radiocarbon analysis of human remains)
*Radiocarbon Date:* Dates for multiple individuals from the site reported elsewhere[61]
*Excavation Details:* Originally excavated in the 1990s, new excavations began in 2013 (concession Ministry of Culture to the Municipality of Villamar, in collaboration with the University of Sassari).
*Human Genetic Analysis:* Marcus *et al.* 2020 (ref. [57])

*Summary:*

Located in the Marmilla district of south-central Sardinia, the site of Villamar dates back to the prehistoric and protohistoric periods, although the overlap of successive eras up to the contemporary era does not allow us to understand all phases of the settlement's life. Archaeological investigations to date have focused on a large Punic-era cemetery, from which a total of approximately 60 tombs have been excavated[62]. The tombs vary in structure, ranging from simple trenches to elaborate chamber tombs with shaft entryways, and many contain multiple burials. The pottery has dated most tombs to the mid-4th to the early 2nd century BCE, also supported by radiocarbon analyses carried out on some individuals buried in a hypogeal chamber tomb[57,61]; the frequent reuse of the funerary structures makes this context unique, where ancient alterations can complicate the understanding of the original funerary contexts.

- **VIL009 (US327_CR1, tomb 16)** is a genetically female juvenile recovered from tomb 16 and dating to the Iron Age/Punic period. The individual was interred in a multiple burial containing disarticulated remains from a minimum number of 28 individuals, two of whom were cremated. The assemblage is noteworthy for an over representation of sub-adults[57].

#### Iberia

##### 3.32 Arroyal I

*Country:* Spain
*Region:* Cantabria, Valdeprado del Río
*Coordinates:* 42.9098°, -4.0770°
*Radiocarbon Date (Roy4, MAMS-14857):* 3837 ± 25, 2453-2201 calBCE (2-sigma)[63]
*Excavation Details:* Excavated in 2011-2012 under the auspices of the University of Burgos.
*Human Genetic Analysis:* Olalde *et al.* 2018 (ref. [64]), mitochondrial data in Szécsényi-Nagy *et al.* 2017 (ref. [63])

*Summary:*

Located in Cantabria, the site of Arroyal I consists of a large megalithic grave composed of a rectangular chamber (3 x 3.5 m), a corridor (6 m), and an overlying mound. It was used as a collective burial in the Late Neolithic (3300-2900 BCE); following a period of abandonment, the site was reopened and remodeled in the Chalcolithic period. Several burials include ceramics typical of the Bell Beaker culture[63]. For a more detailed description of the site see Ref [63].

- **XXX001.A161129 (Roy4, I0461)** is a genetically female individual recovered from an isolated pit burial inside the burial mound. This individual is associated with the Late Chalcolithic Bell Beaker culture[63,64].

##### 3.33 Camino del Molino

*Country:* Spain
*Region:* Caravaca de la Cruz, Murcia
*Coordinates:* 38.1028°, -1.8654°

*Radiocarbon Date (Cmol123, tooth, Beta-261529):* 4210 ± 40 BP, 2905-2636 calBCE (2-sigma)[63]
*Excavation Details:* Excavated in 2008 under the auspices of the University of Murcia.
*Human Genetic Analysis:* Olalde *et al.* 2018 (ref. [64]), mitochondrial data in Szécsényi-Nagy *et al.* 2017 (ref. [63])

*Summary:*

Located in the province of Murcia, Spain, the site of Camino del Molino consists of a large multiple burial containing the remains of c. 1363 individuals[63]. The burial consists of a circular pit of 7 m in diameter and 1.6 m deep carved into the rock. While 182 individuals remain in complete or partial anatomical position, most have been disturbed by extensive post depositional movement. Anthropological assessment indicates a roughly equal sex ratio and a preponderance of young individuals (47.5%) in the skeletal assemblage[63]. Accompanying artifacts include copper and flint objects, pottery vessels, and the remains of 50 canids. The site's use period corresponds to the Middle and Final Copper Age, and the burial appears to be associated with the settlement of Molinos de Papel, located 500 m from the burial site[63].

- **XXX001.A161123 (Cmol123, I0455)** is an Early Copper Age genetically male individual[63,64].

##### 3.34 Concheiro do Moita do Sebastião

*Country:* Portugal
*Region:* Muge, Santarem
*Coordinates:* 39.1020°, -8.709°
*Site Occupation:* 6000-5300 BCE, Mesolithic period
*Sample Date:* 6000-5300 BCE, Mesolithic period
*Excavation Details:* The site was identified in 1864 by Carlos Ribeiro of the Portuguese Geological Commission. Multiple excavation campaigns have been undertaken since then, including a project in the 1950s led by the French archaeologist Jean Roche[63].
*Human Genetic Analysis:* One individual from Moita do Sebastao (excluding CMS002) published in Villalba-Mouco *et al.* 2019 (ref. [65]).

*Summary:*

Situated on the Atlantic coast of Portugal, the site of Moita do Sebastao is one of the largest in a series of Late Mesolithic shell middens in the Muge region. The site contains evidence of permanent occupation, including fire places, pits, and post-holes associated with the construction of huts, as well as approximately 100 burials[63,65]. Stable isotope analysis suggests that the Late Mesolithic population represented at Moita do Sebastao consumed a mixed diet consisting of both marine and terrestrial resources[66]. Interestingly, previous analyses have documented a relatively high rate of dental caries at Moita do Sebastao, which may be attributable to the presence of cariogenic fruits in the diet[66].

- **CMS002 (54)** is an individual of unknown age and sex associated with the Late Mesolithic period.

##### 3.35 Concheiro do Cabeço da Arruda

*Country:* Portugal
*Region:* Muge, Santarem
*Coordinates:* 39.1020°, -8.7099°
*Site Occupation:* 6000-5300 BCE, Mesolithic period
*Sample Date:* 6000-5300 BCE, Mesolithic period
*Radiocarbon Date (CCA002, bone, TO-359):* 6960 ± 70 BP, 5986-5721 calBCE (2-sigma)[66]
*Excavation Details:* Multiple excavation campaigns in the 19th and 20th centuries. A detailed description of excavations is given in ref. [67].

*Summary:*

The site of Cabeço da Arruda consists of a Mesolithic shell midden located in the Muge region on Portugal's Atlantic coast. During the Mesolithic period, the site was situated along tributaries of the Tagus estuary, and isotopic and zooarchaeological analysis at Cabeço da Arruda and surrounding sites suggests consumption of a mixed marine and terrestrial diet[66].

- **CCA002 (42)** is an individual of unknown age and sex dating to the Mesolithic period.

##### 3.36 Cueva de las Lechuzas

*Country:* Spain
*Region:* País Valencià, Alicante
*Coordinates:* 38.6303°, -0.863°
*Site Occupation:* c. 3300-2300 BCE (archaeological context)
*Sample Date (CLL005):* c. 3300-2300 BCE (archaeological context)
*Excavation Details:* Excavated by J. M. Soler
*Human Genetic Analysis:* Villalba-Mouco *et al.* 2021 (ref. [68])

*Summary:*

Located in a cave near Alicante in País Valencià in Southeastern Iberia, the site of Cueva de las Lechuzas consists of a Late Neolithic/Chalcolithic collective necropolis containing the remains of at least 18 individuals[68]. The site was discovered during quarrying on the small hill of Cabezo de las Cuevas. Excavations led by J. M. Soler recovered human remains and a rich assemblage of grave goods characteristic of the Southeastern Iberia Late Neolithic/Chalcolithic period[68].

- **CLL006 (Lech 8)** is an adult genetically female individual contextually dated to the Late Neolithic/Chalcolithic period 3300-2300 BCE.

##### 3.37 El Hundido

*Country:* Spain
*Region:* Burgos, Castilla y León
*Coordinates:* 42.4192°, -3.4847°
*Site Occupation:* 2500-2000 BCE, Chalcolithic and Early Bronze Age
*Radiocarbon Date (EHU002, bone, CSIC-1896):* 3933 ± 32 BP, 2564-2299 calBCE (2-sigma)[63]
*Excavation Details:* Excavations in 2005 led by Carmen Alonso under the auspices of Cronos SC Arqueología y Patrimonio[63].
*Human Genetic Analysis:* Olalde *et al.* 2019 (ref. [69]), mitochondrial data in Szécsényi-Nagy *et al.* 2017 (ref. [63])

*Summary:*

Situated in a corridor connecting the Meseta and the Valle del Ebro, the collective burial of El Hundido contains the remains of nearly 100 individuals[63]. The majority of interments date to the Late Neolithic/Chalcolithic period, after which the tomb was deliberately burned. Following a hiatus of c. 500 years, three new stone graves were constructed during the Bell Beaker period[63]. A detailed description of the site is given in ref. [63].

- **EHU002 (UE450, Hund2)** is a Late Chalcolithic adult male individual.

##### 3.38 Els Trocs Cave

*Country:* Spain
*Region:* Huesca, Aragón
*Coordinates:* 42.4964°, 0.5070°
*Site Occupation:* 5200-3700 BCE
*Radiocarbon Date (Troc1, MAMS-16159):* 6280 ± 25 BP, 5312-5212 calBCE (2-sigma)[70]
*Excavation Details:* Excavations in 2009-2012, 2014, and 2016 led by Manuel A. Rojo Guerra (Valladolid University) and José I. Royo Guillén (Aragon Government). The excavations were conducted in association with their research project "Pathways of the Neolithic" (HAR2009-09027)[63,71].
*Human Genetic Analysis:* Haak *et al.* 2015 (ref. [38]), Mathieson *et al.* 2015 (ref. [39]), mitochondrial data in Szécsényi-Nagy *et al.* 2017 (ref. [63])

*Summary:*

The site of Els Trocs is situated near the modern town of San Feliú de Veri, Bisaurri, in Aragón, Spain. Located in the Central Iberian Pyrenees, the cave sits on the southern side of Mount Els Trocs at an altitude of 1564 meters above sea level (masl)[72]. Four occupation layers attest to intermittent use of the cave from the Early Neolithic through the Roman period[63]. Remains analyzed in this study derive from Trocs I, an Early Neolithic occupation layer characterized by a broken ceramic floor and extensive deposition of faunal remains. Based on zooarchaeological, isotopic, and lithic analyses, researchers have hypothesized that agropastoralists may have used Els Trocs as an upland site for the practice of pastoral transhumance[63,72].

Disarticulated remains from 5 adults and 4 juveniles were deposited simultaneously at Trocs I and display evidence of peri- and post-mortem trauma[73]. One individual from this context was included in the present study:

- **XXX001.A16575 (Troc1, I0409)** is a genetically female juvenile of approximately 5-6 years old. This individual falls within Phase 1 of the site chronology, associated with the Early Neolithic period. Troc1 displays evidence of peri- and post-mortem blunt force trauma to the cranium[73].

#### 3.39 Es Forat de ses Aritges

*Country:* Spain
*Region:* Ciutadella de Menorca, Illes Balears
*Coordinates:* 40.0049°, 3.8616°
*Site Occupation:* c. 1400-1000/800 BCE, radiocarbon analysis of human skeletal elements
*Sample Date (EFA011):* 1200-1000 BCE, Late Bronze Age, archaeological context
*Excavation Details:* Excavated in 1995 by the ASOME research group under the auspices of the Universitat Autònoma de Barcelona.
*Human Genetic Analysis:* Villalba-Mouco *et al.* 2021 (ref. [68])

*Summary:*

Located in southwestern Menorca, the c. 14 $m^2$ collective burial of Es Forat de ses Aritges consists of a natural cave sealed with a stone wall[63]. The remains of c. 100 individuals were recovered from the cave, although most were disarticulated. Radiocarbon dating supports the repeated use of the collective burial over several hundred years, from c. 1400-1000/800 BCE[63,68]. Aside from skeletal material, the excavations recovered a variety of grave goods including pottery, copper and tin artifacts, and objects fashioned from bones and teeth[63].

- **EFA011 (FA7-1961)** is an adult female individual interred in a collective burial.

#### 3.40 La Almoloya

*Country:* Spain
*Region:* Pliego, Murcia
*Coordinates:* 37.9528°, -1.5082°
*Site Occupation:* 2200-1550 BCE, Bronze Age, El Argar period
*Sample Date (ALM029):* 1750-1550 BCE, archaeological context

*Excavation Details:* Excavated from 2013 to today by the ASOME research group under the auspices of the Universitat Autònoma de Barcelona.
*Human Genetic Analysis:* Villalba-Mouco *et al.* 2021 (ref. [68])

*Summary:*

Situated on a plateau (585 masl) in the northern foothills of Sierra Espuña, the site of La Almoloya is a large c. 3100 $m^2$ settlement associated with the El Argar period[68]. Radiocarbon analysis suggests that the site's occupation spanned all three phases of El Argar, from c. 2200-1550 calBCE. The site appears to have evolved from a frontier settlement into an urban and palatial center by the final phase (c. 1750-1550 calBCE)[68].

- **ALM029 (AY-104)** is an adult genetically female individual associated with the final phase of El Argar.

#### 3.41 La Mina

*Country:* Spain
*Region:* Soria, Castilla y León
*Coordinates:* 41.2521°, -2.5245°
*Site Occupation:* 3900-3600 BCE
*Sample Date:* 3900-3600 BCE, Neolithic, archaeological context
*Excavation Details:* Excavations in 2010 and 2012. Samples for aDNA analysis were collected on site with gloves and facemask and immediately stored at cool temperatures[38].
*Human Genetic Analysis:* Haak *et al.* 2015 (ref. [38]), Mathieson *et al.* 2015 (ref. [39])

*Summary:*

The site of La Mina includes a classic Neolithic passage grave with a >5 m corridor oriented in a south-southeast direction[38]. Radiocarbon analysis dates the structure to the 4$^{th}$ millennium BCE, a period corresponding to the beginning of Megalithism in the inner Iberian Peninsula[38]. Prior to closure, the tomb was modified to increase the mound diameter and height, and a menhir and wall of orthostats were added. This expansion has been interpreted as evidence that the site served a ceremonial function[38].

- **I0405 (Mina3)** is a genetically male individual from the Neolithic period.

#### 3.42 Lorca Sites

*Country:* Spain
*Region:* Murcia
*Coordinates:* 37.67°, -1.70°
*Site Occupation:* 2200-1550 BCE, Bronze Age, El Argar period

*Sample Date (CDL001):* 2000-1550 BCE, archaeological context
*Radiocarbon Date (LOT001, bone, OxA-7667):* 3560 ± 35 BP, 2021-1773 calBCE (2 sigma)[68]
*Radiocarbon Date (MMI002, rib fragments, OxA-7672):* 3510 ± 40 BP, 1943-1699 calBCE (2 sigma)[68]
*Sample Date (ZAP001):* 1750-1550 BCE, archaeological context
*Excavation Details:* Rescue excavations over the last 30 years.
*Human Genetic Analysis:* Samples published in Villalba-Mouco *et al.* 2021 (ref. [68]), although only LOT001 yielded sufficient data for population genetic analysis.

*Summary:*

Rescue excavations conducted over the last three decades have recovered scattered burials throughout the modern town of Lorca in Murcia province, Spain. The individuals analyzed here were recovered from burials near **Castillo de Lorca** and **Madres Mercedarias Iglesia**, as well as beneath **Los Tintes** and **Zapatería 11** streets[68]. All four contexts date to the Argaric period (2200-1550 calBCE). The scattered excavations have uncovered residential and productive areas as well as more than 50 tombs; if these constitute one continuous settlement, Lorca may have been one of the largest urban areas of the El Argar Culture[63].

- **CDL001 (Castillo de Lorca Loct4)** is a 40–50-year-old female individual contextually dated to the second or third El Argar phase.
- **LOT001 (Los Tintes 2-1)** is a 35-45-year-old genetically male individual[68].
- **MMI002 (Madres Mercedarias Iglesia Tomb 11-1)** is a 28-35-year-old female individual.
- **ZAP001 (Lorca Zapatería Urna 3 Tumba 3)** is a mature adult male contextually dated to the third phase of El Argar.

##### 3.43 Paris Street

*Country:* Spain
*Region:* Cerdanyola, Barcelona
*Coordinates:* 41.4919°, 2.1389°
*Site Occupation:* 2850-2250 BCE
*Sample Date (I1553 and I0220):* 2850-2250 BCE (radiocarbon-based estimates for the hypogeum’s use period)
*Excavation Details:* Excavations initiated following discovery of the burial during construction work in 2003.
*Human Genetic Analysis:* Individual I01553 published in Olalde *et al.* 2018 (ref. [64]).

*Summary:*

The site of Paris Street consists of a large hypogeum located in the town of Cerdanyola del Vallés in Barcelona province, Spain. In addition to c. 9000 human remains, the hypogeum contained lithics and ceramics characteristic of the Bell Beaker culture[64]. Radiocarbon-based estimates for the hypogeum's use period range from 2850-2250 BCE, providing a contextual age for the samples included in this study[64].

- **XXX001.A161345 (I1553)** is a genetically female individual associated with the Chalcolithic Bell Beaker culture.
- **XXX001.A161274 (I0220)** is an individual of unknown sex whose low endogenous DNA content precluded human genetic analyses.

##### 3.44 Panoría

*Country:* Spain
*Region:* Granada (Darro), Andalucía
*Coordinates:* 37.3504°, -3.2934°
*Site Occupation:* 3500-2000 BCE
*Sample Date: 3500–3300 cal BC (Based on the 17 radiocarbon dates of Phase B of Tomb 3)*
*Excavation Details:* Excavated in 2015 and 2019 by the University of Granada

*Summary:*

The megalithic cemetery of Panoría is located in the southeast of the Iberian Peninsula. This site was discovered in 2012 and consists of at least 19 tombs with polygonal, rectangular or trapezoidal funerary chambers and short corridors. In 2015 and 2019, nine tombs were excavated by the GEA research group from the University of Granada. Four of them –Tombs 3, 10, 11 and 15– contained largely undisturbed ritual deposits. To better understand the multi-depositional mortuary practices, special attention has been paid to their chronology. A radiocarbon series of 73 dates was obtained, mainly from the four well-preserved tombs. All dated samples were from human remains. According to statistical models, the Panoría cemetery presents a punctuated pattern of use with four peaks of ritual intensity dated in the 34th, 29th, 25th and 21st centuries calBCE.

- **PAN049 (DA3135_77)** is a mature adult (41-60 years old) identified osteologically as probably female, found in Tomb 3 (Phase B), contextually dated to the Late Neolithic period.

#### Northern Europe

##### 3.45 Linton

*Country:* England
*Region:* Cambridgeshire
*Coordinates:* 52.0991°, 0.277°
*Site Occupation:* Early- middle Saxon period, 5th-8th century CE

*Sample Date: Middle Saxon, later 7th-8th century*
*Radiocarbon Date (Sk351, SUERC-20250):* 1205 ± 30 BP, 635-937 calCE (2 sigma, IntCal13)
*Excavation Details:* Excavations between 2004 and 2010 led by Oxford Archaeology East, with funding from the Cambridge City Council.
*Human Genetic Analysis:* Gretzinger *et al.* 2022 (ref. [74]), other individuals from Linton (excluding SK351) published in Schiffels *et al.* 2016 (ref. [75])

*Summary:*

Located in Cambridgeshire in the valley of the River Granta, the c. 8 ha site of Linton preserves archaeological contexts dating to the Late Neolithic, Middle to Late Bronze Age, Iron Age, and Roman, Early, and Middle Saxon periods, as well as showing post-Medieval activity[75]. The individual analyzed here was interred in one of three Anglo-Saxon era graves, containing five people who were situated adjacent to a former Roman trackway. The roughly rectangular grave was aligned to the northeast and southwest and contained the remains of three individuals Sk351, as well as sk350 and sk352[75].

- **XXX001.A161161 (Sk351, I0791)** is an adult genetically female individual, likely over the age of 45 years, from a triple grave. Sk351 was decapitated prior to burial and interred over, but within the same grave as two children of c. 12 and 5 years old. The postcranial skeleton was deposited in an extended, supine position with the right femur overlying the skull. In addition to perimortem trauma of the fourth and fifth cervical vertebrae associated with decapitation, the individual exhibited developmental anomalies, maxillary sinusitis, Schmorl's nodes, and joint disease[75].

##### 3.46 Oakington

*Country:* England
*Region:* Cambridgeshire
*Coordinates:* 52.2614°, 0.0687°
*Site Occupation:* c. 400-600 CE
*Sample Date:* c. 400-600 CE
*Excavation Details:* First identified in 1926, the cemetery at Oakington was rediscovered in 1993 during the construction of a children's playground. Excavations in 1994 led by the Cambridge County Council's Archaeological Field Unit recovered the remains of 24 individuals, which were subsequently reinterred around the millenum. Further excavations by the same organization in 2006-2007 unearthed skeletons from 17 more individuals. Systematic excavation of the site was carried out between 2010 and 2015 under the auspices of the University of Central Lancashire and Oxford Archaeology East, yielding remains from an additional 128 individuals. Grave 57 was excavated in 2011.
*Human Genetic Analysis:* Gretzinger *et al.* 2022 (ref. [74]), other individuals from Oakington (excluding Oakington1375) published in Schiffels *et al.* 2016 (ref. [75]).

*Summary:*

Situated in Cambridgeshire c. 7 km northwest of Cambridge, the Anglo-Saxon cemetery of Oakington contains a large number of burials mostly dating between the 5th and 6th centuries CE[75]. Although artifact conservation and investigation of the skeletal assemblage is ongoing, preliminary findings suggest an overrepresentation of young individuals in the burials, possibly consistent with the site serving a central role in a regional kinship network[76].

- **XXX001.A161133 (Oakington1375, Grave 57a, I0763)** is an adult female c. 25-30 years of age. The individual was interred in supine position with a fetus in transverse position across the pelvic cavity, suggesting childbirth complications as a likely cause of death. Grave goods included three brooches, one of which was cruciform, amber and glass beads, wrist clasts, belt fittings, and a knife[74,75].

#### 3.47 Renko

*Country:* Finland
*Region:* Tavastia proper
*Coordinates:* 60.8936°, 24.2870°
*Site Occupation:* 1500-1800 CE
*Sample Date:* ~1770CE
*Radiocarbon Date (REN010, phalanx, HELA-4169):* 175 ± 31 BP
*Excavation Details:* Excavated in 1984.
*Human Genetic Analysis:* Mitochondrial data from individuals from Renko published in Översti *et al.* 2019 (ref. [77])

*Summary:*

The stone Church of St. Jacob in Renko was constructed in the early 16th century, although it has been hypothesized that a wooden church stood on the site as early as the turn of the 15th century. The stone structure was renovated and reopened in the late 18th c. after being abandoned in the mid-17th century. Excavations in 1984 revealed ~70 burials inside the church. Only part of these were excavated due to the location of many graves under the building's structural walls, and the human remains were not systematically studied or stored for research. The archaeological survey and excavation of the yard outside the church was conducted by K. Salo in 2008-2009 during the drainage renovations of the church, including an osteological analysis, with notes on pathologies where applicable.

- **REN010 (JK1939; Grave 19, individual 58)** is an adult genetically male individual of approximately 40-50 years of age, with pathological changes in their maxilla, nasal bone, and phalanges, possibly indicative of an infectious disease. The individual had lost 11 teeth during life. They also had visible parodontitis, caries in on one tooth, and a severe abscess the width of four teeth, reaching into the jawbone.

Siberia

#### 3.48 Khaptsagai

*Country:* Russian Federation
*Region:* Upper Lena River, Siberia
*Coordinates:* 53.0445°, 105.5106°
*Site Occupation:* Glazkovo stage of the Late Neolithic, archaeological context
*Sample Date: 2200-1200 BCE*
*Radiocarbon Date (KPT003, tooth, MAMS-37927):* 4021 ± 19 BP, 2576-2480 calBCE (2-sigma, IntCal13). Calibrated date as published in Yu *et al.* 2020, incorporating a 385 year offset due to the reservoir effect.
*Excavation Details:* Excavations in 1927 as part of an archaeological expedition in the Kachug region.
*Human Genetic Analysis:* Yu *et al.* 2020 (ref. [78])

*Summary:*

Situated in the area of the Upper Lena River on the banks of the Manzurka River, Khaptsagai consists of both Neolithic dwelling sites and a burial ground containing at least twenty graves[78]. Along with the human remains, excavations recovered numerous grave goods including nephrite rings and knives, stone arrowheads and spear points, chisels, awls, and stone needles in an ornamented case[78]. The Neolithic individuals from Khaptsagai relied on a hunter-gatherer subsistence strategy; consistent with other studies of individuals from the cis-Baikal region, stable isotope analysis of human remains from Khaptsagai suggests a reliance on $C_3$ plant resources with a mix of terrestrial and freshwater aquatic protein sources[78,79].

- **KPT003 (8302)** is a genetically male individual.

### IV. OCEANIA

#### 4.1 Futuna, Vanuatu

*Country:* Vanuatu
*Region:* Futuna
*Coordinates:* -19.5116°, 170.23°
*Site Occupation:* 680-980 CE (radiocarbon analysis of five individuals from rockshelters FuRS1A and FuRS12)
*Radiocarbon Date (FUT008, tooth, MAMS-29689):* 1376 ± 29 BP, 603-759 calCE (2-sigma)[80]
*Excavation Details:* Excavations on Futuna were undertaken by Richard and Mary Shutler in 1963-1964 as part of a larger initiative by the Pacific Science Congress of 1961 with support from the Bishop Museum. Following a 50-year hiatus, excavations under the auspices of the

Australian Research Council (DP160103578) and the French Ministry of Foreign Affairs (MEAE, commission des fouilles) were conducted up to 2020.
*Human Genetic Analysis:* Posth *et al.* 2018 (ref. [80])

*Summary:*

The remains analyzed here were recovered from one of the many rockshelters on Futuna, a raised coral island in Vanuatu measuring approximately 5 km long and reaching its highest point at 666 masl. Located in the island's northeastern Ipau district, the c. 13.7 x 3.6 m rockshelter FuRS12 contained the remains of at least 15 individuals, all of which were dated to between 970-1270 years BP. Situated towards the back of the rockshelter close to the bedrock, many of the burials in FuRS12 were lined or covered with rocks and included grave goods.

- **FUT008 (FURS12 burial 12)** is an adult of unknown sex.

##### 4.2 Talasiu, Tongatapu, Tonga

*Country:* Tonga
*Region:* Tongatapu
*Coordinates:* -21.17°, -175.11°
*Site Occupation:* 750-650 BCE, Late Lapita/Post-Lapita Period (radiocarbon and archaeological analysis)
*Sample Date:* 750-650 BCE
*Excavation Details:* Excavations in 2008 recovered the burned/partially burned remains of four individuals which had begun to erode out of a road cut. Additional human remains have been unearthed since 2013 during excavations led by Frederique Valentin and Geoffrey Clark. These operations were supported by the Ministry of Internal Affairs (Kingdom of Tonga) with funding from the French Government (MEAE, Commission des fouilles à l'étranger) and Australian Research Council grant (DP200102872)
*Human Genetic Analysis:* Individuals from Talasiu (excluding TON003) are published in Posth *et al.* 2018 (ref. [80]) and Skoglund *et al.* 2017 (ref. [2]). Since that time, on-going excavations have been conducted at the site which increased the number of burials.

*Summary:*

Located on the coastline of Tongatapu near the Fanga 'Uta Lagoon, the site of Talasiu consists of a c. 90 cm thick shell midden spanning approximately 450 m. A number of burials containing one or more individuals were excavated at the site. Radiocarbon analyses of skeletal materials, paleobotanical remains, charcoal, and worked shell grave goods date the deposition of midden and burials to between 750-650 BCE,.

- **TON003 (Sk9.3)** is an adult individual of indeterminate sex[81].

#### 4.3 Tilu

*Country:* Papua New Guinea
*Region:* Madang Province
*Coordinates:* -5.1257°, 145.8014°
*Site Occupation:* c. 1300-1400 CE (Pre-colonial Madang Period)
*Sample Date:* Nägele *et al.* 2025 (ref. [82]) directly dated another human specimen (TIL001 [original lab code T-1702]) to c.1260–1450 CE, which overlaps with charcoal dates from the same excavation layer (c.1300–1400 CE). Because TIL001 comes from Spit 4, just below TIL003 [T-1701] in Spit 3, it is highly likely that these specimens are very similar in date (they are not from the same individual).
*Excavation Details:* Excavations in 2014 were undertaken by the University of Otago and the National Museum and Art Gallery of Papua New Guinea, alongside members of the local Malmal community. This site had previously been excavated by Brian Egloff in the 1970s and the 2014 research sought to clarify the sequence and timing of occupation as it related to Bel language dispersals and pottery manufacture in the Madang area. Occupation at the site dates to c. 1300-1400 CE based on radiocarbon dating of one human tooth and six wood charcoal specimens. A single wood charcoal radiocarbon date reported in Gaffney *et al.* 2018 (ref. [83]) might suggest occupation as early as 1050-1150 CE, but this is unconfirmed.
*Human Genetic Analysis:* Individuals (TIL001–TIL004) from Tiluare described in Nägele *et al.* 2025 (ref. [82]). However, TIL003 did not yield enough DNA and was deemed to have poor preservation, therefore being excluded from their analysis.

*Summary:*

Located roughly 12 km north of Madang township on the northeast coast of Papua New Guinea, the clan area of Tilu at Malmal village includes two mounds of roughly 20 m long. In addition to a small number of loose fragmentary human remains, pottery, obsidian, shell and animal bones were recovered from the site[83]. Material culture analysis demonstrates recent occupation during the Pre-colonial Madang Period, a phase of Post Lapita pottery manufacture immediately prior to the ethnographic record.

- **TIL003 (original specimen code T-1701)** is a right mandibular fragment. M2 and M3 are in situ and display no macroscopic wear, although there is a slight calculus build up and periodontal erosion.

#### S2. Supplementary Notes

##### Supplementary Note 1: Strain Heterozygosity

As bacteria like *S. mutans* are haploid, the frequency and distribution of multiallelic sites provides a metric for exploring the presence of strain mixture and/or background contamination in metagenomic datasets[84]. To identify ancient individuals likely infected with more than one strain of *S. mutans*, we performed heterozygous SNP calling on all high-coverage ancient samples, here defined as those having a mean coverage of the UA159 reference genome greater than 10.0x. For each sample, we computed the proportion of SNPs classified as heterozygous by MultiVCFAnalyzer and constructed a histogram visualizing counts of SNP positions by derived allele frequency (**Supplementary Figure 1)**.

For a monomorphic *S. mutans* infection, the vast majority of loci should be homozygous. Nevertheless, a small number of heterozygous loci are expected due to errors in SNP calling caused by ancient DNA damage and/or mismapping from environmental taxa; therefore, we consider samples with less than 10% of heterozygous SNP positions to harbor single *S. mutans* strains. Out of 35 high coverage ancient *S. mutans* samples, 19 (54%) have fewer than 10% heterozygous SNP positions, indicating the presence of a monomorphic bacterial infection. For example, out of 10,902 SNP calls in the *S. mutans* genome reconstructed from individual VLI009, only 51 (0.5%) are classified as heterozygous; furthermore, these multiallelic positions are distributed evenly across the reference genome, suggesting random noise caused by mismapping from environmental sources (**Supplementary Figure 1**).

A high proportion of heterozygous SNP calls may be due to (a.) mixture of multiple strains and/or closely related species or (b.) background contamination. Visualization of derived allele frequencies across genomic positions can help to differentiate between these two hypotheses and provide clues as to the proportion of strain mixtures. Individuals infected with multiple *S. mutans* strains should exhibit a large number of multiallelic loci distributed evenly across the reference genome. For a mixture of two strains in equal abundance (symmetric), we expect heterozygous positions to form a unimodal distribution with a derived allele frequency of approximately 50%. The *S. mutans* genome reconstructed from individual REV004, for example, exhibits 47.5% heterozygous SNPs, and derived allele frequencies at these positions form a unimodal distribution (**Supplementary Figure 1**). Strain mixtures in uneven abundance (asymmetric) should exhibit a multimodal distribution with derived allele frequency peaks reflecting the relative proportion of the major and minor strain. For example, 22.7% of SNP positions in the DON004 genome are classified as heterozygous; these positions form a bimodal distribution with derived allele frequencies peaking around c. 20% and c. 80%, reflecting a likely case of asymmetric strain mixture (**Supplementary Figure 1**).

Beyond strain mixture, a high frequency of heterozygous SNP positions may reflect background contamination from closely related taxa. Background contamination may present as a large number of heterozygous positions with low derived allele frequencies, which may not exhibit distributions characteristic of symmetric and/or asymmetric strain mixtures. We predict that multiallelic positions deriving from background contamination may also exhibit an uneven distribution across the *S. mutans* reference genome. *S. mutans* is known to exchange genomic regions with other species in the oral cavity; recently transferred loci may create “heterozygosity hotspots” particularly prone to mismapping in metagenomic contexts. However, we expect that MALT filtering may significantly reduce the frequency of multiallelic sites in these strains by removing reads that map more closely to non-*mutans* taxa. As an example, the genome reconstructed from KIV007 exhibited a frequency of multiallelic sites of 14.5% prior to MALT filtering. Notably, these multiallelic positions fell in distinct bands across the *S. mutans* genome, suggesting a possible case of mismapping. Consistent with this hypothesis, the MALT filtering step reduced the number of multiallelic sites from 173 to 13, resulting in a final heterozygous site frequency of 1.9% (KIV007). Thus, in order to reduce the impact of background contamination, we utilized the MALT-filtered dataset for all subsequent analysis.

#### Sources Cited

1. Manhire, A. A Report on the Excavations at Faraoskop Rock Shelter in the Graafwater District of the South-Western Cape. *Southern African Field Archaeology* **2**, 3–23 (1993).

2. Skoglund, P. *et al.* Reconstructing Prehistoric African Population Structure. *Cell* **171**, 59–71 (2017).

3. Sealy, J. C., Patrick, M. K., Morris, A. G. & Alder, D. Diet and dental caries among Later Stone Age inhabitants of the Cape Province, South Africa. *Am. J. Phys. Anthropol.* **88**, 123–134 (1992).

4. Andrades Valtueña, A. Beyond phylogenies: advancing analytical approaches for the field of ancient pathogenomics. (Friedrich Schiller University Jena, 2021).

11. Ávila López, R. *Excavaciones Arqueológicas En San Gregorio Atlapulco, Xochimilco, México*. (Subdirección de Salvamento Arqueológico, Instituto Nacional de Antropología e Historia (INAH), México, 1995).

12. Hernández Espinoza, P. O. Entre flores y chinampas: La salud de los antiguos habitantes de Xochimilco. in *Sociedad y Salud en el México Prehispánico y Colonial* (ed. Márquez Morfín, L. & Hernández Espinoza, P) 327–366 (CONACULTAINAH/Promep, 2006).

13. Hernández Espinoza, P. O. *La Regulación Del Crecimiento de La Población En El México Prehispánico*. ((Instituto Nacional de Antropología e Historia (INAH), 2006).

22. Irvine, B. T. An Isotopic Analysis of Dietary Habits in Early Bronze Age Anatolia. (Freie Universität Berlin, 2017).

29. Oinonen, M., Vasks, A., Zarina, G. & Lavento, M. Stones, bones, and hillfort: Radiocarbon

dating of ķivutkalns bronze-working center. *Radiocarbon* **55**, 1252–1264 (2013).

35. Магомедов, Р. Г. *Материалы к изучению культур эпохи бронзы в Приморском Дагестане*. (Махачкала, 2000).

42. Stecher, M., Grünberg, J. M. & Alt, K. W. Bioarchaeology of the Mesolithic individuals from Bottendorf (Thuringia, Germany). in *Mesolithic Burials- Rites, Symbols and Social Organisation of Early Postglacial Communities* (eds. Grünberg, J. M., Gramsch, B., Larsson, L., Orschiedt, J. & Meller, H.) (International Conference Halle (Saale), Germany, 18th-21st September 2013.).

43. Leinthaler, B., Bogen, C. & Döhle, H.-J. Von Muschelknöpfen und Hundezähnen – Schnurkeramische Bestattungen bei Esperstedt. in *Archäologie auf der Überholspur* –

*Ausgrabungen an der A38, Archäologie in Sachsen-Anhalt, Sonderband 5* (ed. Meller, H.) 59–82 (Halle, 2006).

51. Dobeš, M. & Limburský, P. *Pohřebiště Staršího Eneolitu a šňůrové Keramiky ve Vliněvsi. S Příspěvky Želimira Brnića, Jakuba Likovského, Miroslava Popelky, René Kyselého a Jaroslava Hlaváče – Gräberfeld Des älteren Äneolithikums Und Der Schnurkeramik in Vliněves. Mit Beiträgen von Želimir Brnić, Jakub Likovský, Miroslav Popelka, René Kyselý Und Jaroslav Hlaváč. Archeologické Studijní Materiály 22*. (2013).

52. Limburský, P. *Pohřebiště Kultury Se Zvoncovitými Poháry ve Vlíněvsi. K Problematice a Chronologii Konce Eneolitu a Počátku Doby Bronzové – The Bell Beaker Cemetery in Vlíněves. Dissertationes Archaeologicae Brunenses/Pragensesque 13*. (Prague, 2012).

53. Limburský, P. *et al. Pohřební Areály únětické Kultury ve Vliněvsi – Burial Areas of the Únětice Culture in Vliněves*. (Praha, 2018).

54. Hallager, B. P. & MacGeorge, P. J. Late Minoan III burials at Khania: the tombs, finds, and deceased in Odos Palama. in *Studies in Mediterranean archaeology* (Åströms, 1992).

55. Skourtanioti, E. *et al.* Ancient DNA reveals admixture history and endogamy in the

prehistoric Aegean. *Nat Ecol Evol* **7**, 290–303 (2023).

67. Peyroteo Stjerna, R. On Death in the Mesolithic: Or the Mortuary Practices of the Last Hunter-Gatherers of the South-Western Iberian Peninsula, 7th–6th Millennium BCE. (Uppsala University, 2016).

68. Villalba-Mouco, V. *et al.* Genomic transformation and social organization during the Copper Age–Bronze Age transition in southern Iberia. *Sci. Adv.* **7**, (2021).

69. Olalde, I. *et al.* The genomic history of the Iberian Peninsula over the past 8000 years. *Science* **363**, 1230–1234 (2019).

70. Rojo-Guerra, M. *et al.* Pastores trashumantes del Neolítico antiguo en un entorno de alta montaña: secuencia crono-cultural de la Cova de Els Trocs (San Feliú de Veri, Huesca).

*Universidad de Valladolid* http://hdl.handle.net/10261/146650 (2013).

76. Sayer, D. *Early Anglo-Saxon Cemeteries: Kinship, Community and Identity*. (Manchester University Press, Manchester, England, 2020).
